# Pan-cancer survey of tumour mass dormancy and underlying mutational processes

**DOI:** 10.1101/2021.04.25.441168

**Authors:** Anna Julia Wiecek, Daniel Hadar Jacobson, Wojciech Lason, Maria Secrier

**Affiliations:** UCL Genetics Institute, Department of Genetics, Evolution and Environment, University College London, UK

**Keywords:** tumour mass dormancy, immunity, angiogenesis, mutational signatures, APOBEC, hypoxia

## Abstract

Tumour mass dormancy is the key intermediate step between immune surveillance and cancer progression, yet due to its transitory nature it has been difficult to capture and characterise. Little is understood of its prevalence across cancer types and of the mutational background that may favour such a state. While this balance is finely tuned internally by the equilibrium between cell proliferation and cell death, the main external factors contributing to tumour mass dormancy are immunological and angiogenic. To understand the genomic and cellular context in which tumour mass dormancy may develop, we comprehensively profiled signals of immune and angiogenic dormancy in 9,631 cancers from the Cancer Genome Atlas and linked them to tumour mutagenesis. We find evidence for immunological and angiogenic dormancy-like signals in 16.5% of bulk sequenced tumours, with a frequency of up to 33% in certain tissues. Mutations in the *CASP8* and *HRAS* oncogenes were positively selected in dormant tumours, suggesting an evolutionary pressure for controlling cell growth/apoptosis signals. By surveying the mutational damage patterns left in the genome by known cancer risk factors, we found that ageing-induced mutations were relatively depleted in these tumours, while patterns of smoking and defective base excision repair were linked with increased tumour mass dormancy. Furthermore, we identified a link between APOBEC mutagenesis and dormancy, which comes in conjunction with immune exhaustion and may partly depend on the expression of the angiogenesis regulator *PLG* as well as interferon and chemokine signals. Tumour mass dormancy also appeared to be impaired in hypoxic conditions in the majority of cancers. The microenvironment of dormant cancers was enriched in cytotoxic and regulatory T cells, as expected, but also in macrophages and showed a reduction in inflammatory Th17 signals. Finally, tumour mass dormancy was linked with improved patient survival outcomes. Our analysis sheds light onto the complex interplay between dormancy, exhaustion, APOBEC activity and hypoxia, and sets directions for future mechanistic explorations.

## 1 INTRODUCTION

Tumour evolution is shaped by a variety of internal and external forces that act at different stages during cancer development, sometimes in an antagonistic manner, and drive distinct trajectories to advanced disease (McGranahan and Swanton 2017; Gerlinger et al. 2014; Temko et al. 2018). Within this rapidly changing context, adaptation of cancer cells is paramount for survival and much of it is achieved through cellular plasticity (Yuan, Norgard, and Stanger 2019). As a manifestation of this plasticity, cancer dormancy has emerged as an important contributor to the early stages of tumour development, as well as to cancer progression and metastatic dissemination (Phan and Croucher 2020; Jahanban-Esfahlan et al. 2019; Park and Nam 2020). Its two facets, cellular dormancy driven by arrest in the G0 state of the cell cycle (Phan and Croucher 2020), and tumour mass dormancy (TMD), described as an equilibrium between cell proliferation and cell death shaped by the microenvironment that constrains tumour growth (Aguirre-Ghiso 2007; H.F. Wang et al. 2019; Holmgren, O’Reilly, and Folkman 1995), are complementary but distinct mechanisms that contribute to the plasticity of cancer expansion (Huang 2021; Shen and Clairambault 2020). The former concept was coined already in 1954 by Geoffrey Hadfield (HADFIELD 1954) and has since led to the generation of more detailed mechanistic insights explaining its regulation by the DREAM complex (Sadasivam and DeCaprio 2013; MacDonald et al. 2017; Kim et al. 2021), with key dependencies on the p53/p21 activation axis (Itahana et al. 2002; Heldt et al. 2018; Barr et al. 2017). The latter, initially termed “population dormancy” by Judah Folkman in 1972 (Gimbrone et al. 1972) and then renamed to tumour mass dormancy, remains poorly understood due to the lack of suitable data and models (Boire et al. 2019).

The theoretical model of TMD is embedded into the “3Es of immunoediting” paradigm (Dunn, Old, and Schreiber 2004b), arising as a temporary equilibrium between tumour elimination and immune escape (Teng et al. 2008; Koebel et al. 2007). As the cancer lesion develops, a period of immunoediting follows when the immune system interacts with the malignant cells establishing a dynamic equilibrium: the immunogenic cells are eliminated, and non-immunogenic tumour cells arise (Dunn, Old, and Schreiber 2004a). This keeps the tumour in a dormant state (H.F. Wang et al. 2019; Aguirre-Ghiso 2007), hypothesised to account for the “latency” between the initiation of the first mutator phenotype and symptom manifestation in early disease, or for the disease-free period preceding cancer recurrence (Gužvić and Klein 2013; Damen, van Rheenen, and Scheele 2020). During this period of dormancy, the continued cytotoxic response triggered by pro-inflammatory signalling cytokines like interferon ɣ, as well as prolonged exposure to pro-inflammatory signalling in general, causes the cytotoxic cells to become inactive, a phenomenon known as T cell exhaustion (Dunn, Koebel, and Schreiber 2006; Wherry and Kurachi 2015; Yi, Cox, and Zajac 2010). Finally, the angiogenic switch or immune escape shifts the balance in favour of cancer progression (Jahanban-Esfahlan et al. 2019).

A variety of molecular mechanisms have been proposed to mediate these switches. The urokinase receptor (uPAR) regulates tumour growth by controlling β1 integrin signalling which drives a cascade of Ras/ERK mitogenic activation via the focal adhesion kinase (FAK) and the EGF receptor (EGFR) (Aguirre Ghiso, Kovalski, and Ossowski 1999; White et al. 2004; Liu et al. 2002; Aguirre Ghiso 2002). Downregulation of any of these components has been shown to lead to tumour growth arrest and dormancy (Aguirre-Ghiso 2007). Additionally, blocking uPAR activates p38, which induces dormancy under p53 upregulation and downregulation of JUN (Ranganathan et al. 2006). Metastatic lesions appear to manifest reduced p38 activity under sustained ERK activation. As a result, a low ERK:p38 protein expression ratio is often employed to assess tumour dormancy (Aguirre-Ghiso et al. 2003). This state is further corroborated by a limited ability in recruiting new blood vessels and vasculature remodelling, resulting in angiogenic dormancy (Moserle, Amadori, and Indraccolo 2009). This condition is often characterised by VEGF inhibition in the presence of anti-angiogenic factors such as angiostatin, endostatin, thrombospondin, etc., or chemokines (CXCL9, CXCL10) (Senft and Ronai 2016; Lyu et al. 2013; Aguirre-Ghiso 2007; Park and Nam 2020).

Overall, it is clear that TMD results from an interplay between immunological and angiogenic dormancy, where cell growth is counterbalanced by apoptosis due to poor vascularisation and immune cytotoxicity, followed by exhaustion. It becomes evident that the states of T cell exhaustion and dormancy are not a simple dysfunction, but a purposeful homeostatic mechanism enabling the prevention of pathological immune responses. Hence, it is of crucial importance to be able to identify and target the dormant tumours early. This temporary equilibrium state provides a unique clinical opportunity, but its prevalence in cancer and the genetic determinants that may sustain it are currently unknown.

In this study, we surveyed the landscape of TMD along the two axes that shape it, immunological and angiogenic dormancy, in a cohort of 9,631 tumours from 31 tissues available from TCGA. We show that TMD can be captured from bulk sequencing datasets, that it is pervasive across a variety of cancer types and that its emergence is linked with key somatic alterations. We also investigate the environmental context of TMD and its relevance in the clinic.

## 2 RESULTS

### 2.1 Pan-cancer characterisation of immunological, angiogenic and tumour mass dormancy

To evaluate the levels of immunological and angiogenic dormancy across multiple cancer tissues, we manually curated lists of genes associated with the two programmes from the literature (Supplementary Table 1). The lists included receptor molecules, as well as soluble mediators, surface and structural proteins, enzymes and transcription factors, several of which are observed to be frequently mutated during cancer development (Supplementary Figure 1). Immunological and angiogenic dormancy programme scores were assigned on a per-sample basis using two distinct methodologies that exploited the expression of genes associated with either process, focusing either on differences between up/downregulated gene activity (‘scaled difference of means’) or on the largest variation explained by principal component analysis (‘PCA’) (see Materials and Methods, Supplementary Figures 2–4). Both immunological and angiogenic dormancy represent potential mechanisms for restricting the expansion of primary tumour cell populations due to either impaired vascularization or immunosurveillance (Aguirre-Ghiso 2007), therefore both processes can contribute to the development of TMD. As such, an overall per-patient TMD programme score was also derived using the expression of genes associated with both types of dormancy. We assessed the robustness of the two scoring methods to changes in the gene signature employed or small variations in gene expression, and showed that the ‘scaled difference of means’ approach was comparatively more stable to such fluctuations (Supplementary Figure 5). Therefore, we chose this method for the downstream analysis.

Having established a framework for quantifying immunological, angiogenic and tumour mass dormancy programmes, we next profiled these cellular programmes across 9,631 samples of solid primary tumours from the Cancer Genome Atlas (TCGA). We observed a spectrum of TMD across tumours that ranges from highly dormant to highly expanding (Figure 1A-C, Supplementary Figures 6–7). TMD is thought to emerge when tumour cell proliferation is balanced by apoptosis due to factors such as limitations in blood supply or an active immune system (Holmgren, O’Reilly, and Folkman 1995). Consequently, primary tumour samples with high TMD programme scores (upper quartile of the score range) which also showed a proliferation/apoptosis ratio below 1 (see Materials and Methods), indicative of limited primary tumour lesion expansion, were classed as exhibiting TMD (Figure 1D). Overall, 16.5% of samples across different tissues exhibited substantial evidence for a TMD-like phenotype (with up to 33% prevalence in certain cancers) and they were further subdivided into those indicating angiogenic dormancy (4.4%), immunological dormancy (5%), or both (7%) (Figure 1E and Supplementary Table 2). Head and neck (32%), sarcoma (32%) and breast (26%) cancer were among the cancers with highest rates of TMD. Angiogenic dormancy alone was most widespread in breast (14%) and head and neck cancers (13%). Some rarer cancers like thymoma or pheochromocytoma and paraganglioma also showed remarkably high prevalence of TMD (~33% each) with the latter also exhibiting the highest immune-mediated dormancy (21%). The systematic differences in TMD programme scores across different tissues (Figure 1E-F) suggest that the tissue environment may impact the ability of the tumour to enter a TMD state. Across the board, immune-mediated dormancy levels appeared higher than those of angiogenic dormancy, suggesting a dominant role for immune surveillance in determining TMD.

**Figure 1.**
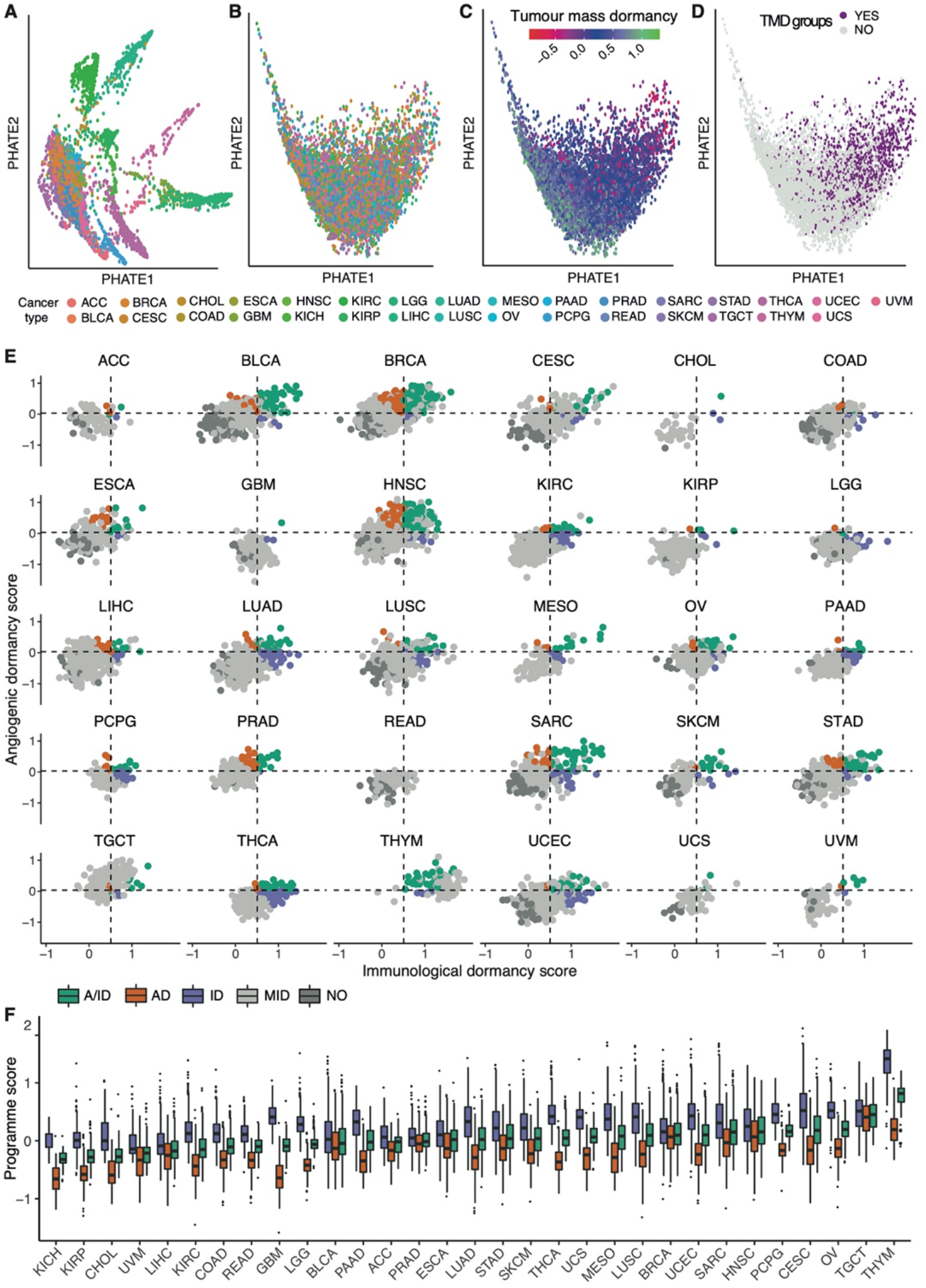
The pan-cancer landscape of TMD. **(A-D)** PHATE dimensionality reduction applied to 9,631 primary tumor samples based on the expression of genes within the TMD and exhaustion programmes before **(A)** and after **(B-D)** removal of tissue specific expression patterns. The maps are coloured by their corresponding tissue type **(A-B)**, TMD programme score **(C)** and TMD status **(D)**. **(E)** Relationship between immunological and angiogenic programme scores within individual TCGA cancer tissues. Samples are colored by their angiogenic (orange, AD), immunological (purple, ID) and tumour mass (green, A/ID) dormancy status. Samples showing no evidence of TMD (NO) are coloured in dark grey, and slowly expanding tumours in light grey (MID). Horizontal and vertical dashed lines represent the upper quartile of the pan-cancer angiogenic and immunological dormancy programme scores, respectively. KICH was not plotted because it lacked TMD samples. **(F)** Variation in TMD, immunological and angiogenic dormancy scores across primary tumor TCGA samples stratified by tissue type. The tissues are sorted by their TMD levels.

Even though TMD is expected to manifest primarily in early forming tumours (Dunn, Old, and Schreiber 2004b), we surprisingly found that a sizeable proportion of late-stage tumours also exhibited similarly high TMD-linked activity levels (Supplementary Figure 8). When comparing the prevalence of TMD between early- and late-stage cancers, expanding tumours appeared enriched in the later stages (Fisher’s exact test p < 0.0001, 1.5-fold enrichment), as expected. Remarkably, samples with angiogenic dormancy were also marginally enriched (1.3-fold) in late-stage cancers (Fisher’s exact test p < 0.05).

### 2.2 The genomic background of TMD

To gain a better understanding of the genomic context in which TMD can develop, we asked whether samples with high and low TMD differed in their association with known drivers of tumorigenesis. We found 15 genes with either a statistically significant enrichment or a depletion of mutations within samples classed as showing TMD across the 31 solid cancer tissues, with several depletion signals characterizing both early- and late-stage tumours (Fisher’s exact test adjusted p<0.05, Figure 2A, Supplementary Figures 9A, 10A, Table 3). Furthermore, mutations in *MUC4*, a gene involved in angiogenesis and metastasis (Zhi et al. 2014), were specifically enriched in stomach adenocarcinoma, while *EGFR*, *FAT3/4*, *LRP1B*, *KAT6B* mutations, mainly linked with cell proliferation and immune responses (Katoh 2012; H. Chen et al. 2019; Simó-Riudalbas et al. 2015; Sigismund, Avanzato, and Lanzetti 2018), were depleted in colon cancer (Fisher’s exact test adjusted p<0.05, Supplementary Figure 11). Many of these findings were reaffirmed using a random forest classification approach (Supplementary Figure 12).

**Figure 2.**
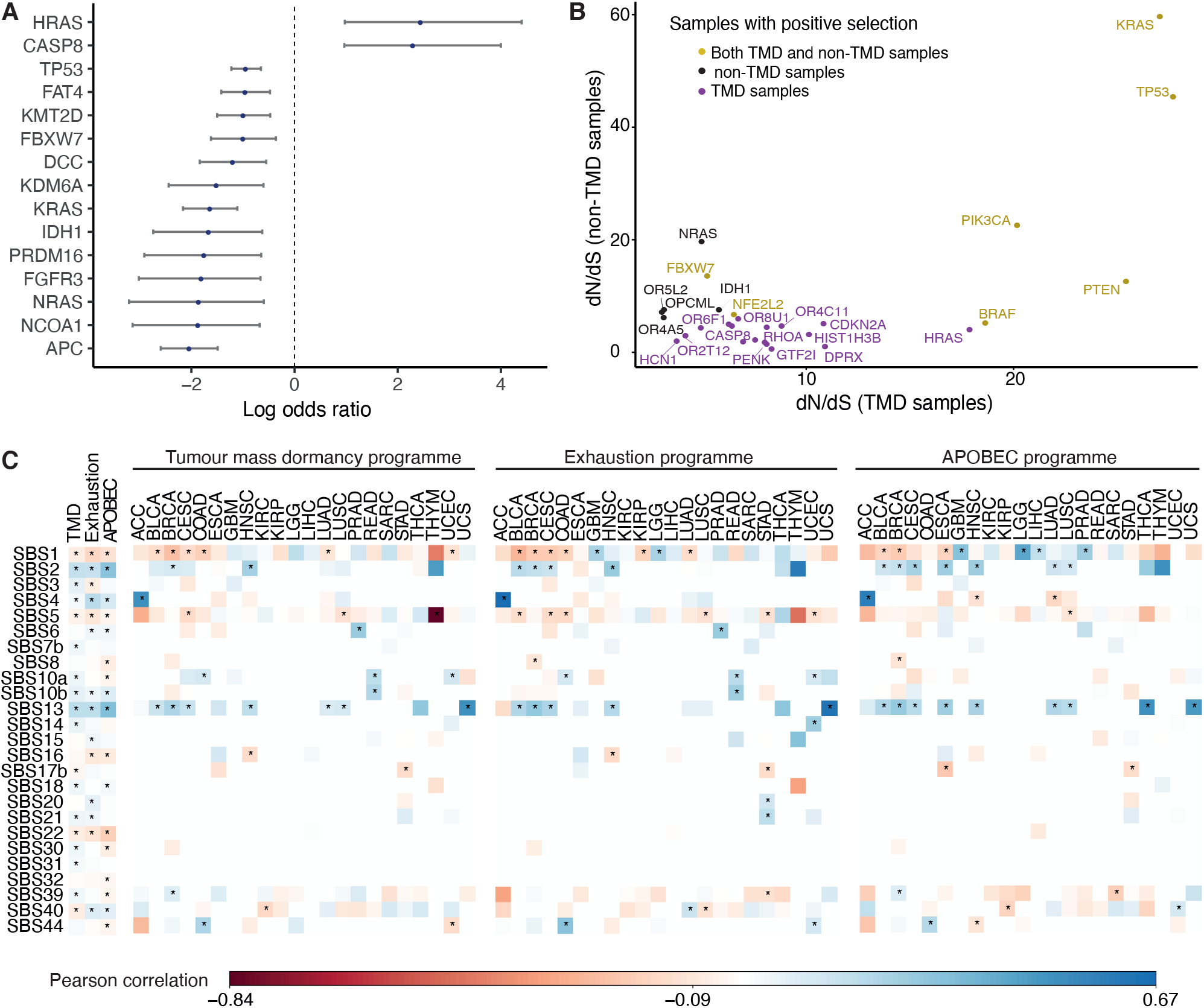
Genomic drivers and mutational processes linked with TMD. **(A)** Genes presenting an enrichment or depletion of mutations within high TMD samples. Blue circles represent odds ratios on a log2 scale and the confidence intervals for each of the individual Fisher’s exact tests are depicted. **(B)** Genes showing signals of positive selection in samples with high (purple) and low (black) TMD, as well as across both groups (yellow). **(C)** Matrices depicting the Person correlation between mutational signatures and the TMD, exhaustion and APOBEC programmes, both pan-cancer (first 3 columns) and within individual cancer tissues. Statistically significant correlations (p<0.05) are highlighted with an asterisk. Only signatures with at least one significant correlation are shown.

The results of the pan-cancer analysis largely reflected the balance of proliferation, cell death and pro/anti-angiogenic signals (Naumov et al. 2006; Semenza 2003; Aguirre-Ghiso 2007) one would expect in the context of dormancy. Two genes stood out as having a ~5-fold enrichment of mutations in patients displaying TMD: *HRAS* and *CASP8*. Interestingly, while *HRAS* alterations were positively associated with dormancy, other members of the same oncogenic family, *KRAS* and *NRAS* showed a depletion of mutations in the context of this phenotype. Specifically, we found an enrichment of *HRAS* hotspot mutations at the Q61 and G13 positions across all solid primary tumour samples, but a depletion of *KRAS* G13 and G12, as well as *NRAS* Q61 hotspot alterations (Supplementary Figure 13). Two of these hotspots presented cancer stage specificity: *HRAS* G13 mutations were enriched in TMD in late-stage cancers, while *KRAS* G12 mutations were depleted in early-stage tumours (Supplementary Figures 9B, 10B). The oncogenic activation of the Ras protein has been associated with pro-angiogenic signaling through repression of thrombospondin-1 (Watnick et al. 2015). It has been suggested that while distinct oncogenic Ras alterations might have similar ability to promote cell cycle progression, they might have different abilities to induce the pro-angiogenic programme (Aguirre-Ghiso 2007), which may explain the discrepancy of mutation signals in the *RAS* genes in relation to TMD.

*CASP8*, a gene encoding a cysteine-aspartic acid protease involved in the execution of apoptosis by cleaving and thereby activating caspase-3 and caspase-7 (Tummers and Green 2017), was also preferentially mutated in dormancy. While *CASP8* loss of function would be predicted to impair the ability of cancer cells to initiate apoptosis, silencing of *CASP8* in breast cancer cell lines has also been shown to decrease cancer cell growth by delaying G0/G1-to S-phase transition and increasing the expression of CDK inhibitors p21 and p27 (De Blasio et al. 2016).

The genes presenting a depletion of point mutations within dormant samples identified by our analysis have key functions in regulating tumour growth, including *TP53*, the master regulator that coordinates signals of stress such as DNA damage and aberrant growth signaling and can induce cell cycle arrest or apoptosis (Vogelstein, Lane, and Levine 2000; Polyak et al. 1997), or *APC*, which suppresses tumour growth through repression of the Wnt signaling pathway (Boman and Fields 2013). Both the *APC* and *DCC* genes promote apoptosis by downregulation of survivin gene expression (T. Zhang et al. 2001) and caspase-9 cleavage (Forcet et al. 2001) respectively. Control of tumour vascularization was reflected in depletion of mutations in *PRDM16*, which inhibits angiogenesis by suppressing the expression of a HIF target semaphorin 5B (Kundu et al. 2020), and in *NCO1A*, a transcriptional coactivator that upregulates the expression of the VEGFα pro-angiogenic factor (Qin et al. 2015). Mutations in the *IDH1* gene, shown to be depleted in samples with TMD, result in the production of the 2-hydroxyglutarate metabolite which regulates the activity of α-ketoglutarate dependent dioxygenases and causes the ubiquitination and proteasomal degradation of *HIF1A*, a key sensor of hypoxia and initiator of angiogenesis (Ye et al. 2013). Moreover, 2-hydroxyglutarate can also result in the downregulation of leukocyte chemotaxis factors (Turcan et al. 2012; Tommasini-Ghelfi et al. 2019), which could contribute to the ability of tumour cells to escape immunological dormancy through immune system evasion.

In addition to the enrichment analysis, using a maximum-likelihood dN/dS method (Martincorena et al. 2017) we detected signals of positive selection for mutations within *CASP8* and *HRAS* in samples with TMD, but not in expanding tumour samples (Figure 2B). In contrast, *IDH1* and *NRAS*, both of which showed a depletion of mutations within TMD samples, showed signals of positive selection only in expanding tumours (Figure 2B). Interestingly, *CASP8* showed similarly increased mutation rate both in early- and late-stage cancers with TMD, while *HRAS* mutations appeared linked with TMD only in late-stage tumours – pointing towards a timing specificity of TMD-linked evolutionary pressures (Supplementary Figure 14). The associations between mutations in these genes and TMD were also validated in an independent dataset from the International Cancer Genome Consortium (ICGC), along with several others including *KRAS* and *TP53* (Supplementary Figure 15). While signals of positive selection are more difficult to validate in the sparser independent cohorts available due to the limited sample size, nevertheless *CASP8* and *TP53* selection signals were robustly recovered both in oral cancers from ICGC as well as in breast cancers from the METABRIC cohort (Supplementary Figure 16). These findings support the importance of *CASP8* and *RAS* mutational status in the context of TMD.

In terms of broader structural variation in the genome, we found no copy number alteration events (amplifications and deletions) specifically enriched in tumours with TMD (data not shown), potentially because such events would be preferentially selected for in fast growing tumours. Moreover, tumours with angiogenic and immunological dormancy showed a modest, but significant decrease in mutational burden when compared to tumours without TMD (Supplementary Figure 17).

### 2.3 Mutational processes linked with dormancy and exhaustion

In addition to investigating the links with specific driver events, we also set out to characterise broader mutational processes associated with TMD. Different risk factors of cancer induce DNA damage in the cells in a context dependent manner, such that nucleotide substitution patterns associated with such mutational processes can be observed within cancer genomes. These patterns of trinucleotide substitutions are termed “mutational signatures” and have been widely characterised across cancers (Alexandrov, Nik-Zainal, Wedge, Aparicio, et al. 2013; Alexandrov et al. 2020). We carried out mutational signature analysis to survey the contribution of known mutagenic processes and risk factors to the genomes of dormant tumours. The mutational signature prevalence was correlated with dormancy and exhaustion programme scores (Figure 2C, left and centre panels).

We observed that exposure to signatures SBS1, originating from aging-induced deamination of 5-methylcytosines (Alexandrov et al. 2015), SBS5, also ageing-linked, and SBS22, linked with aristolochic acid exposure (Jelaković et al. 2015), decreased as TMD and exhaustion increased. In contrast, smoking (SBS4) and defective base excision repair linked with polymerase epsilon or *NTHL1* mutations (SBS10a/b, SBS30) (Jager et al. 2019; Shivji et al. 1995) were associated with an increase in TMD. Finally, we noted a consistently strong correlation between TMD/exhaustion and mutational signatures SBS2 and SBS13, associated with mutagenesis induced by a class of cytidine deaminases called APOBEC (apolipoprotein B mRNA editing catalytic polypeptide-like) (Alexandrov et al. 2020). These enzymes induce mutagenesis in viral genetic material and are part of anti-viral defence, but can also act on and damage host DNA (Green and Weitzman 2019). The correlation of these mutational footprints of APOBEC with TMD was observed both pan-cancer and across individual cancer types, including bladder, breast, cervical and head and neck cancers (Figure 2C). Reassuringly, there was a similarly strong correlation between SBS2 and SBS13 and the mean expression of the AID/APOBEC cytidine deaminases (Figure 2C, right panel). All the correlations highlighted were consistent between early- and late-stage cancers (Supplementary Figure 18).

### 2.4 Dormancy and exhaustion programmes correlate with APOBEC enzymatic activity

To further explore the association between TMD and exhaustion with APOBEC mutagenesis, we calculated the correlation of our programme scores and the mean expression of genes belonging to the APOBEC family (Figure 3A-F). Across all cancers and regardless of clinical stage, we observed significant correlations between the activity of the APOBEC programme as a whole, TMD and exhaustion programmes (Figure 3 A-G, Supplementary Figures 19-23). These associations were stronger than would be expected by chance (Supplementary Figure 24) and consistently validated in independent datasets from ICGC and cBioPortal (Supplementary Figures 25-28). In particular, we observed a marked association between APOBEC mutagenesis and immunological dormancy, which we stipulate may precede or come in conjunction with immune exhaustion, the latter showing the strongest correlation.

**Figure 3.**
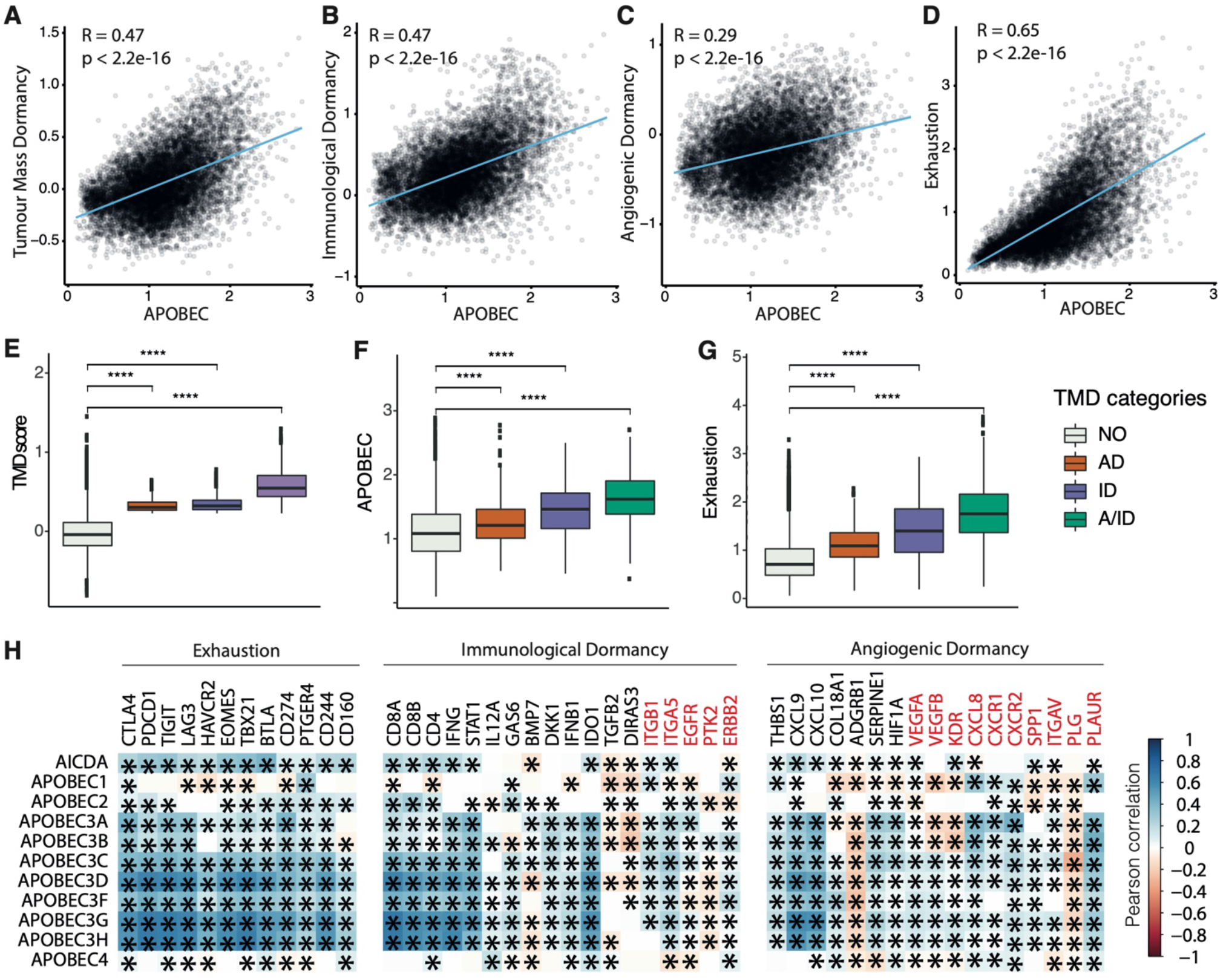
TMD and exhaustion programmes correlate with APOBEC transcriptional activity. **(A-D)** Pan-cancer correlation between the activity of the APOBEC programme and **(A)** TMD, **(B)** immunological dormancy, **(C)** angiogenic dormancy and **(D)** exhaustion programmes. **(E-G)** TMD, APOBEC and exhaustion programme scores compared between samples with angiogenic dormancy (AD), immunological dormancy (ID), both angiogenic and immunological dormancy (A/ID) and expanding tumours without evidence of TMD (NO); **** p<0.0001. **(H)** Pan-cancer correlation between individual genes in the dormancy and exhaustion programmes and APOBEC/AID enzyme family gene expression. Genes downregulated in the immunological and angiogenic dormancy programmes are shown in red. Significant correlations are marked with an asterisk.

We also observed positive correlations between most but not all genes included in the immunological, angiogenic and exhaustion programmes, on the one side, and the expression of the AID/APOBEC enzyme family members, on the other (Figure 3H). This may be because of downstream regulation of expression, e.g. through mRNA degradation, compartmentalization, or inhibition of transcription. Notably, among all genes, the correlation was highest for the mRNAs of enzymatically active APOBEC3 and AID members of the APOBEC family. The correlation scores were similar for *APOBEC3A-H*, which is to be expected as these genes form a cluster on chromosome 22. Within the TMD programme, type II interferon ɣ gene and the interferon-upregulated genes such as *STAT1* showed a positive correlation with APOBEC. This is interesting as interferon ɣ signaling is associated with anti-tumour response by favoring rejection of highly immunogenic tumours (Mittal et al. 2014; Benci et al. 2016). However, prolonged interferon ɣ signaling promotes epigenetic changes to *STAT1* and promotes expression of ligands for multiple T cell inhibitory receptors (Benci et al. 2016). Thus, we would expect a spectrum of exhaustion signaling to be present within the tumour, ranging from cells signaling danger to cells with anti-cytotoxic response. Anti-angiogenic markers, such as the urokinase receptor *PLAUR*, which downregulate angiogenic processes and thus restrict the supply of T cells (Oh, Hoover-Plow, and Plow 2003), were also positively correlated with APOBEC expression. Similar associations were observed for several immune checkpoints (*CD244*, *CD166*, *TIGIT*, *CTLA4*, *PDCD1*) and exhaustion markers (*EOMES*, *TBX21*).

Overall, the multiple correlations between APOBEC expression and the dormancy/exhaustion levels in tumours suggest a complex interplay between the activity of APOBEC deaminases and the immune microenvironment, which we set out to explore in greater depth.

### 2.5 TMD activity differences in the context of APOBEC mutagenesis

To further elucidate the association between TMD and APOBEC activity, we sought to investigate the specific TMD signals that might have the strongest links with APOBEC-attributable mutagenesis, as quantified by the single-base substitution mutational signatures SBS2 and SBS13. In order to identify APOBEC-enriched tumours, we employed t-distributed stochastic neighbour embedding (tSNE) to cluster samples based on their overall mutational signature profiles (Figure 4A-B). Unsurprisingly, the clustering was impacted by the tumour tissue of origin, as mutational signatures are often tissue-specific (Degasperi et al. 2020) (Figure 4A). However, samples enriched for APOBEC-associated mutations, defined as the total enrichment of signatures SBS2 and SBS13, covered a fairly heterogenous set of tissues (Figure 4B). Groups with distinct mutagenesis patterns were defined using expectation-maximization clustering (Supplementary Figure 29), and the procedure was repeated 100 times to obtain robust clusters. Samples were defined as APOBEC-enriched if they fell into the APOBEC-associated cluster more than 50 times (see Materials and Methods), which broadly overlaps with the APOBEC enrichment score approach developed by Roberts et al (Roberts et al. 2013) (Supplementary Figure 30).

**Figure 4.**
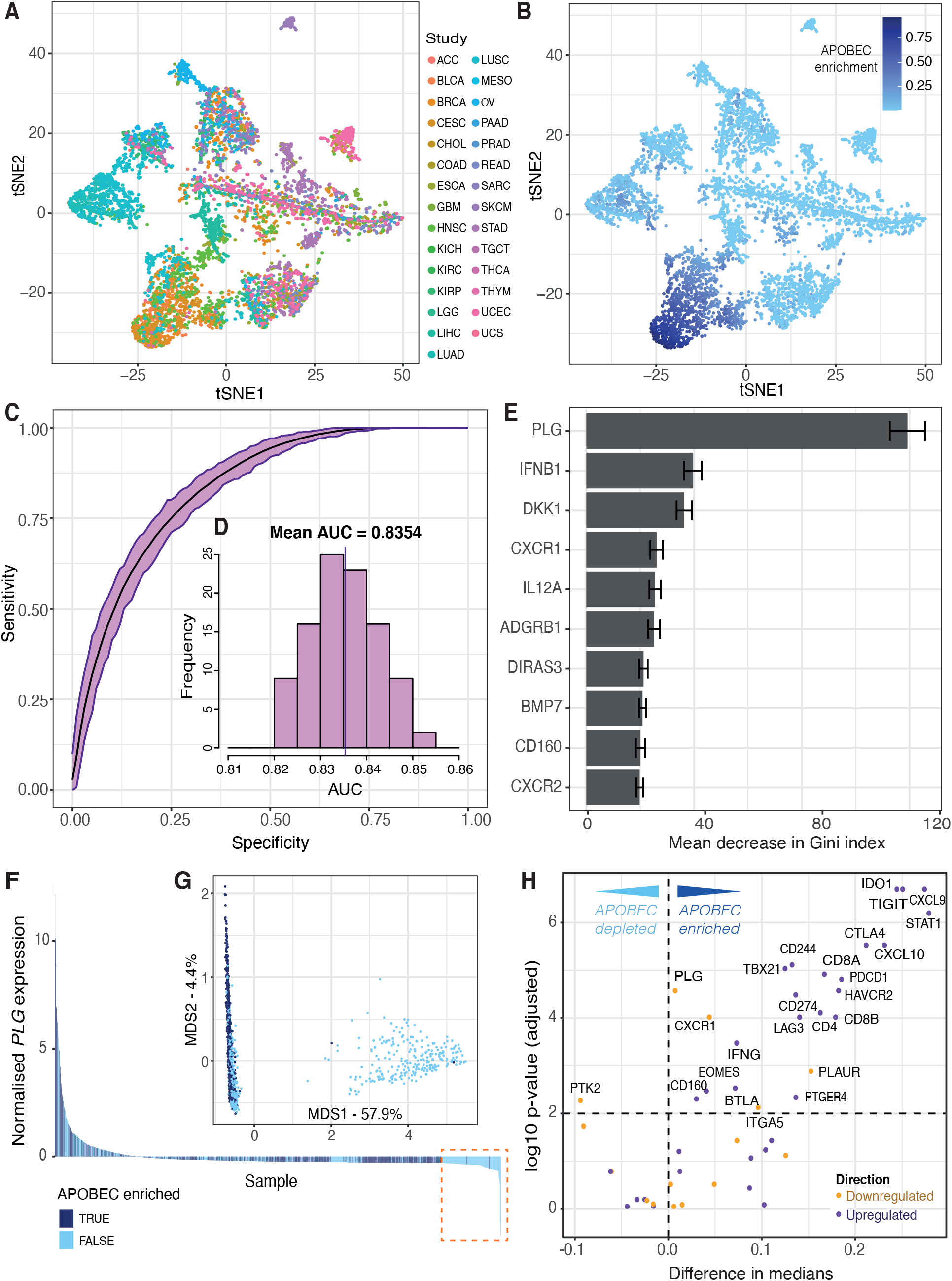
TMD and exhaustion programmes distinguish an APOBEC mutagenesis cluster. tSNE dimensionality reduction of 6,410 primary tumour samples based on single-base substitution signature profiles labelled by **(A)** TCGA study and **(B)** the total contribution of mutations from APOBEC signatures SBS2 and SBS13. **(C)** Receiver operating characteristic (ROC) curves of 100 random forest classifiers of APOBEC signature enrichment based on expression of genes involved in the TMD and exhaustion programmes. **(D)** Distribution of Area Under Curve (AUC) values across all 100 random forest classifiers. **(E)** The top 10 ranked genes with highest importance for APOBEC signature enrichment classification, ranked by mean associated decrease in Gini index across all 100 classifiers. **(F)** Waterfall plot displaying samples included in an exemplary classifier ranked by normalised *PLG* expression, coloured by APOBEC enrichment labels. The dashed orange box highlights the subset of samples with low PLG expression and marked depletion of APOBEC signatures. **(G)** Multidimensional scaling plot displaying samples included in the exemplary classifier. **(H)** Volcano plot displaying the difference in median normalised expression for each gene involved in the TMD and exhaustion programmes between APOBEC-enriched and non-enriched groups, coloured by anticipated direction of regulation. Differences in medians above 0 correspond to an increase in expression in the APOBEC-enriched group. Genes with a difference in median <0 would instead be higher expressed in the non-enriched cluster (APOBEC depleted). A significance cut-off of p<0.01 was applied after Benjamini-Hochberg multiple testing correction.

To determine whether APOBEC mutagenesis was specifically associated with particular aspects of the TMD programme, we used random forest classifiers to rank TMD genes based on how much their expression helps to distinguish between the APOBEC-enriched and depleted groups (see Materials and Methods). The accuracies for predicting APOBEC enrichment across 100 built models were remarkably high and broadly replicable (mean AUC = 0.8354, range: 0.8203-0.8534) (Figure 4C-D). Additionally, the gene ranking in the models was also highly reproducible (Figure 4E). *PLG*, which encodes for the plasminogen protein and triggers angiostatin release to inhibit angiogenesis (Oh, Hoover-Plow, and Plow 2003), displayed the highest importance by far, defined as the mean associated decrease in Gini index following its removal from a model (mean = 119.2). Upon further investigation of *PLG* expression, we found that its high predictive importance could largely be attributed to a substantial group of low-*PLG*-expressing samples which were not labelled as APOBEC-enriched (Figure 4F). This becomes more apparent when considering the multidimensional scaling plot attributable to one of the random forest classifiers, which clearly displays a segregated cluster of non-APOBEC-enriched samples defined by a specific *PLG* expression threshold derived from the classifier (Figure 4G). Therefore, APOBEC mutagenesis appears to be completely lacking in the cases when *PLG* is not expressed, suggesting a link between APOBEC enzymatic activity and the uPA/uPAR system of angiogenic modulation.

Finally, we compared the expression of genes associated with TMD between APOBEC mutagenesis enriched and depleted groups. The large majority of genes whose upregulation drives TMD displayed a higher median expression in APOBEC-enriched samples compared with non-enriched samples (Figure 4H). It is worth noting that some genes were found to be significantly changed whilst displaying negligible difference in medians, likely reflecting the effects of a large sample size rather than a real biological difference. Overall, these results reaffirm the positive correlation between TMD and APOBEC enzyme activity, apparent both from an expression and mutagenesis perspective.

### 2.6 Heterogeneity of TMD maintenance in hypoxic conditions

Owing to the reported association between hypoxic environments and angiogenesis (L. Chen, Endler, and Shibasaki 2009), we next sought to investigate whether hypoxia might differentially impact TMD. Hypoxia was quantified on a per-sample basis using established transcriptional signatures (see Materials and Methods), and the calculation procedure was validated using three separate hypoxia signatures (Supplementary Figure 31).

From a pan-cancer perspective, we observed that hypoxia scores were significantly lower in samples displaying any type of dormancy, compared to samples with no evidence of TMD (Figure 5A). This was further corroborated by grouping tumours into distinct subsets (bins) based on their hypoxia levels: the proportion of samples with no evidence of TMD increased significantly as hypoxia scores increased (Figure 5B). In particular, we noted the lowest hypoxia levels appearing in tumours with angiogenic dormancy. These results align with the established consensus that the hypoxia-inducible factor (HIF) pathway is a key regulator of angiogenesis (Pugh and Ratcliffe 2003; Krock, Skuli, and Simon 2011), with angiogenic dormancy expected to develop in normoxic conditions. Similar patterns were also observed on a tissue-specific basis, particularly in the cases of sarcoma, stomach, lung squamous and esophageal carcinomas (Figure 5C). Cancer studies which tended to present low hypoxia, such as pheochromocytoma and paraganglioma (PCPG), thymoma (THYM) and thyroid carcinoma (THCA) often presented some evidence of TMD, whereas studies presenting high hypoxia, such as colon and rectum adenocarcinomas (COAD/READ) and glioblastoma (GBM), were abundant in expanding tumours (Supplementary Figure 32). Notable exceptions included bladder cancer and uveal melanoma, both of which presented samples with evidence of TMD at the higher end of their respective hypoxia spectra (Figure 5C-F). While prior research has highlighted HIF-1α expression as a promoter of angiogenesis and tumour invasion in bladder carcinoma (Theodoropoulos et al. 2004) and metastasis in uveal melanoma (Asnaghi et al. 2014), our analysis suggests that a period of TMD-induced latency may also be compatible with hypoxia in these cancers.

**Figure 5.**
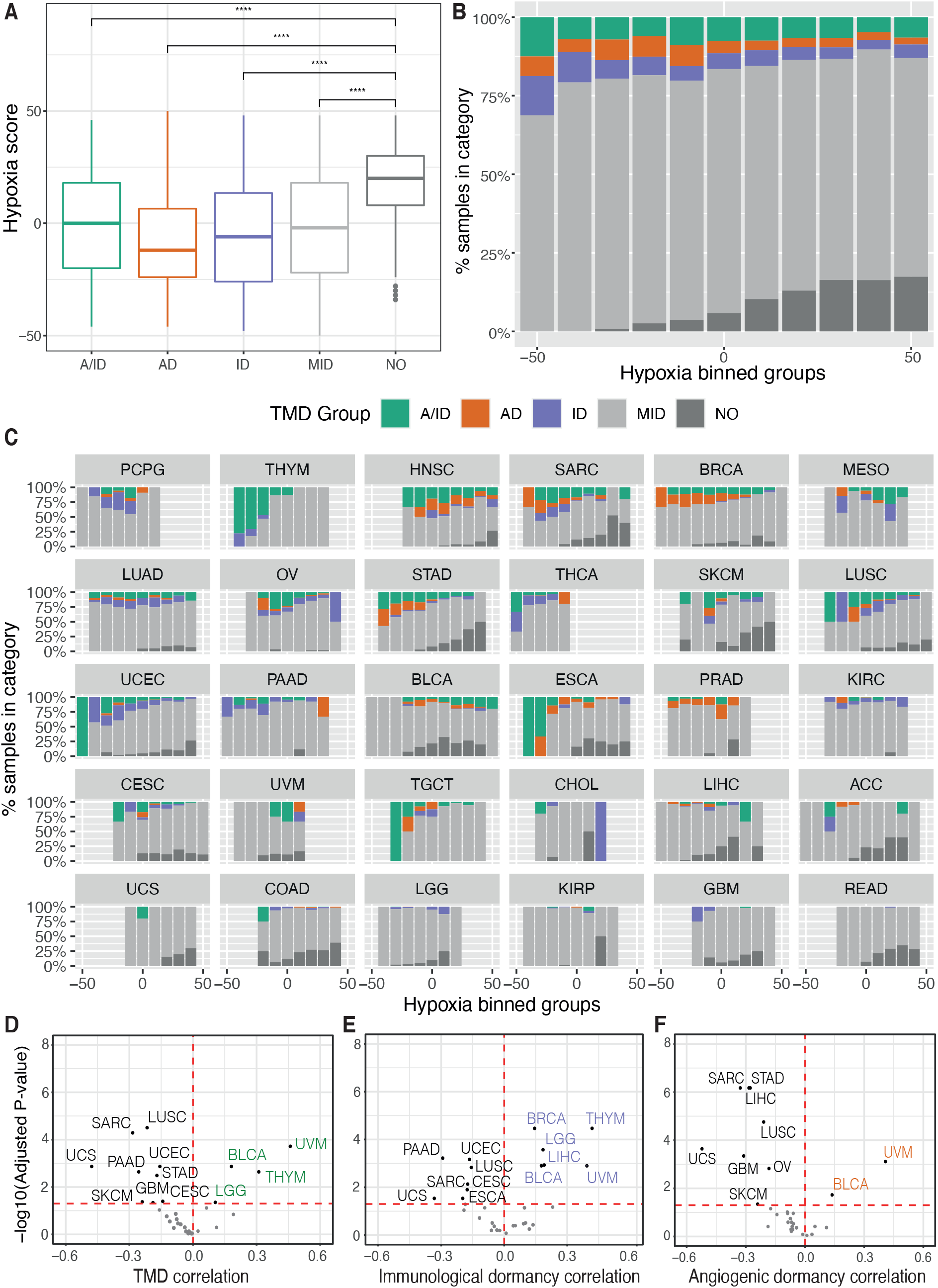
TMD is reduced in the context of hypoxia. **(A)** Comparison of hypoxia scores across samples falling within distinct TMD categories: both angiogenic and immunological dormancy (A/ID), angiogenic dormancy (AD), immunological dormancy (ID), slowly expanding tumours (MID), expanding tumours without evidence of TMD (NO). **** p<0.0001. **(B-C)** Distributions of dormancy status within hypoxia groups, defined by binning the hypoxia scores into intervals of 10: **(B)** pan-cancer, and **(C)** by TCGA cancer tissue, arranged in descending proportion of dormant samples. KICH was not plotted because it lacked TMD samples. **(D-F)** Volcano plots displaying the cancer-study-specific Pearson correlations of **(D)** TMD, **(E)** immunogenic dormancy and **(F)** angiogenic dormancy programme scores against hypoxia scores. Cancers showing an increased dormancy in the context of hypoxia are highlighted in corresponding colours, while cancers with high dormancy in normoxic conditions are depicted in black.

While a considerable heterogeneity across cancer tissues was observed, patterns of decreased hypoxia in the context of TMD were noted in the majority of cancers (Figure 5D-F). This would imply that hypoxia may be an impediment to TMD maintenance across the majority of tumours, consistent with its increased prevalence in the later stages of tumour development (Petrova et al. 2018).

### 2.7 The microenvironmental context of TMD

Since TMD is by definition dependent on immune surveillance, we also set out to confirm this and potentially identify new components of the cancer microenvironment which are permissive to this type of dormancy. Specifically, because the balance in the populations of immune and stromal cells is central to forming anti-tumour responses, we investigated how immune composition might be shaped in the context of TMD, exhaustion and APOBEC programmes. Using cell type-specific transcriptional markers (see Materials and Methods), we observed strong positive correlations between TMD and various types of T cells (Figure 6A). Antitumoral cytotoxic cells such as CD8+ and CD4+ T cells, Th1/2 cells, as well as tumour promoting regulatory cells and macrophages were correlated with the APOBEC, exhaustion and dormancy programmes. This observation is in line with the mechanism proposed in the literature whereby an initial immune response caused by neoantigens presented by the dormant tumours is followed by exhaustion of such signals (Ghorani et al. 2020). In addition to their cytotoxic activity, CD8+ and CD4+ T cells have been previously shown to limit tumour growth through the secretion of anti-proliferative cytokines, such as IFN- *γ* which can stimulate the expression of p21 and p27 cell cycle inhibitors (Wall et al. 2003; Chin et al. 1996), and antiangiogenic chemokines, such as CXCL9 and CXCL10 (Ikeda, Old, and Schreiber 2002).

**Figure 6.**
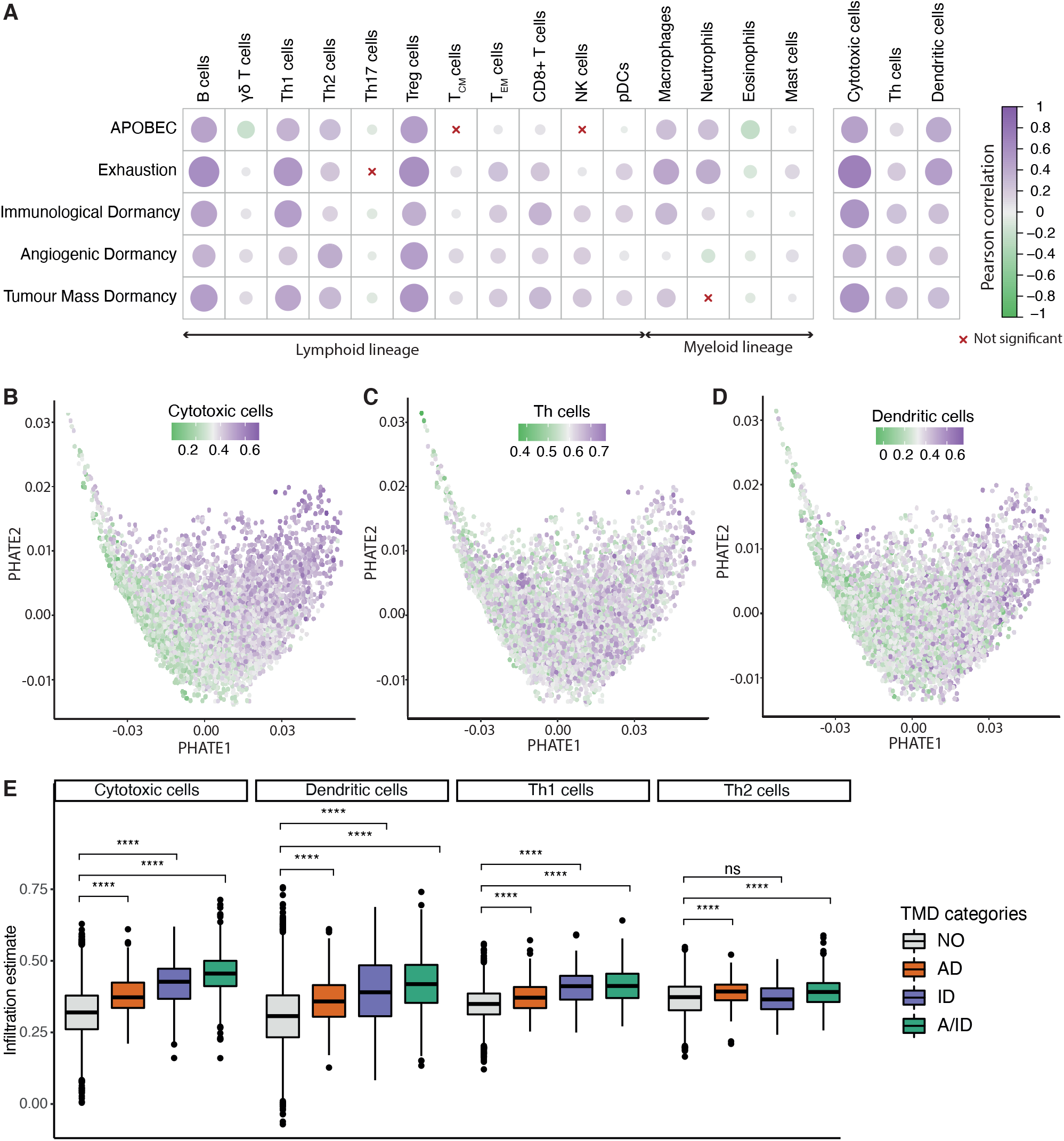
Tumour microenvironment activity correlates with the TMD programme. **(A)** Correlation between cell infiltration estimates and the APOBEC, exhaustion and dormancy programme scores across the TCGA primary tumor samples. The three broader categories on the right hand side summarise the combined marker expression for all cytotoxic, T helper and dendritic cells. **(B-D)** PHATE dimensionality reduction of 9,631 primary tumor samples, based on the expression of genes driving the TMD and exhaustion programmes, with removal of tissue specific expression patterns and coloured by **(B)** cytotoxic cell enrichment score, **(C)** Th cell enrichment score and **(D)** dendritic cell enrichment score. **(E)** Cytotoxic T cell, Th1/Th2 and dendritic cell abundance compared between samples with angiogenic dormancy (AD), immunological dormancy (ID), both angiogenic and immunological dormancy (A/ID) and samples without TMD (NO); **** p<0.0001.

Compared to Th1 and Th2 cells, natural killer (NK) cells showed a weaker correlation with dormancy programmes, consistent with reports of tumour growth control by the immune system being mostly associated with adaptive T cell responses (Finn 2006). The microenvironment of samples with high TMD was also depleted in inflammatory Th17 cells, which secrete the angiogenesis inducer IL-17A (Bailey et al. 2014). Interestingly, samples displaying signals of angiogenic dormancy had higher enrichment of Th2 instead of Th1 cells. While the Th1 activity is linked with IFN-ɣ production and tumour suppression, Th2 activity has been implicated in cancer progression (Zhao et al. 2019). Despite the broader heterogeneity of T helper cell signals, they showed a similar overall trend as that of cytotoxic and dendritic cells when projected across the TMD landscape (Figure 6B-D). All these classes except for the Th2 cells appeared most enriched in tumours with evidence of both immunological and angiogenic dormancy (Figure 6E). Similar associations were observed for early- and late-stage cancers (Supplementary Figures 33-36).

### 2.8 TMD is prognostic in cancer

To understand the clinical relevance of TMD, we carried out survival analysis and found that patients with expanding tumours that presented no evidence of TMD had a significantly reduced prognosis (Figure 7A), consistent with the expectation that fast-proliferating tumours should be more aggressive than their dormant counterparts. To further dissect the potential sources of this variation along the immunity/angiogenesis axis, we employed Cox proportional hazards models to determine the effects of different types of dormancy on survival whilst accounting for potential confounders such as patient age, gender, cancer type, and tumour stage. While all three categories were associated with decreased risk when compared to patients with no evidence of TMD, much of this variation was accounted for by tumour stage, with only angiogenic and tumour mass dormancy still marginally significant after correction (Figure 7B, Supplementary Figures 37-41). While the tumour stage effect may not be fully uncoupled from the TMD effect, TMD did present a marginal protective role within the early-stage tumours (stages I/II), as well as a larger protective role in late-stage tumours (stages III/IV) (Supplementary Figure 42).

**Figure 7.**
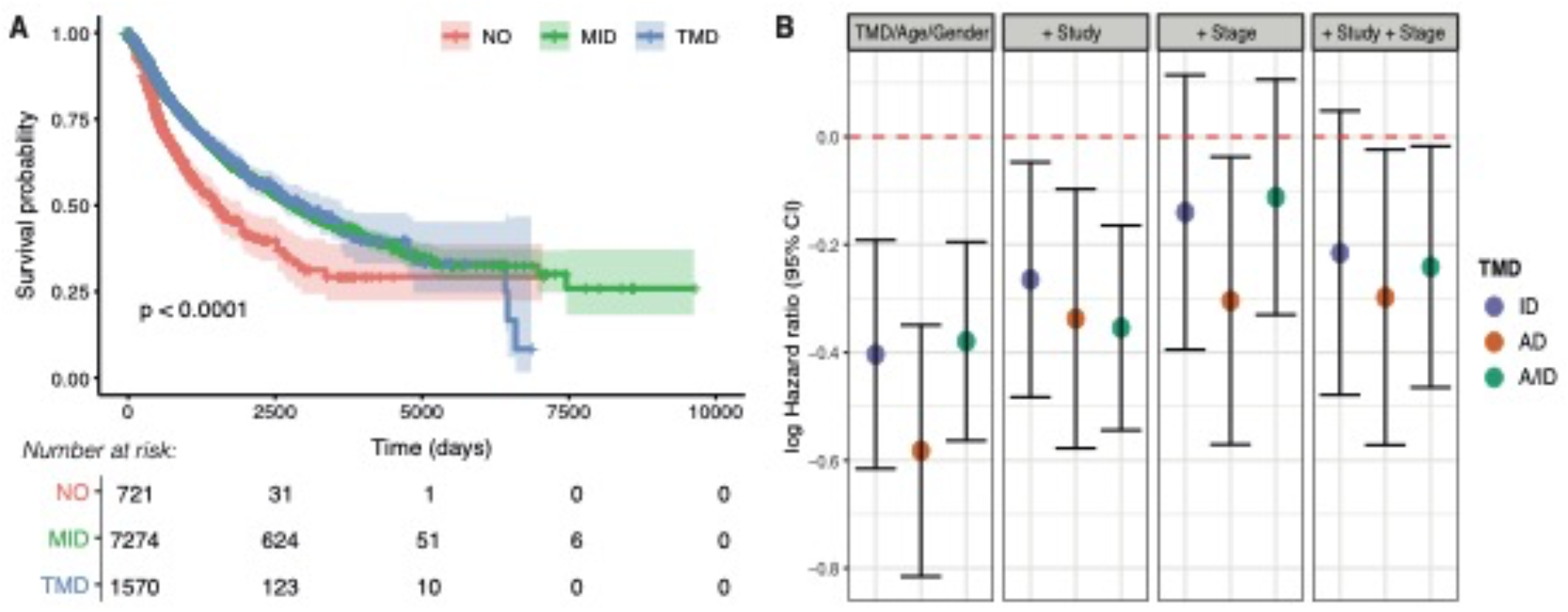
TMD is linked with better prognosis pan-cancer. **(A)** Overall survival compared between tumours with distinct levels of TMD: strong evidence for TMD (TMD), middle/weak evidence (MID) and no evidence (NO). The number of patients at risk for the corresponding time points are displayed below. **(B)** Log hazard ratios of survival for tumours with angiogenic dormancy (AD), immunological dormancy (ID), both angiogenic and immunological dormancy (A/ID) compared to the ‘NO’ dormancy category, whilst accounting for age and gender, followed by the inclusion of TCGA study, tumour stage (early/late), and both TCGA study and tumour stage.

## 3 DISCUSSION

TMD is a key state in tumour development and progression, but its evolutionary constraints are largely unknown. In this study, we have presented an overview of TMD across 31 cancer tissues and highlighted potential novel genomic hallmarks of this state. While TMD remains a poorly characterised programme whose current known markers are undoubtedly incomplete, our approach to evaluating TMD in bulk tumours through an equilibrium of proliferation, apoptosis, cytotoxicity, exhaustion and angiogenesis signals has enabled us to systematically capture this temporary state across a variety of cancers. We show that TMD-like signals are present across a multitude of cancer tissues, and that immunological and angiogenic dormancy are not always concomitant. We confirm many of the expected associations reported in the literature, including lower mutational burden, correlations with CD8+, regulatory and helper T cells (Teng et al. 2008), improved prognosis (Park and Nam 2020) and a strong link with immune exhaustion (Mittal et al. 2014). Exhaustion is a phenotype that likely immediately follows TMD, but in our analysis the two states are largely overlapping. This is most likely due to the bulk nature of the samples analysed, where we may be capturing parts of the tumour that starting to expand in addition to areas still displaying TMD. Interestingly, we find TMD tends to occur in normoxic rather than hypoxic environments in the majority of cases, which is different than what is observed for cellular quiescence (Qiu et al. 2017; Butturini et al. 2019).

In addition to expected associations, our methodology has also enabled us to capture previously unreported links between TMD and clinical or genomic characteristics. Firstly, while TMD would be expected to dominate in early-stage tumours, we also notice TMD-like signals in a comparable fraction of late-stage tumours (18%) (Supplementary Figure 8). In the absence of longitudinal data that could shed further light onto the causes, we can only hypothesise this could be due to a mechanism of slowly advancing cancers that are maintained in a semi-TMD state, potentially enabled by a strong, consistent immune response.

Secondly, we find that oncogenic mutations in *CASP8* (inactivating) and *HRAS* (activating) are positively selected for during the development of TMD. This is either because they might confer an advantage in terms of maintaining TMD, or because they might enable cells to escape TMD and expand further. Interestingly, the selection pressure on *CASP8* in the context of TMD was seen across the entire tumour progression spectrum, while *HRAS* mutations appeared positively selected specifically in late-stage tumours. Counterbalancing these selective forces are *KRAS* hotspots that are preferentially depleted in the context of TMD. These findings could suggest distinct adaptive mechanisms in the context of early forming versus disseminated cancer in a temporary dormant state, as well as a complex “tug-of-war”-like regulation of tumour growth/arrest via Ras oncogenesis. Beyond these mutational associations, most of the classical cancer drivers were amplified or deleted preferentially in non-dormant tumours, while TMD appeared largely devoid of specific copy number marks. These observations may be partly explained by an association with early stage cancers, where these changes have not yet occurred as a means of expansion and escape from immune surveillance.

In concordance with the reduced rate of cell proliferation in TMD, we found that ageing-induced mutations, which are believed to accumulate at a relatively constant rate in cancer, were lesser represented in dormant tumours. Correlations between TMD and several other mutational processes (smoking, defective DNA repair) were noted, which might suggest that multiple neoplastic mechanisms that are tissue-specific could favour this state.

A remarkably consistent signal came from the mutagenesis trace left by the APOBEC enzyme family. APOBEC activity appeared strongly linked with the exhaustion signals that presumably follow TMD, and with immunological dormancy. Several studies (Yamazaki et al. 2019; Swanton et al. 2015; Burns, Temiz, and Harris 2013) have proposed that APOBEC expression may be one of the drivers of tumorigenesis, suggesting a role for APOBECs in evading immune response. APOBEC3 signatures are prevalent in leukaemias (Rebhandl et al. 2014), while AID was demonstrated to induce hypermutation and chromosomal translocation in B cell lymphomas (Robbiani and Nussenzweig 2013). On the other hand, APOBEC may promote the formation of neoantigens and activate cytotoxic T cell responses. It was shown that the APOBEC3 class of enzymes is upregulated in immune cells (monocytes, macrophages and pDCs) upon treatment with interferon ɣ (F.X. Wang et al. 2008; Stenglein et al. 2010). In ovarian carcinoma, the expression of APOBEC3G was positively correlated with T cell infiltration, expression of cytotoxic granzyme and perforin (*GZMB*, *PRF1*), and improved clinical outcome (Leonard et al. 2016). Moreover, APOBEC mutagenesis has previously been linked with responses to immunotherapy in lung, melanoma and head and neck cancers (S. Wang et al. 2018; Faden et al. 2019; Driscoll et al. 2020) through the induction of heteroclitic neoepitopes. We have demonstrated increased APOBEC mutagenesis in a TMD setting across multiple tumour types, and have shown that this process appears linked with the regulation of angiogenesis by the uPAR system. It is possible that the mutational footprint left in the genome by such activity enables immune recognition and maintenance of TMD in the time span until exhaustion and immune escape occurs. This could suggest a window of opportunity for the employment of checkpoint inhibitors, possibly in combination with APOBEC inhibitors, in early-stage tumours with TMD.

Our pan-cancer analysis unveils a complex interplay between TMD, exhaustion and APOBEC activity, and a mutational context that may enable these phenotypes to develop. However, it is possible that some of the associations uncovered may be due to the direct association between APOBEC activity and immune responses/exhaustion or hypoxia, which have been previously reported in the literature as highlighted above, rather than with tumour mass dormancy itself. Future experimental studies will be needed to verify and further understand the implications of increased APOBEC mutagenesis in a TMD-specific context. Moreover, changes in gene expression may not accurately reflect changes in abundance of protein products in our defined dormancy and exhaustion programmes. Biochemical assays and studies in model organisms will give a more definite answer to the nature of these relationships.

An important limitation of this analysis consists in the fact that there are no currently available datasets describing transcriptional changes specifically in the context of tumour mass dormancy. This phenotype is difficult to study due to its temporary nature and few model systems for this state exist. We have therefore not been able to unequivocally determine whether the chosen scoring method and even the cut-offs employed for assessing TMD capture this phenomenon in the most accurate manner. However, we are taking a conservative approach focusing only on the most extreme values, which are most likely to reflect this state. This approach appears methodologically robust to small fluctuations in expression/gene signatures, and the associations identified have been validated in independent datasets. Moreover, the key findings of *CASP8* positive selection in TMD, associations with APOBEC mutagenesis and cytotoxic microenvironment were also reproducible using the PCA-based scoring methodology (Supplementary Figures 43-48). Further experimental evidence is needed to verify the biological nature of these associations and the most appropriate way of evaluating TMD, once experimental model systems for this state will have become more widely available and easier to study.

Another limitation refers to the fact that the samples analysed have been profiled using bulk sequencing methods, which means that we are only capturing an average cellular signal across the entire tumour which may not reflect its true heterogeneity. Furthermore, since TMD is a temporary phenotype, a single static snapshot of tumours, even in this large dataset, may not be sufficient to capture the entire array of TMD phenotypes that might develop in different tissues. Therefore, multi-regional sequencing and longitudinal studies should be conducted in the future to shed further light on the diversity and dynamics of TMD phenotypes.

Furthermore, our study focused mainly on pan-cancer markers of TMD, due to the scarcity of this phenotype. From our analyses, we observe TMD is frequent in certain tissues such as head and neck, breast, bone, and has weak evidence or complete absence of signals in others, e.g. chromophobe renal cell carcinoma. Some of the smaller studies from TCGA may be insufficiently powered to capture such signals. Therefore, future studies should focus on an in-depth characterization of TMD in distinct tissues and should be able to pinpoint tissue-specific genomic markers linked to TMD. Finally, the potential genomic biomarkers of dormancy that we identified need to be confirmed through experimental validation in suitable in vitro/in vivo models of TMD.

Overall, our study provides evidence for TMD, immunological and angiogenic dormancy signals across a variety of cancer types, and highlights key associated intrinsic and extrinsic hallmarks, including *CASP8*/*RAS* dependencies, APOBEC mutagenesis and hypoxia. These findings pave the way for further exploration of the mechanisms underlying TMD emergence, maintenance and exit.

## 4 MATERIALS AND METHODS

### 4.1 Datasets

FPKM normalised RNA-sequencing expression data as well as mutation annotation files aligned against the hg38 genome from the Muse pipeline, were downloaded using the *TCGAbiolinks* R package (Colaprico et al. 2016) for 9,712 treatment-naïve primary tumour TCGA samples across 31 solid cancer types. For patients with multiple samples available, one RNA-seq barcode entry was randomly selected for each individual patient resulting in 9,631 total entries. All expression data were log-transformed for downstream analysis.

For validation, RNA-seq and matched whole-genome sequencing data were downloaded for 12 cancer studies from the ICGC Data Portal (J. Zhang et al. 2019), accompanied by RNA-seq and matched targeted sequencing from cBioPortal (Cerami et al. 2012) for the following datasets: blca_mskcc_solit_2012 = Bladder Cancer (MSKCC, J Clin Onco 2013); brca_metabric = Breast Cancer (METABRIC, Nature 2012 & Nat Commun 2016); prad_mskcc = Prostate Adenocarcinoma (MSKCC, Cancer Cell 2010); sarc_mskcc = Sarcoma (MSKCC/Broad, Nat Genet 2010); whole-genome sequencing from rt_target_2018_pub = Pediatric Rhabdoid Tumor (TARGET, 2018) and prostate_dkfz_2018 = Prostate Cancer (DKFZ, Cancer Cell 2018); and whole-exome sequencing from: brca_smc_2018 = Breast Cancer (SMC 2018); GIS031 = Lung adenocarcinoma (GIS, Nat Genet 2019); paad_qcmg_uq_2016 = Pancreatic Adenocarcinoma (QCMG, Nature 2016); prad_broad = Prostate Adenocarinoma (Broad/Cornell, Nat Genet 2012); utuc_cornell_baylor_mdacc_2019 = Upper Tract Urothelial Carcinoma (Cornell/Naylor/MDACC, Nat Commun 2019); wt_target_2018_pub = Pediatric Wilm’s Tumor (TARGET, 2018). The data were processed and analysed in the same manner as the TCGA data.

### 4.2 Quantifying the dormancy, exhaustion and APOBEC programmes

Mean log-transformed expression values of genes deemed to be associated with a given programme by manual curation of the literature (Supplementary Table 1) were used to produce per-sample programme scores for TMD, immunological/angiogenic dormancy, exhaustion and APOBEC activity. We have specifically selected genes that have been associated with immunological and angiogenic <u>dormancy</u>, rather than generic immunity or angiogenesis processes, in order to ensure that any associations identified downstream are likely to be TMD-related.

The APOBEC and exhaustion programme scores were calculated by taking the mean expression of genes within the respective programmes, as all genes in the programmes were expected to be upregulated in the respective states. The angiogenic and immunological dormancy (AD/ID) programme scores, as well as the TMD scores, were calculated using two different approaches:

(a) Scaled difference of means

For every sample in the cohort, a score *S* was derived by subtracting the sum of expression values of downregulated genes (*E_d_*) in the respective programme from the sum of expression values of upregulated genes (*E_u_*) and dividing by the total number of genes within the programme (*N_u_*+*N_d_,* where *N_u_* is the total number of upregulated genes and *Nd* is the total number of downregulated genes):

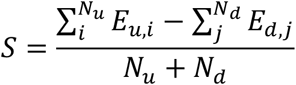

(b) Principal component analysis (PCA)

PCA was performed across the pan-cancer dataset using the up- and downregulated gene signature of the respective programme. The dormancy score was extracted as the coordinate of the first principal component (PC1).

The proliferation/apoptosis ratio was calculated by dividing the average expression of E2F target genes (including MKI67, a classical proliferation marker (Miller et al. 2018)), obtained from the “HALLMARK_E2F_TARGETS” gene list, by the average expression of genes involved in apoptosis from the “HALLMARK_APOPTOSIS” gene list. Both gene lists are part of the “H: hallmark gene sets” collection deposited at MSigDB.

Samples with high TMD were identified as being within the upper quartile of the TMD programme score range and presenting a ratio of proliferation/apoptosis < 1. Dormant samples were further classified as having immunological and/or angiogenic dormancy based on whether they were in the upper quartile of the angiogenic (AD) and/or immunological dormancy (ID) programme score range, respectively. Samples with no evidence for TMD (expanding tumours, marked as ‘NO’) were identified as being in the lower quartile of the TMD programme score range and showing a ratio of proliferation/apoptosis > 1. The remaining samples (‘MID’ category) were classed as having middle levels of expansion with some weak potential evidence of dormancy, but biologically unlikely to be dormant.

### 4.3 Assessing the robustness of the AD/ID programme scores

To assess the relative robustness of the three methodologies employed to calculate the TMD and AD/ID scores, we took two approaches:

(1) We systematically removed one gene from the respective programme at a time and recalculated the scores.
(2) A small amount of random noise was added to the expression of each gene in the signature, measured in each sample and scores were recalculated. The jitter R function was used to introduce variable amounts of noise in the data, by changing the factor variable level between 1-200). For each level of noise, the approach was repeated 100 times and the scores were recalculated.

The overall variability of the fold changes in the new score compared to the original scores was compared between the ‘scaled difference of means’ and PCA-based methods.

### 4.4 Assessing the robustness of APOBEC associations

Correlations between the mean expression of APOBEC-related genes and random gene expression programmes were calculated by randomly selecting an equal number of genes to that found in the TMD signatures (35) from the genome and assessing the mean expression correlation with APOBEC activity. 1000 iterations were performed. Associations varied as shown in Supplementary Figure 24, and were consistently lower than the one calculated for TMD. P-values were frequently significant, but this is likely due to the large number of samples.

### 4.5 PHATE dimensionality reduction

The *phateR* R package (Moon et al. 2019) was used to perform the dimensionality reduction based on the expression of genes associated with the TMD and exhaustion programmes. A constant seed was used for reproducibility. The *ComBat* function from the *sva* R package (Leek et al. 2012) was used to remove tissue-specific expression patterns from the TCGA RNA-seq data.

### 4.6 Association between cancer driver mutations and TMD

The COSMIC database (Tate et al. 2019) was used to source a list of 723 known drivers of tumorigenesis (Tiers 1+2). COSMIC genes mutated in at least 1% of samples across all solid primary tumour samples were tested for enrichment or depletion of mutations between samples with high and low TMD using a Fisher’s exact test. Only missense, nonsense, nonstop, frameshift deletion/insertion and inframe insertion/deletion mutations were considered in the analysis. For *HRAS*, *KRAS* and *NRAS*, a Fisher’s exact test was also performed to test for enrichment or depletion of specific recurrent hotspot mutations, reported by the cBioPortal data hub and based in part on methodology from Chang et al (Chang et al. 2016) and Gao et al (Gao et al. 2017). The analysis was also repeated on a cancer-by-cancer basis, where COSMIC genes mutated in at least 5% of samples within the cancer-specific study sample were tested for enrichment or depletion of mutations with TMD samples. The p-values were corrected using Benjamini-Hochberg (BH) procedure to accommodate for multiple testing.

Enrichment analysis was reaffirmed using a random forest classification approach. This was conducted using the *randomForest* R package. A balanced dataset of samples with high and low TMD was generated by utilising all samples with low TMD (n = 657), and randomly sampling an equal number of samples with high TMD (n = 1,154). The resulting random forest consisted of 500 trees, and mutation enrichment was determined by calculating the mean decrease in Gini index between the two groups following the removal of a gene.

To identify genes that are positively selected in the context of TMD, we carried out dN/dS analysis separately for high and low TMD samples using the *dNdScv* R package (Martincorena et al. 2017), run with default parameters.

### 4.7 Mutational signature analysis

Mutational signature contributions were inferred using *deconstructSigs* (Rosenthal et al. 2016) and the choice of signatures was further informed using results from *SigProfiler* (Alexandrov, Nik-Zainal, Wedge, Campbell, et al. 2013). Only samples with at least 50 mutations were employed in the analysis, for a total of 6,410 samples.

*SigProfiler* was used to infer mutational signatures from TCGA whole-exome sequencing data. For each TCGA study of interest, input mutational matrices were generated using the *SigProfilerMatrixGeneratorR* function using all samples containing at least 50 mutations and aligning the MAF files to the hg38 genome build. *SigProfilerExtractorR* was used to extract de novo mutational signatures for each cancer type. For each study, the solution with the greatest number of mutational signatures was chosen, for which also the sum of the solution stability (calculated average silhouette coefficient) and minimum stability exceeded 1, and the minimum stability value did not fall below 0.4. For the selected solutions, the identity of the mutational signatures was determined by calculating cosine similarities with COSMIC v3.1 mutational signatures.

Mutational signatures from whole-exome sequencing data were also inferred using the *deconstructSigs* R package. MAF files were aligned against the hg38 genome build and the COSMIC v3.1 mutational signatures were employed in the analysis. For each cancer type, the contribution of all signatures within the *deconstructSigs* solution was set to 0, apart from SBS1, SBS5, signatures identified in the corresponding *SigProfiler* solution, as well as signatures which contribute on average at least 5% of mutations across all samples within the *deconstructSigs* solution. For cancer types where *SigProfiler* did not result in a stable solution, the *deconstructSigs* solution was used with the contribution of all signatures set to 0, apart from SBS1, SBS5 as well as signatures which contribute at least 5% of mutations across samples within the *deconstructSigs* solution.

### 4.8 Tumour microenvironment deconvolution from bulk RNA-seq data

The tumour microenvironment cell infiltration scores were calculated using the *ConsensusTME* R package (Jiménez-Sánchez, Cast, and Miller 2019) based on gene sets from Bindea et al (Bindea et al. 2013). Cell abundance was estimated from the TCGA bulk RNA-seq data using singe sample Gene Set Enrichment Analysis (ssGSEA). Broader categories (cytotoxic cells, Th cells, dendritic cells) were scored by combining the expression across subtype-specific markers.

### 4.9 Identification of APOBEC mutagenesis clusters

First, APOBEC enrichment scores were calculated using the *sigminer* R package (Shixiang Wang et al. 2020). Next, the mutational signature profiles of 6,410 samples underwent dimensionality reduction with tSNE using the *Rtsne* R package, followed by expectation-maximization clustering using the *EMCluster* R package. The optimal number of clusters was determined by considering the associated increase in log-likelihood pertaining to additional clusters and was assumed to be equivalent for all applications of tSNE (Supplementary Figure 9A). Whilst substantial increases were associated with both 10 and 17 clusters, we selected 10 clusters in order to prevent the cluster of APOBEC-enriched samples from being segmented. The APOBEC-enriched cluster was identified as the cluster with the highest median collective enrichment of APOBEC signatures SBS2 and SBS13. The procedure was conducted 100 times and samples which appeared in the APOBEC-enriched cluster on more than 50 occasions were labelled as ‘APOBEC enriched’.

### 4.10 Random forest modelling of APOBEC mutagenesis-gene expression associations

Z-score normalisation of RNA-sequencing expression data was applied across each cancer type individually. The FPKM-normalised values were log-transformed, and the results for each gene were transformed into a Z ~ N(0,1) distribution using the scale function in R. Only genes involved in the dormancy and exhaustion programmes were considered as input in the classifier.

Genes whose expression contributed to APOBEC enrichment classification were identified using machine learning via a random forest approach, conducted using the *randomForest* R package. In total, 5,850 samples were included for which Z-scores and mutational signature profiles were available. Each random forest model required all samples labelled as APOBEC-enriched (n = 952), and an equal number of samples taken randomly without replacement from the non-APOBEC-enriched samples to generate balanced groups. Each random forest consisted of 1,000 trees. The importance of each gene in the model, measured as the associated mean decrease in Gini index following its removal, was also calculated. This procedure was run 100 times.

Changes in median gene expression between the APOBEC enriched and non-enriched groups (Figure 4H) were determined using a two-sample Wilcoxon rank-sum test. P-values were corrected for multiple testing using the Benjamini-Hochberg procedure.

### 4.11 Calculation of hypoxia scores

Hypoxia scores were calculated using three transcription-based hypoxia signatures referred to as Buffa (Buffa et al. 2010), West (Eustace et al. 2013) and Winter (Winter et al. 2007), following the calculation procedure detailed by Bhandari et al (Bhandari et al. 2019). Genes which were included in the curated dormancy or exhaustion signatures were subsequently removed from the respective hypoxia signature. Scores were calculated based on the pan-cancer cohort to enable variation in hypoxia between cancer types to be identified. Since we observed a high degree of agreement between the three signatures (Supplementary Figure 11), the Buffa signature was employed in the downstream analysis on account of its use as a reference by Bhandari et al. A binning approach with a window size of 10 was employed to further subdivide tumours into discrete subgroups with increasing levels of hypoxia.

### 4.12 Survival Analysis

Survival analysis was conducted via the *survival* R package. Hazard ratios were calculated using Cox proportional hazards models and were displayed using the *finalfit* R package. Survival curves were calculated using the Kaplan-Meier formula, and plotted using the *survminer* R package.

### 4.13 Statistics

Pairwise correlations were calculated using the Pearson correlation statistics. The *corrplot* R package was used to carry out statistical analysis and visualise the correlation matrices. Groups were compared using the Student’s t test, Wilcoxon rank-sum test or Kruskal-Wallis test as appropriate. Multiple testing correction was applied using the Benjamini-Hochberg method. The significance threshold was taken as p < 0.05.

### 4.14 Code

All code developed for the purpose of this analysis can be found at the following repository: https://github.com/secrierlab/tumourMassDormancy/tree/v1.0

## 5 Conflict of Interest

The authors declare that the research was conducted in the absence of any commercial or financial relationships that could be construed as a potential conflict of interest.

## 6 Author Contributions

MS designed and supervised the study. AJW, DHJ and WL performed the analyses. All authors wrote and approved the manuscript.

## 7 Funding

AJW and DHJ were supported by MRC DTP grants (MR/N013867/1). MS was supported by a UKRI Future Leaders Fellowship (MR/T042184/1), an Academy of Medical Sciences Springboard Award (SBF004\1042) and a Wellcome Trust Seed Award in Science (215296/Z/19/Z).

## 8 Data Availability Statement

The results published here are based upon data generated by the TCGA Research Network: https://www.cancer.gov/tcga.

## Notes

https://github.com/secrierlab/tumourMassDormancy/tree/v1.0

### Competing Interest Statement

The authors have declared no competing interest.

### Summary of Updates

(1) Implementation of an alternative method for scoring TMD based on principal component analysis, comparison with the original method and re-analysis of the cohort using this latter method, which confirmed the associations previously identified (2) Extensive validation of the mutation and APOBEC associations with TMD in independent datasets from the International Cancer Genome Consortium (ICGC) and cBioPortal (3) Re-analysis of the TCGA data in early and late-stage cancers separately, which confirmed the previously observed associations across cancer stages and helped further interpret selection pressures in the context of TMD (4) Extension of discussion to acknowledge more clearly the limitations stemming from lack of experimental validation and the already reported links between APOBEC and tumour immunity

