## Supplementary Material for "Pan-cancer survey of tumour mass dormancy and underlying mutational processes"

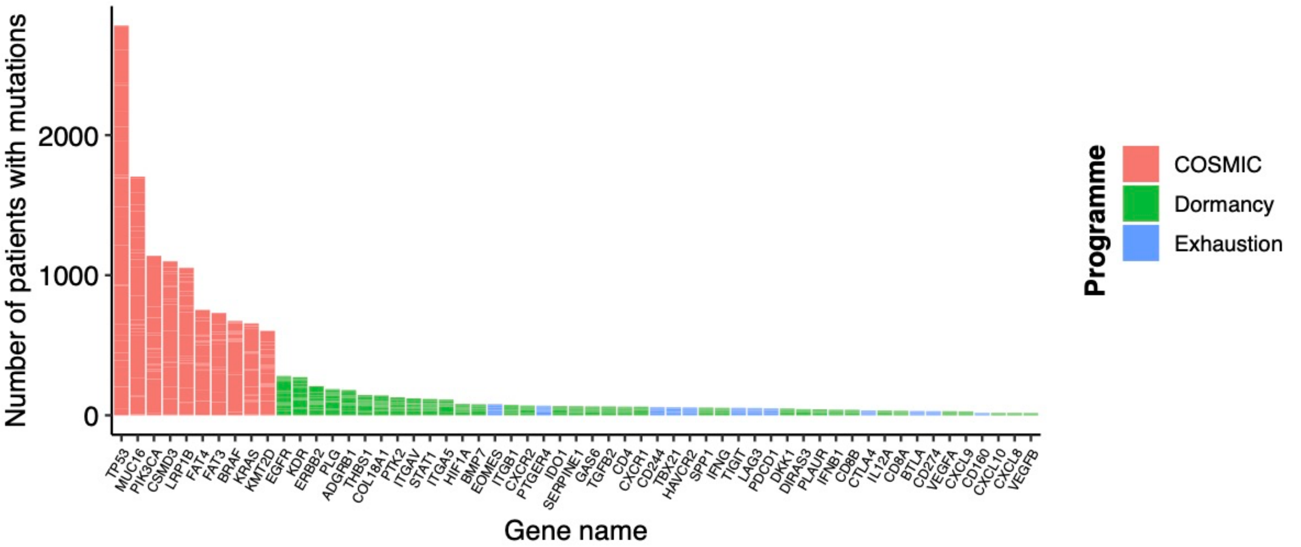


**Supplementary Figure 1. Frequently mutated canonical drivers of cancer compared to markers of dormancy/exhaustion.** The frequency of mutations within genes making up the tumor mass dormancy (green) and exhaustion (blue) programmes within the TCGA cohort solid cancer samples is compared to top 10 most frequently mutated tier 1 and 2 COSMIC cancer drivers (red).

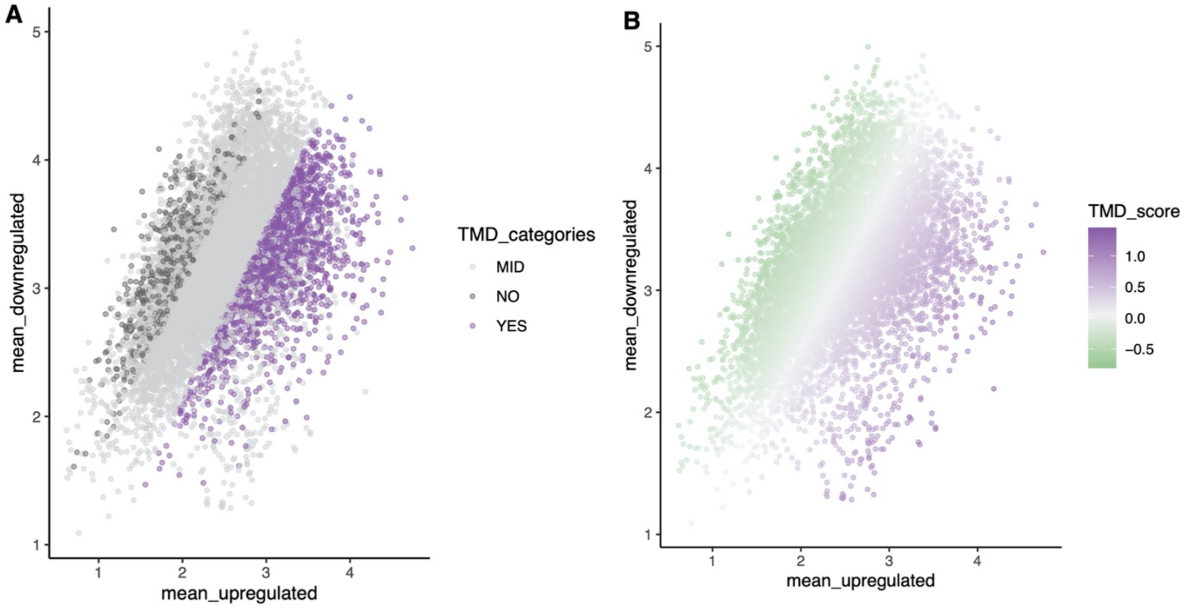


**Supplementary Figure 2. Variation in TMD scores based on expression of up- and downregulated genes involved in the angiogenic and immunological dormancy programmes.** The programme labels are highlighted for each sample in the pan-cancer cohort based on variation of upregulated genes (mean expression shown on the x axis) and of downregulated genes (mean expression shown on y axis): **(A)** TMD classification. **(B)** TMD continuous score.

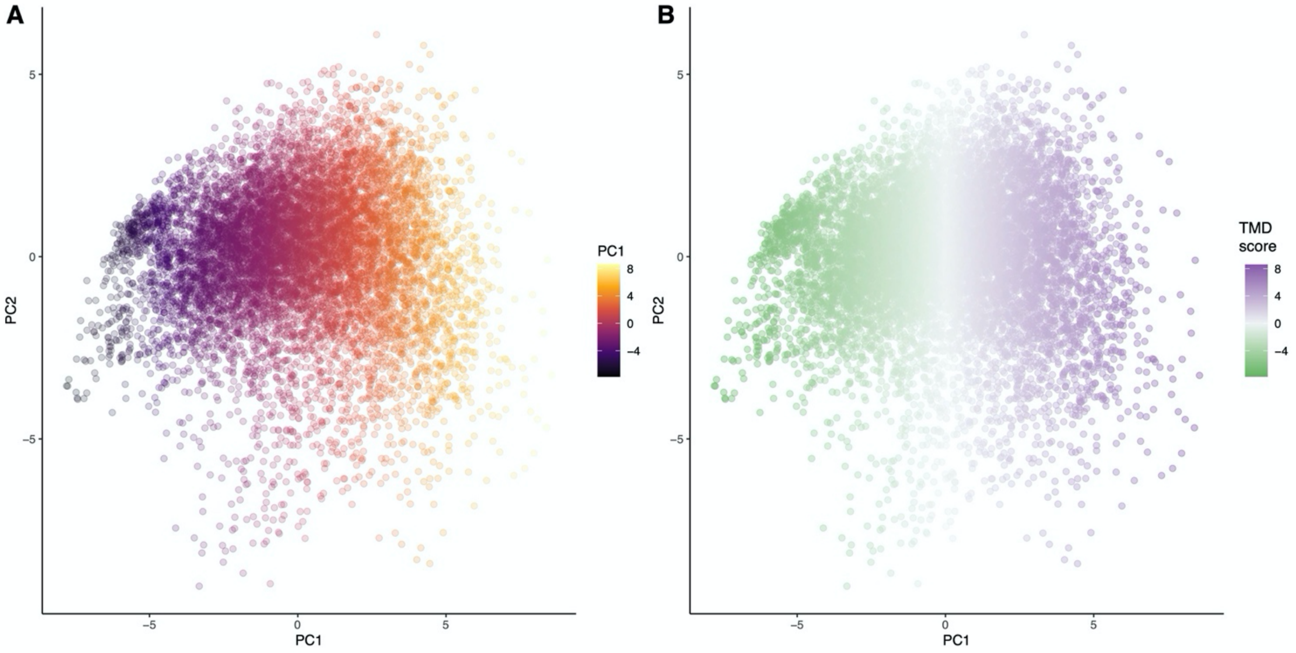


**Supplementary Figure 3. Variation in TMD scores based on PCA. (A)** PCA plot based on the pan-cancer expression of genes included in the dormancy signatures. The variation across the first two principal components is used. **(B)** TMD score variation across the first principal component (PC1).

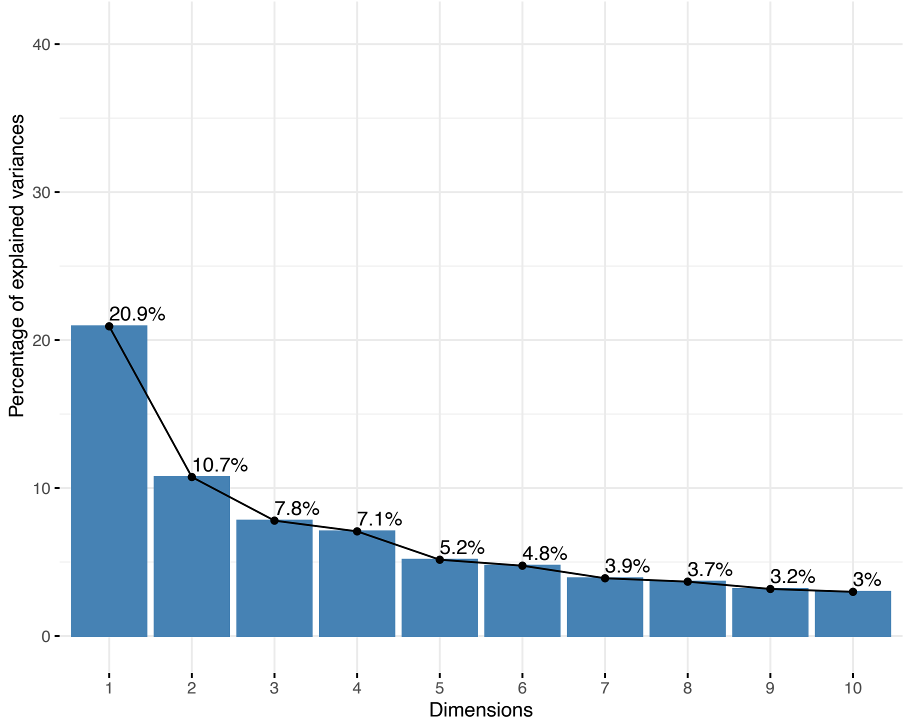


**Supplementary Figure 4. Percentage of explained variance in TMD-related genes by the first 10 principal components.**

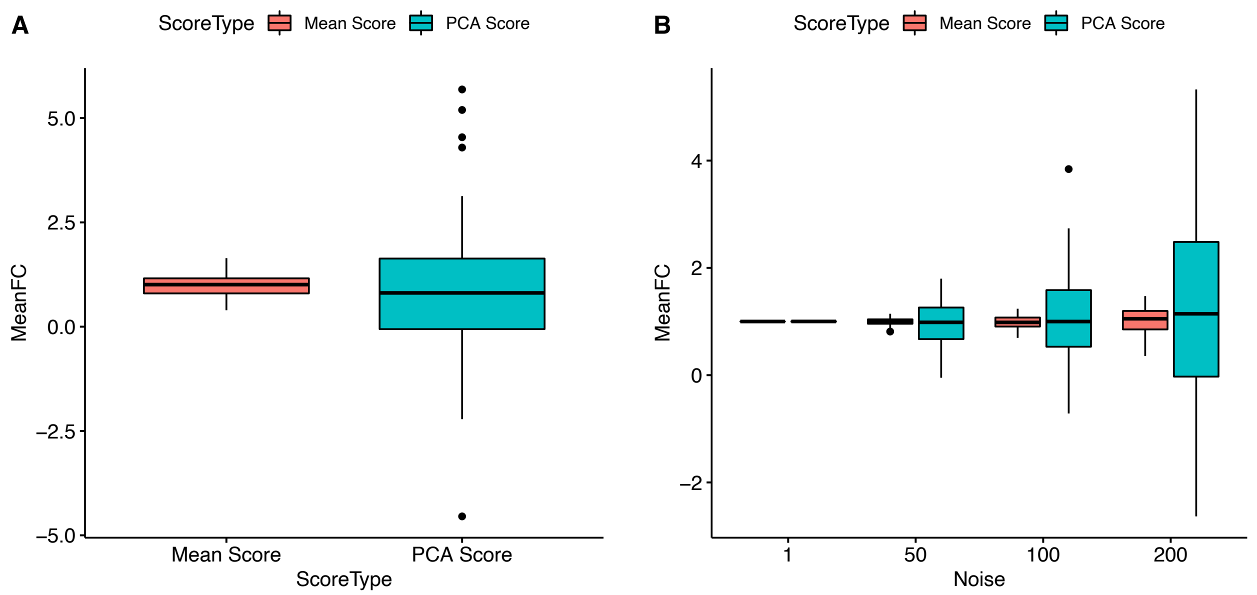


**Supplementary Figure 5. Robustness of employed methodologies. (A)** The mean pan-cancer fold change in TMD score per sample when randomly removing one gene at a time from the signature. **(B)** The mean pan-cancer fold change in TMD score per sample when introducing increasing amount of noise to the measured gene expression data. The mean-based scoring technique appears to be more robust to variations in gene signatures or gene expression.

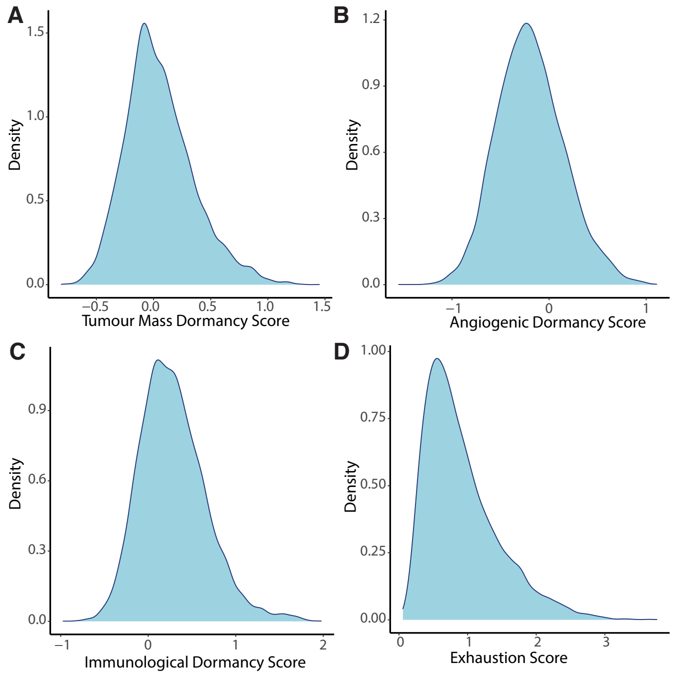


**Supplementary Figure 6. Pan-cancer density distribution of exhaustion and dormancy scores. (A)** TMD score distribution. **(B)** Angiogenic dormancy score distribution. **(C)** Immunological dormancy score distribution. **(D)** Exhaustion score distribution.

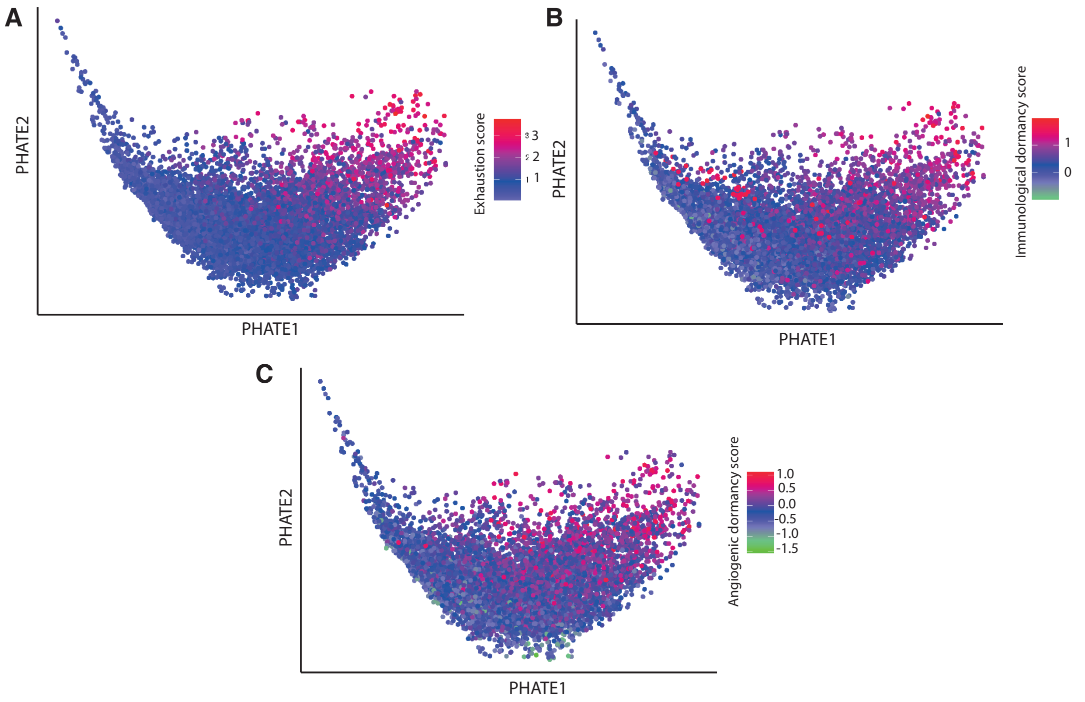


**Supplementary Figure 7. Pan-cancer evaluation of exhaustion, immunological dormancy and angiogenic dormancy programmes.** PHATE dimensionality reduction was applied to 9,631 primary tumor samples based on the expression of genes within the tumor mass dormancy and exhaustion programmes after the removal of tissue specific expression patterns. The maps are colored by their corresponding exhaustion **(A)**, immunological dormancy **(B)** and angiogenic dormancy **(C)** programme scores.

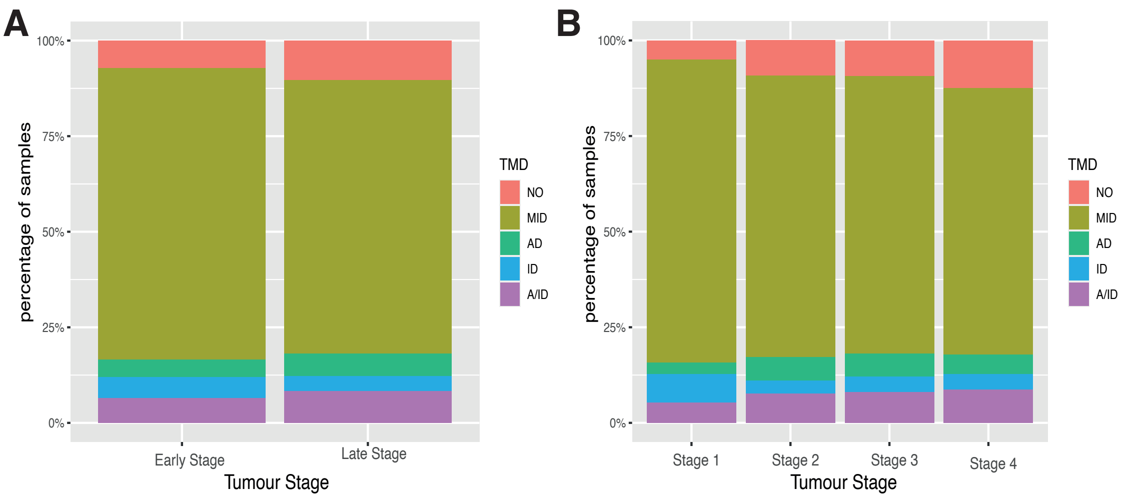


**Supplementary Figure 8. The proportion of samples in each TMD category across cancer stages. (A)** Samples are split into early/late stage cancers. **(B)** Samples are split into stages I to IV. Stages I and II constitute ‘early stage’, and stages III and IV constitute ‘late stage’.

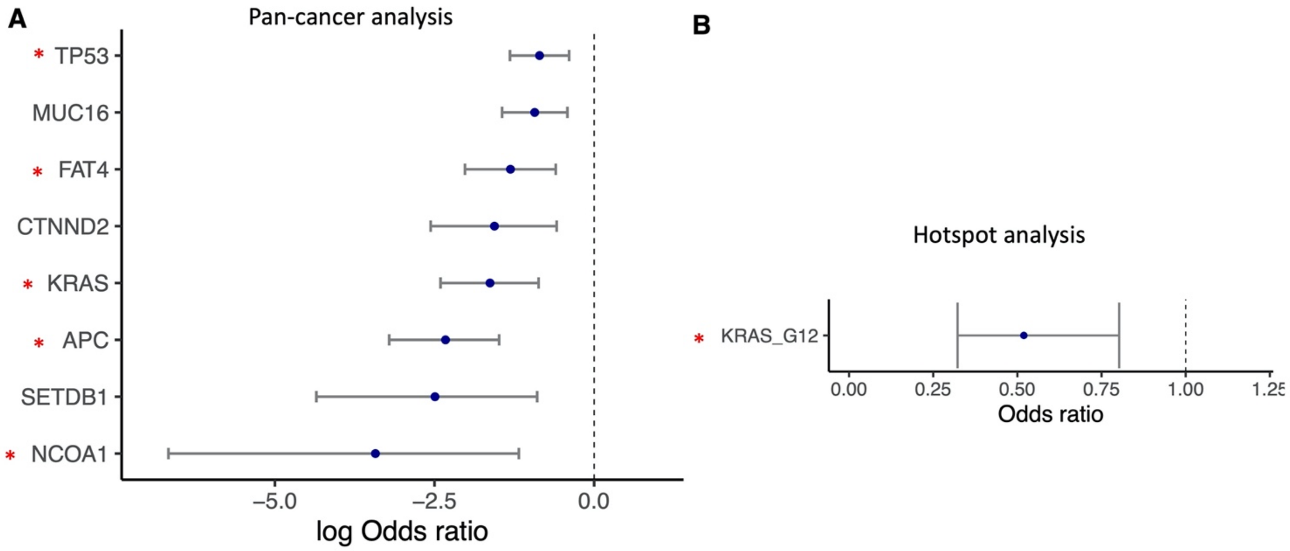


**Supplementary Figure 9. Enrichment of mutated cancer drivers and hotspots in the context of TMD in early-stage cancers. (A)** Genes with depleted mutations in early-stage cancers with TMD. **(B)** Mutational hotspot depleted in TMD in early-stage cancers. Blue circles represent odds ratios and vertical lines represent confidence intervals for each of the individual Fisher's exact tests. Red stars indicate genes which were also found to be significant in the cross-stage analysis.

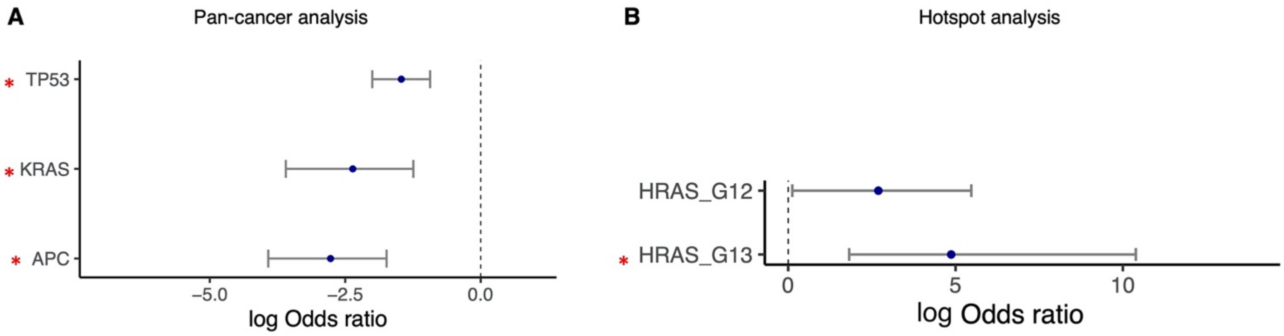


**Supplementary Figure 10. Enrichment of mutated cancer drivers and hotspots in the context of TMD in late-stage cancers. (A)** Genes with depleted mutations in late-stage cancers with TMD. **(B)** Mutational hotspot enriched in TMD in late-stage cancers. Blue circles represent odds ratios and vertical lines represent confidence intervals for each of the individual Fisher's exact tests. Red stars indicate genes which were also found to be significant in the cross-stage analysis.

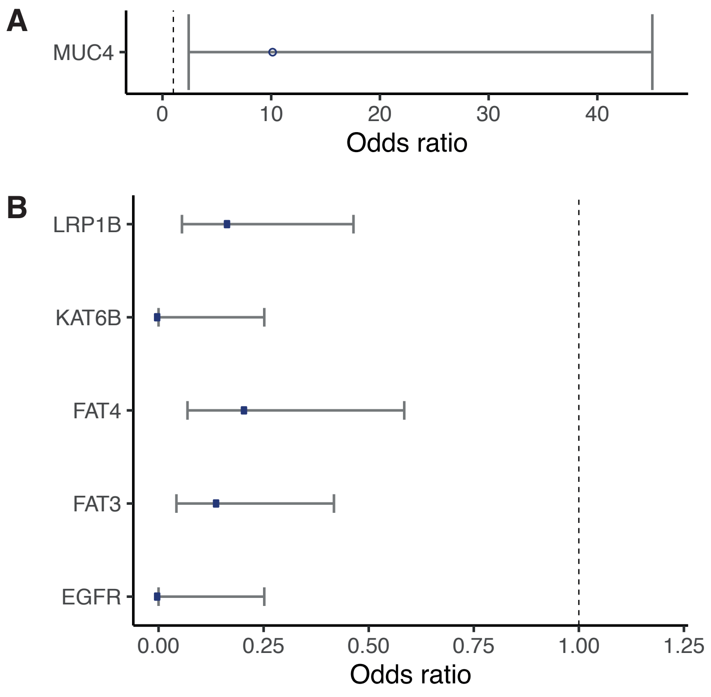


**Supplementary Figure 11. Tissue specific genomic drivers linked with tumour mass dormancy.** Genes with significant enrichment or depletion of mutations within high tumour mass dormancy samples in the colon **(A)** and stomach **(B)** adenocarcinoma TCGA studies are displayed. Blue circles represent odds ratios and vertical lines represent confidence intervals for each of the individual Fisher's exact tests.

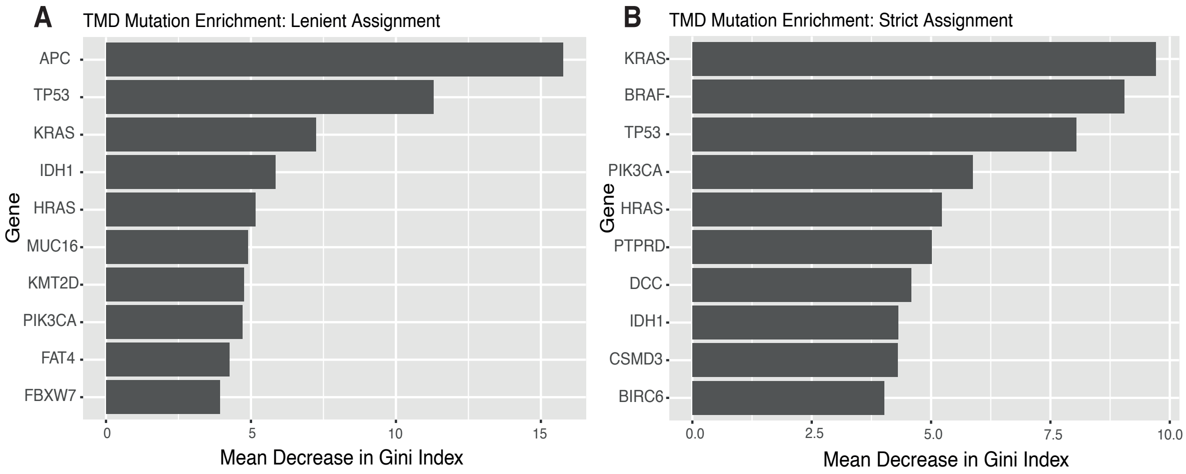


**Supplementary Figure 12. Ranking of genes contributing to the classification of TMD and non-TMD samples based on their mutation status in a random forest model.** The top 10 cancer drivers that are differentially mutated between the TMD and non-TMD samples are ranked by their importance in the classification. Mutational status is determined by **(a)** lenient and **(b)** strict assignment where, in the latter case, mutations must be labelled as 'damaging' or 'deleterious'.

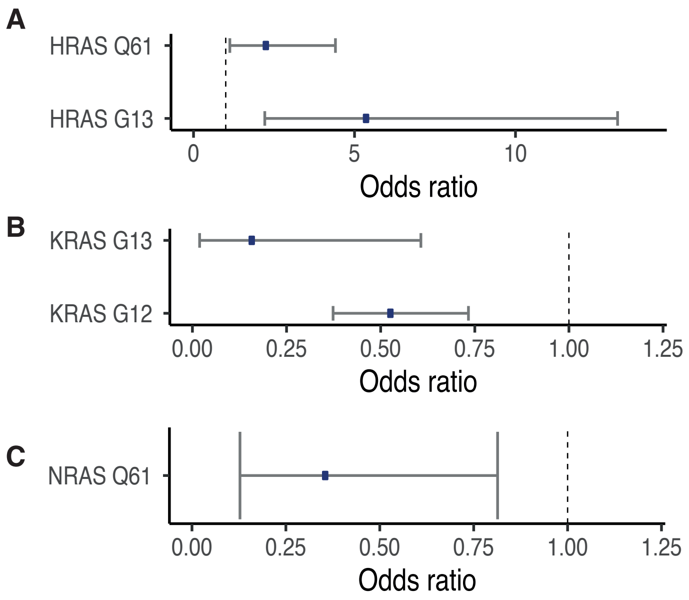


**Supplementary Figure 13. Oncogenic Ras mutations linked with tumour mass dormancy.** Hotspot mutations within the **(A)** HRAS, **(B)** KRAS and **(C)** NRAS oncogenes that are significantly enriched or depleted in samples with high TMD across the TCGA solid primary tumour samples. Blue circles represent odds ratios and vertical lines represent confidence intervals for each of the individual Fisher's exact tests.

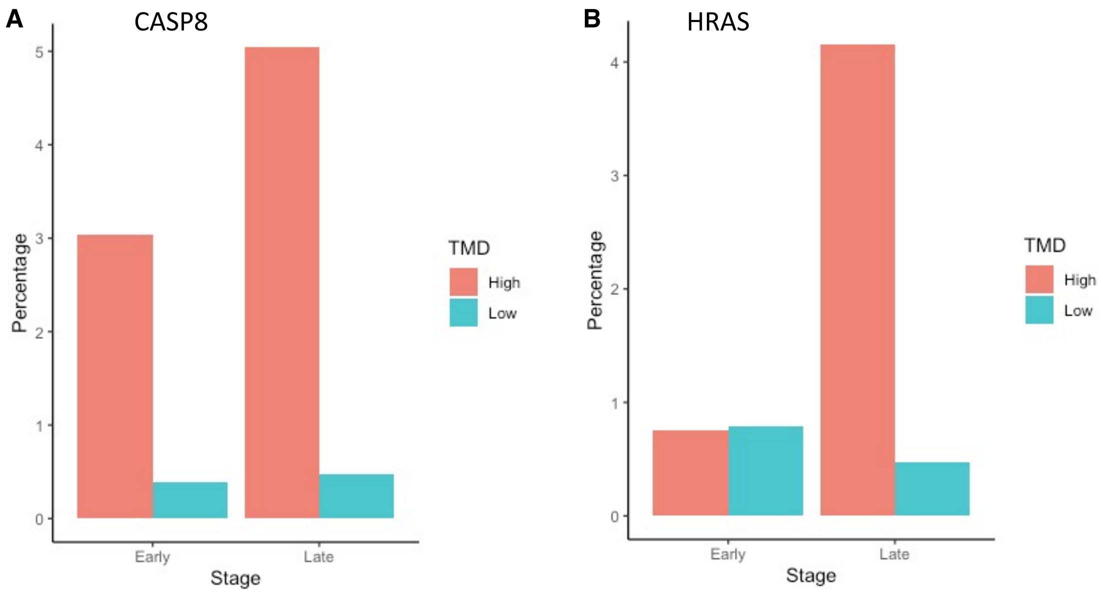


**Supplementary Figure 14. Frequency of mutations in positively selected genes CASP8 and HRAS between early- and late-stage cancers. (A)** CASP8 mutations are similarly increased in frequency in cancers with high TMD in early- and late-stage cancers. **(B)** HRAS mutations appear specifically increased in cancers with high TMD in late-stage tumours only.

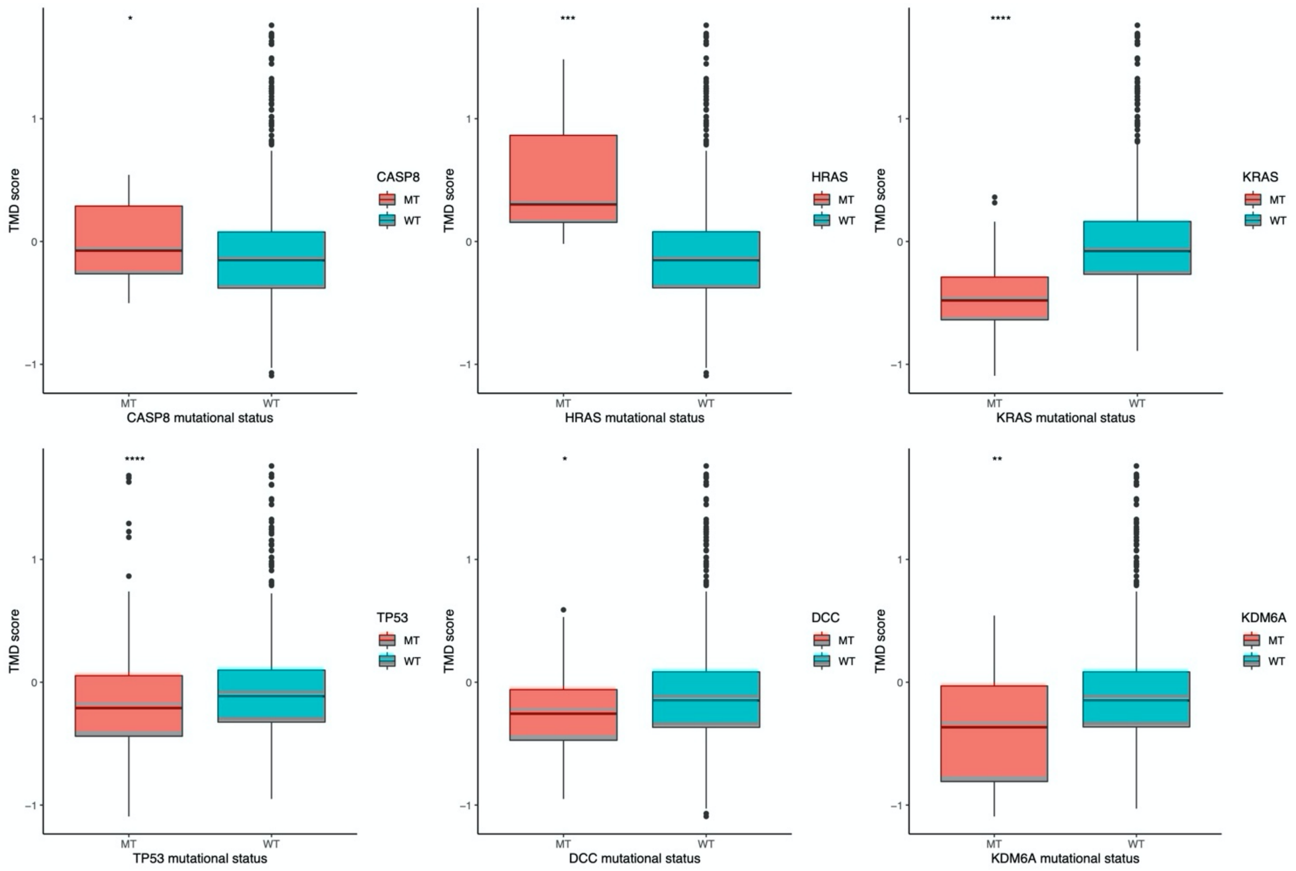


**Supplementary Figure 15. Pan-cancer frequency of mutations in genes linked with TMD in the ICGC dataset.** The TMD score is compared between tumours harbouring nonsynonymous mutations in a specific gene (red) and those without mutation in the respective gene (blue). **** p<0.0001; *** p<0.001; ** p<0.01; * p<0.05.

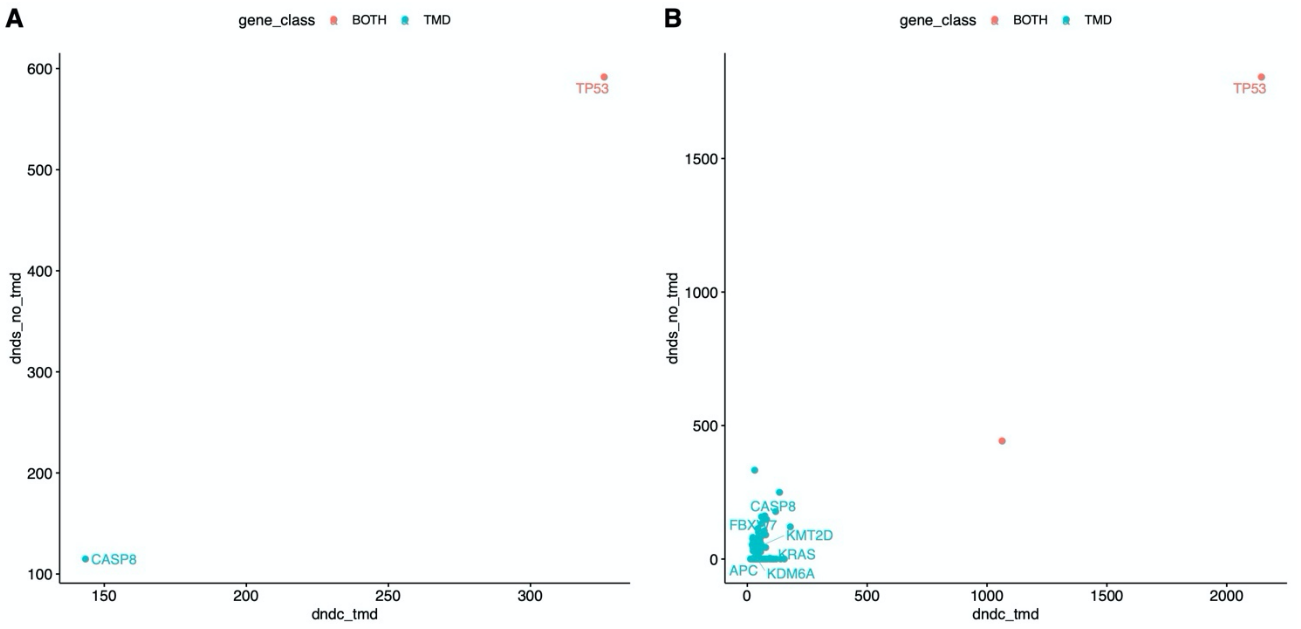


**Supplementary Figure 16. Validation of genes under positive selection linked with TMD in independent datasets. (A)** Validation in oral carcinomas (ORCA-IN cohort) from ICGC. **(B)** Validation in the METABRIC breast cancer cohort. Genes showing signals of positive selection in samples with high TMD are highlighted in blue, and across both groups (high/low TMD) in red.

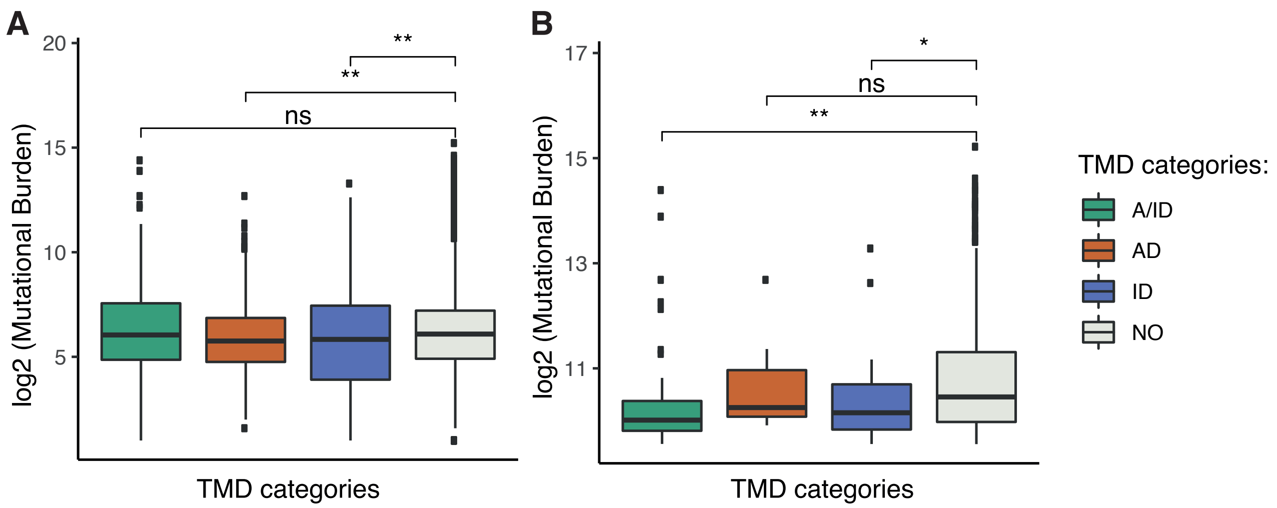


**Supplementary Figure 17. Tumour mutational burden across TMD subtypes.** Tumour mutational burden is compared between samples with angiogenic dormancy (AD), immunological dormancy (ID), both angiogenic and immunological (A/ID) and samples without TMD (NO) before **(A)** and after **(B)** excluding samples with less than 750 mutations.

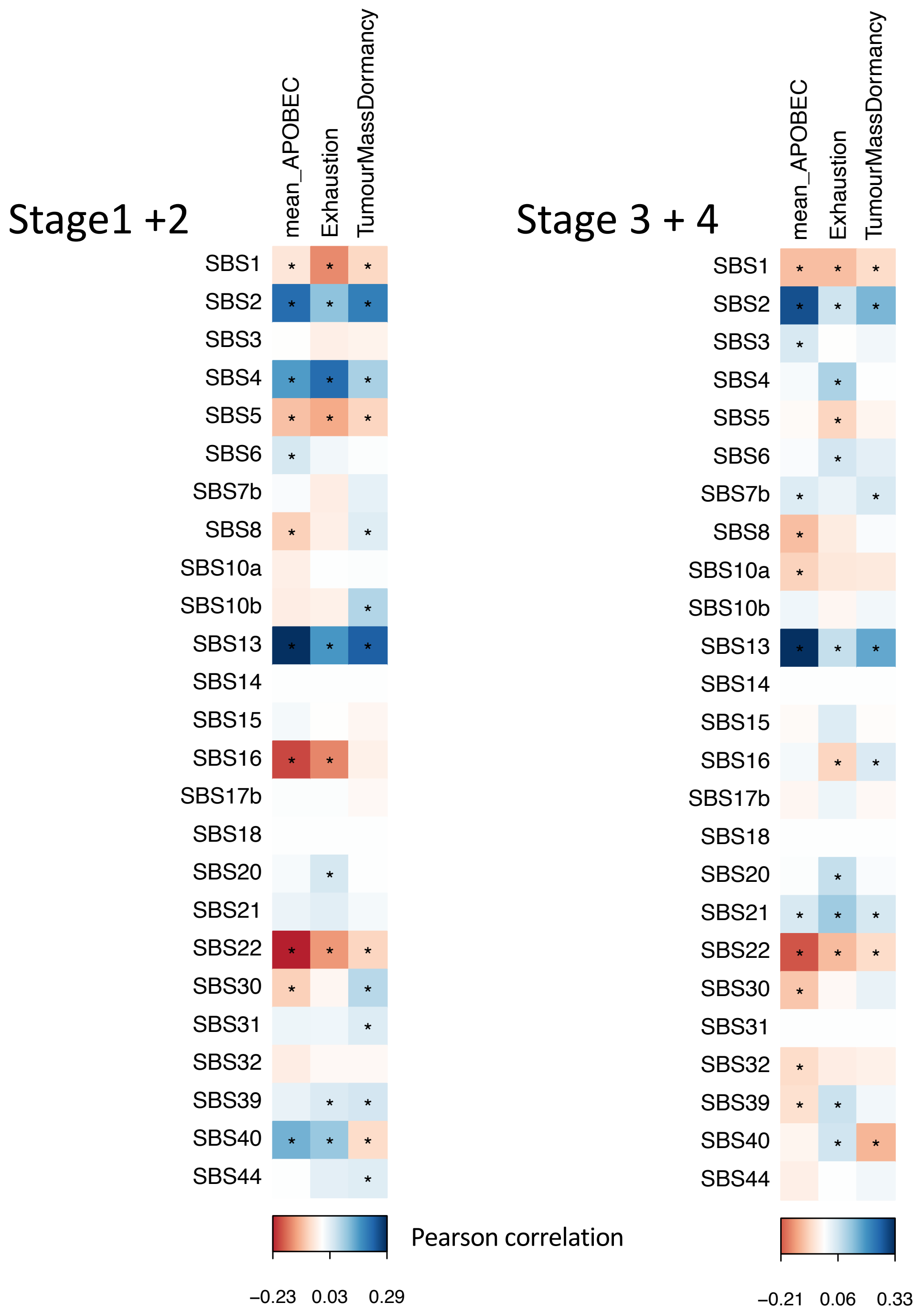


**Supplementary Figure 18. Pan-cancer correlations between mutational signatures and tumour mass dormancy, by cancer stage.** Matrices depicting the Person correlation between mutational signatures and the TMD, exhaustion and APOBEC programmes are shown for stage 1 and 2 cancers (left) and stage 3 and 4 cancers (right). Statistically significant correlations ($p<0.05$) are highlighted with an asterisk.

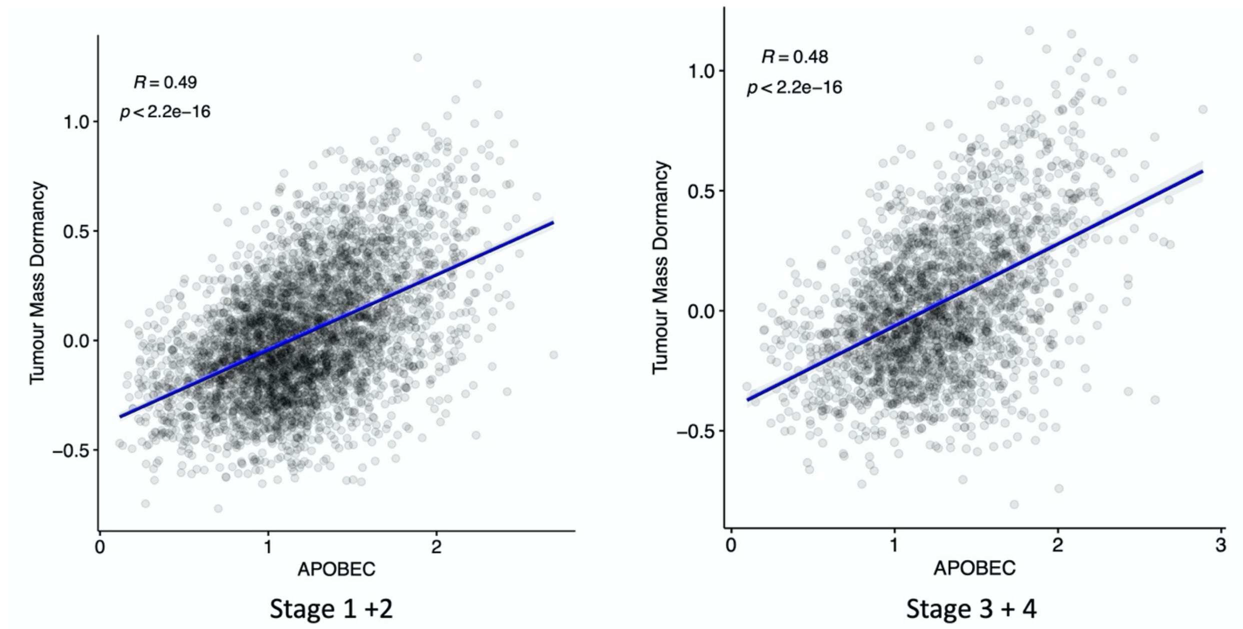


**Supplementary Figure 19. Pan-cancer correlations between TMD and APOBEC, by cancer stage.** Left: correlations in early-stage cancer; right: correlations in late-stage cancers. The x and y axes depict the per-sample expression score for the respective programmes.

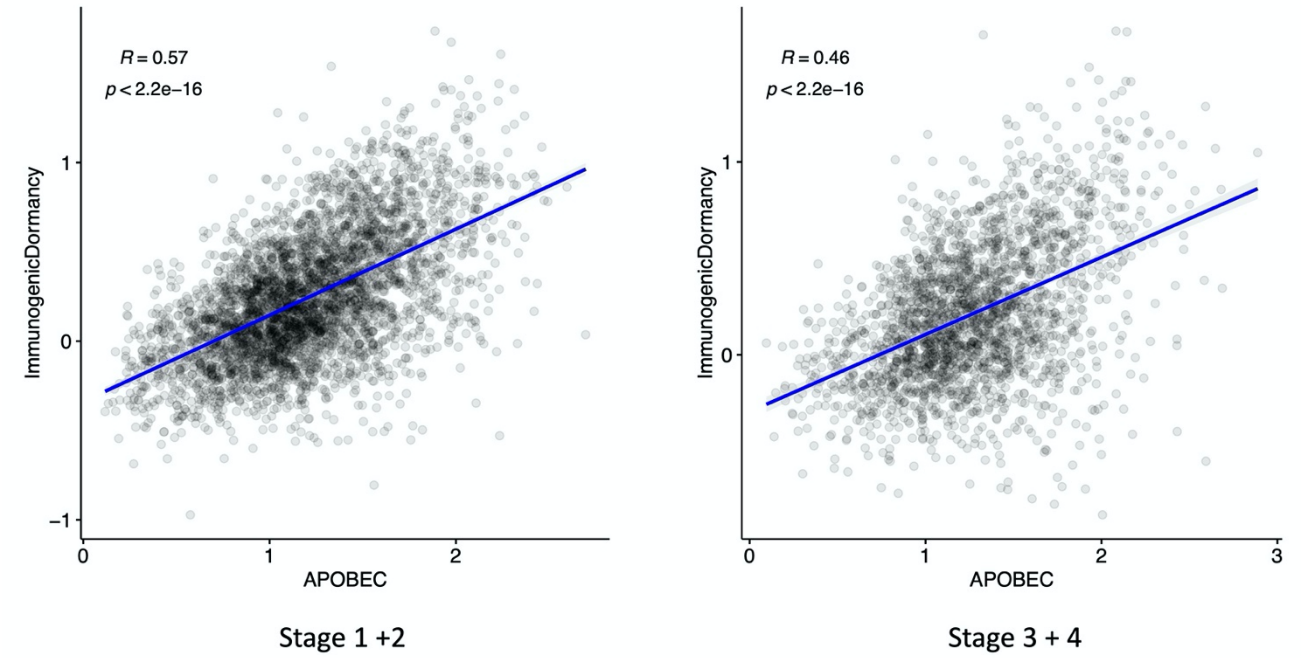


**Supplementary Figure 20. Pan-cancer correlations between immunological dormancy and APOBEC, by cancer stage.** Left: correlations in early-stage cancer; right: correlations in late-stage cancers. The x and y axes depict the per-sample expression score for the respective programmes.

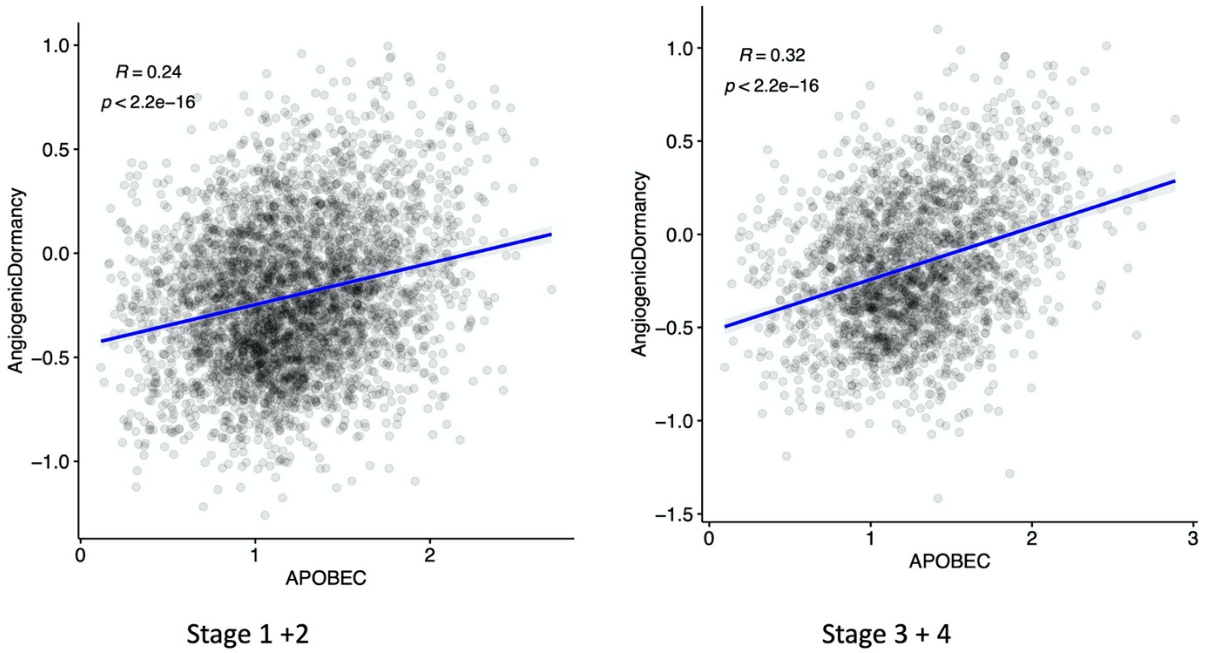


**Supplementary Figure 21. Pan-cancer correlations between angiogenic dormancy and APOBEC, by cancer stage.** Left: correlations in early-stage cancer; right: correlations in late-stage cancers. The x and y axes depict the per-sample expression score for the respective programmes.

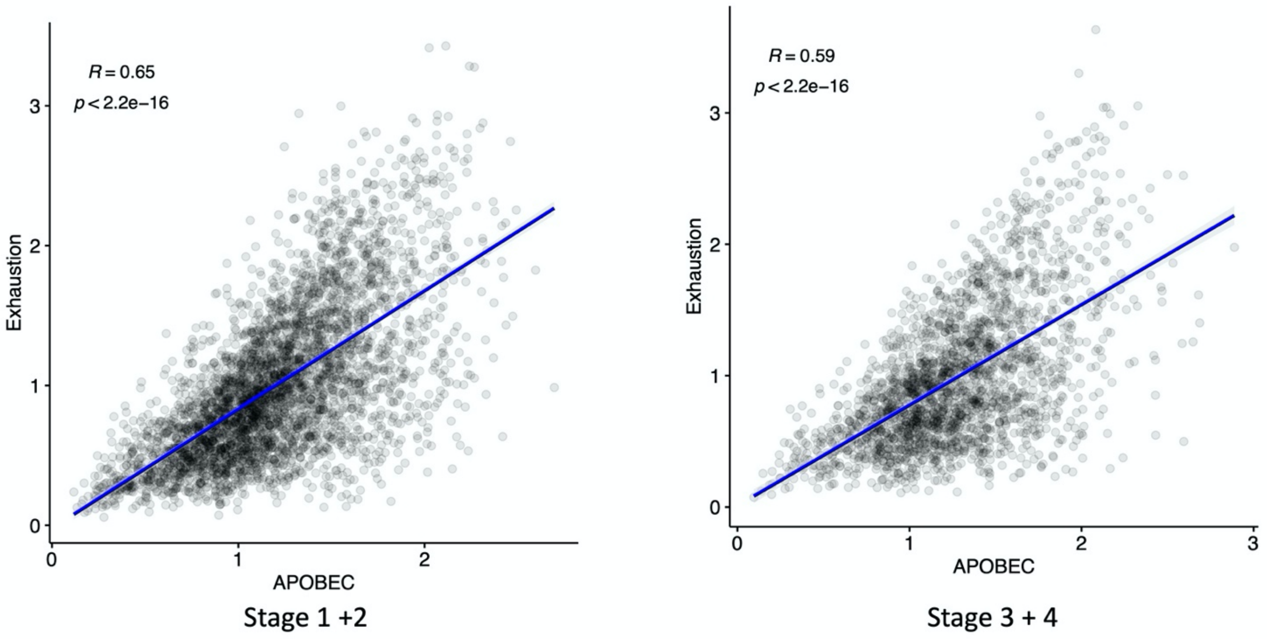


**Supplementary Figure 22. Pan-cancer correlations between exhaustion and APOBEC, by cancer stage.** Left: correlations in early-stage cancer; right: correlations in late-stage cancers. The x and y axes depict the per-sample expression score for the respective programmes.

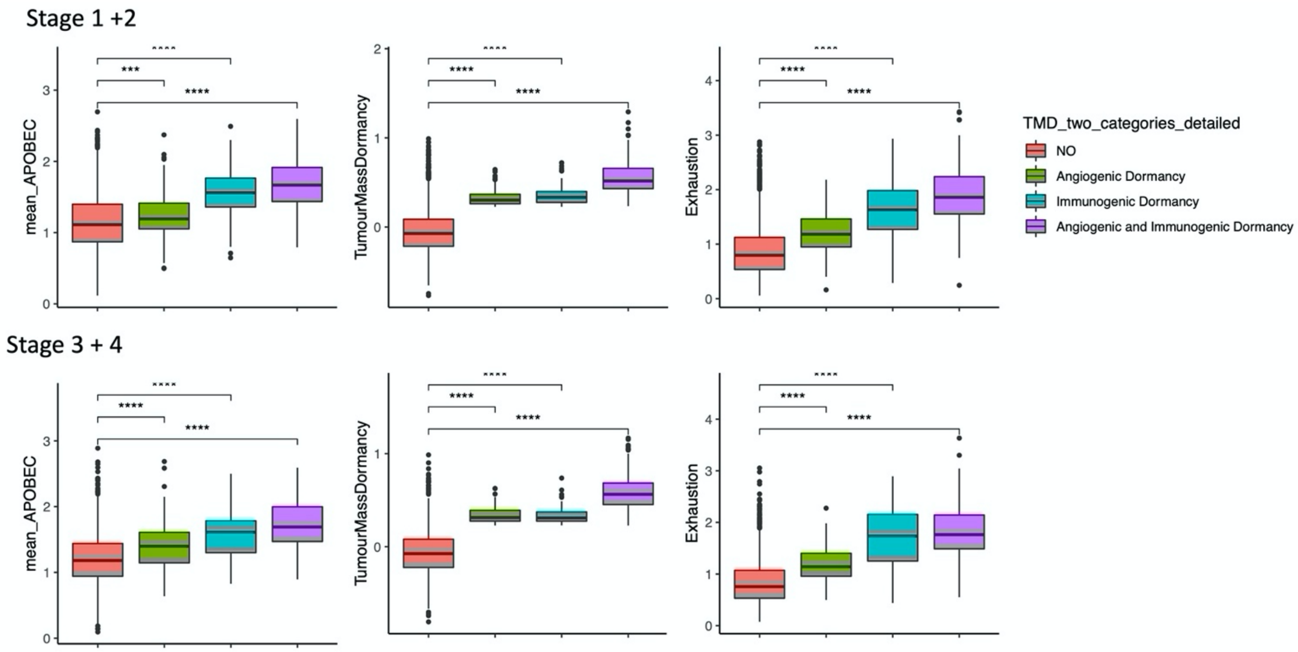


**Supplementary Figure 23. Tumour mass dormancy and linked progammes by clinical stage.** TMD, APOBEC and exhaustion programme scores are compared between samples with angiogenic dormancy, immunological dormancy, both angiogenic and immunological dormancy and expanding tumours without evidence of TMD (NO) in early (top panel) and late stage cancer (bottom panel); **** p<0.0001.

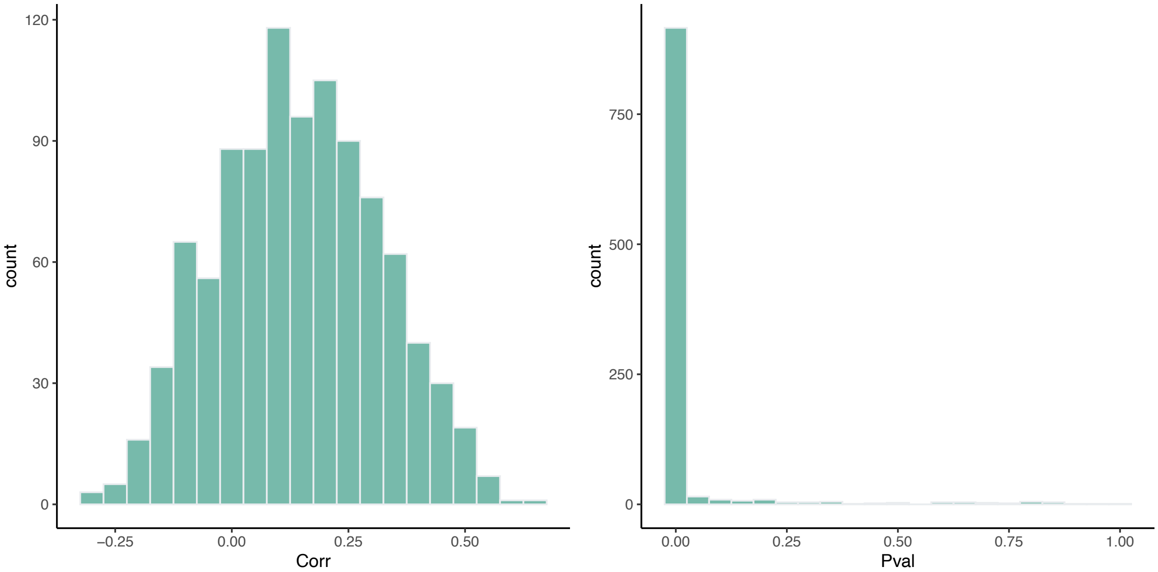


**Supplementary Figure 24. Correlations between APOBEC activity and the average expression of 35 randomly selected genes (matching the size of the TMD signature).** Left: Pearson correlation coefficient distribution across 1000 repeats (mean correlation = 0.144; 89% highest posterior density interval: [-0.12, 0.44]). Right: Distribution of corresponding p-values across 1000 iterations. The small p-values are hypothesized to reflect the large size of the dataset.

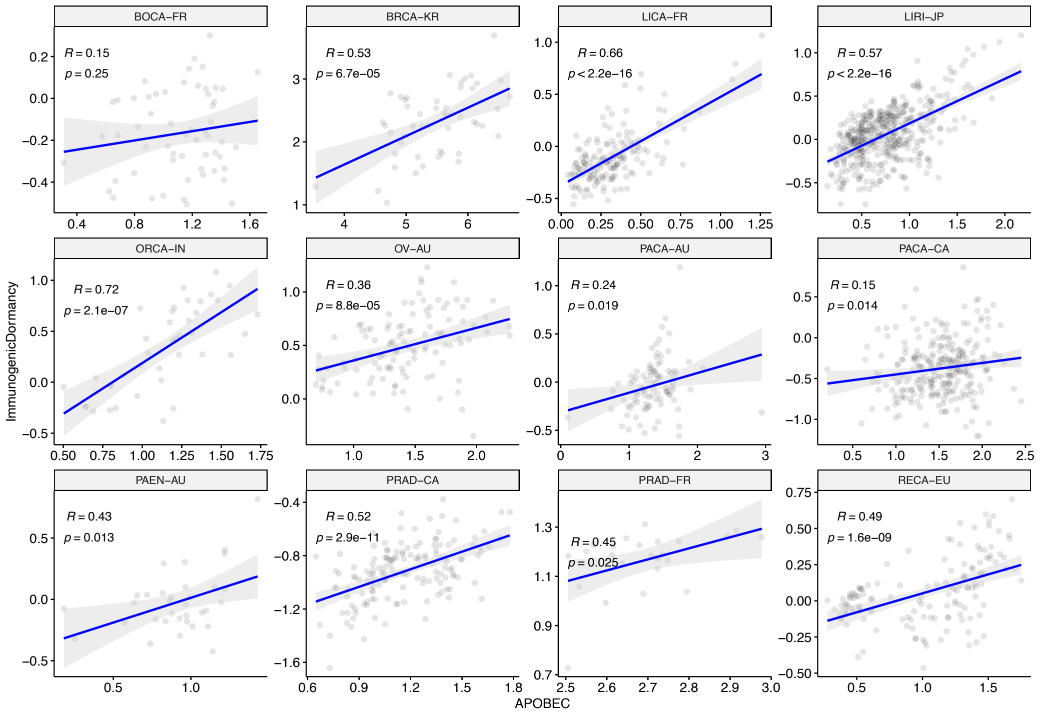


**Supplementary Figure 25. Correlations between immunological dormancy and APOBEC activity in multiple cancers from ICGC.** The x and y axes depict the per-sample expression score for the respective programmes.

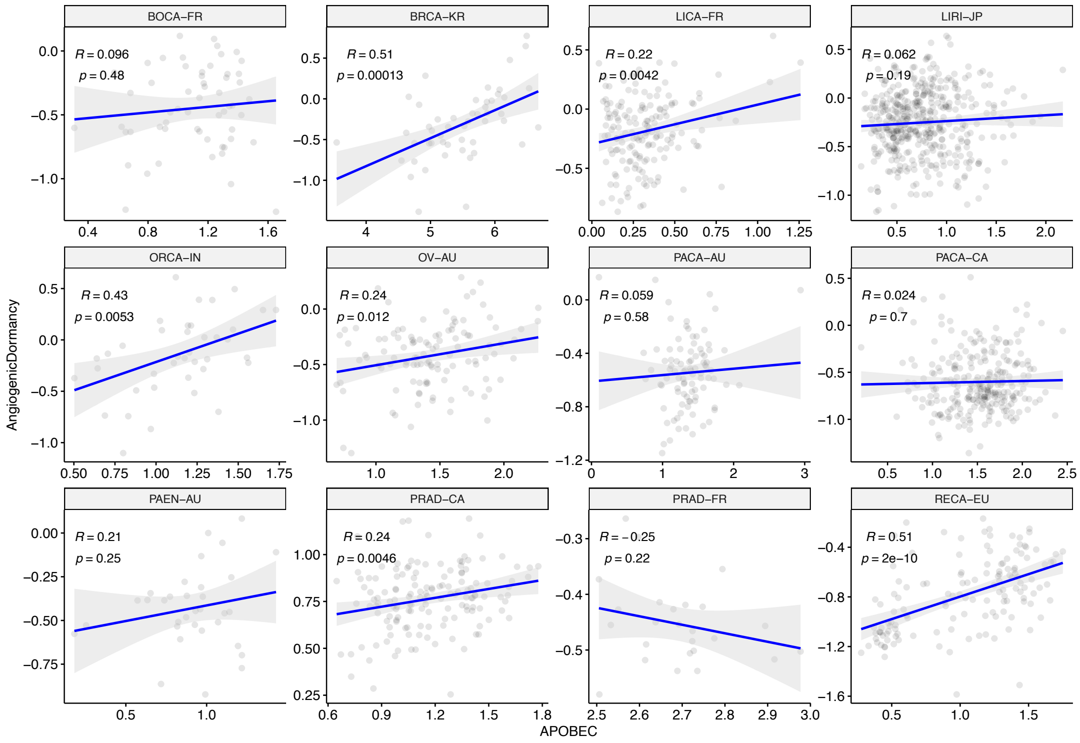


**Supplementary Figure 26. Correlations between angiogenic dormancy and APOBEC activity in multiple cancers from ICGC.** The x and y axes depict the per-sample expression score for the respective programmes.

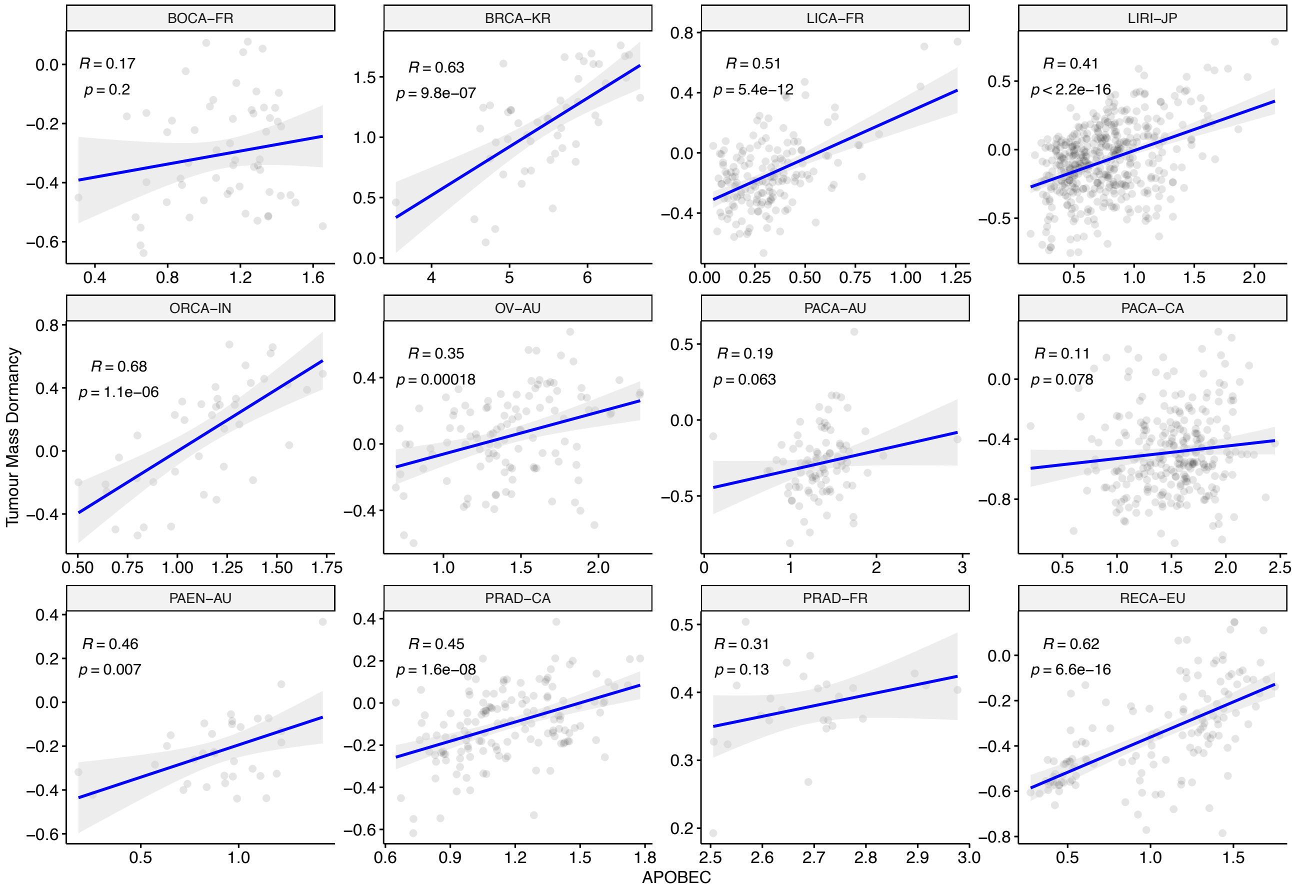


**Supplementary Figure 27. Correlations between TMD and APOBEC activity in multiple cancers from ICGC.** The x and y axes depict the per-sample expression score for the respective programmes.

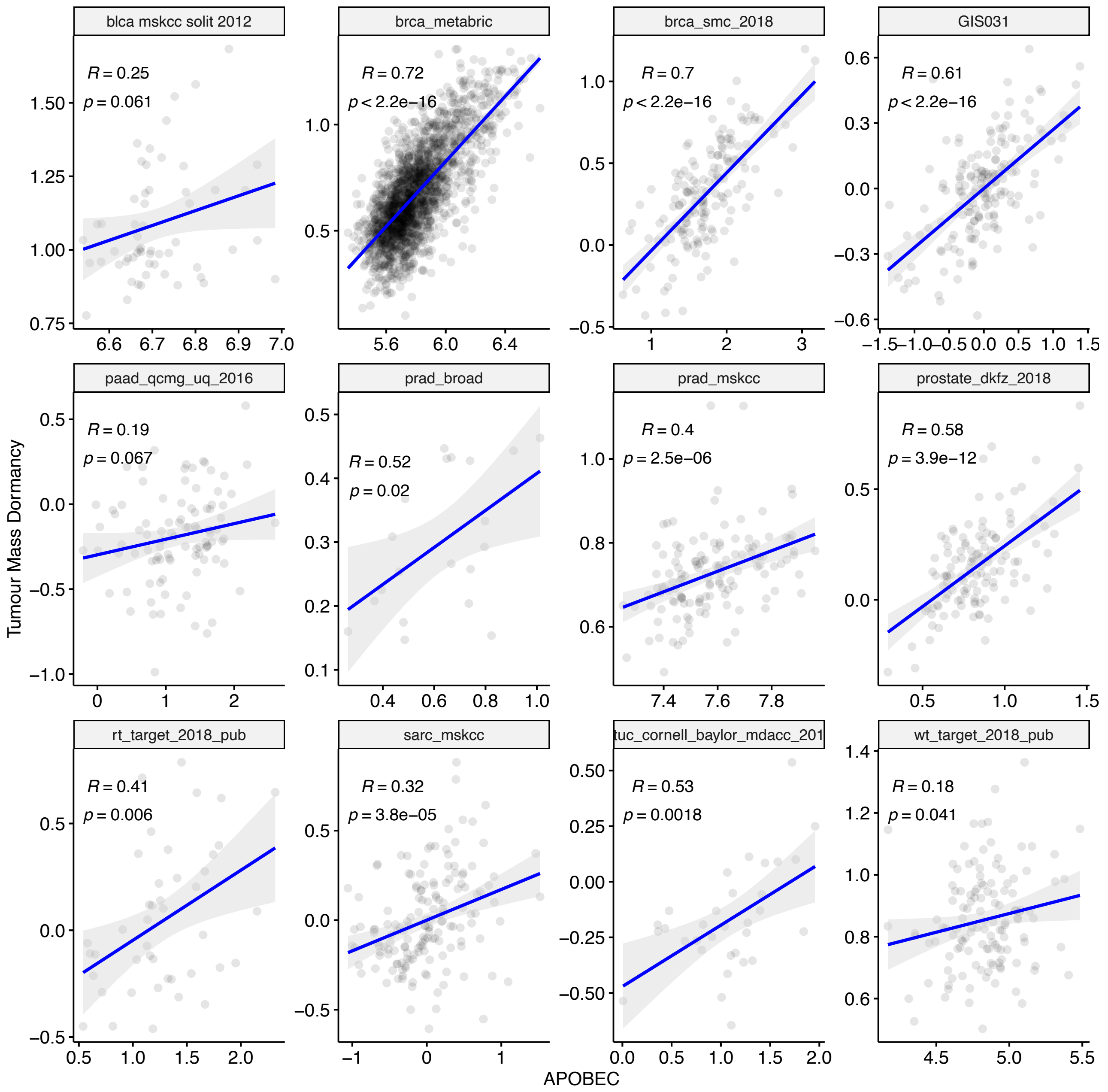


**Supplementary Figure 28. Correlations between TMD and APOBEC activity in multiple studies from cBioPortal.** The x and y axes depict the per-sample expression score for the respective programmes. blca_mskcc_solit_2012 = Bladder Cancer (MSKCC, J Clin Onco 2013); brca_metabric = Breast Cancer (METABRIC, Nature 2012 & Nat Commun 2016); brca_smc_2018 = Breast Cancer (SMC 2018); GIS031 = Lung adenocarcinoma (GIS, Nat Genet 2019); paad_qcmg_uq_2016 = Pancreatic Adenocarcinoma (QCMG, Nature 2016); prad_broad = Prostate Adenocarinoma (Broad/Cornell, Nat Genet 2012); prad_mskcc = Prostate Adenocarcinoma (MSKCC, Cancer Cell 2010); prostate_dkfz_2018 = Prostate Cancer (DKFZ, Cancer Cell 2018); rt_target_2018_pub = Pediatric Rhabdoid Tumor (TARGET, 2018); sarc_mskcc = Sarcoma (MSKCC/Broad, Nat Genet 2010); utuc_cornell_baylor_mdacc_2019 = Upper Tract Urothelial Carcinoma (Cornell/Naylor/MDACC, Nat Commun 2019); wt_target_2018_pub = Pediatric Wilm's Tumor (TARGET, 2018).

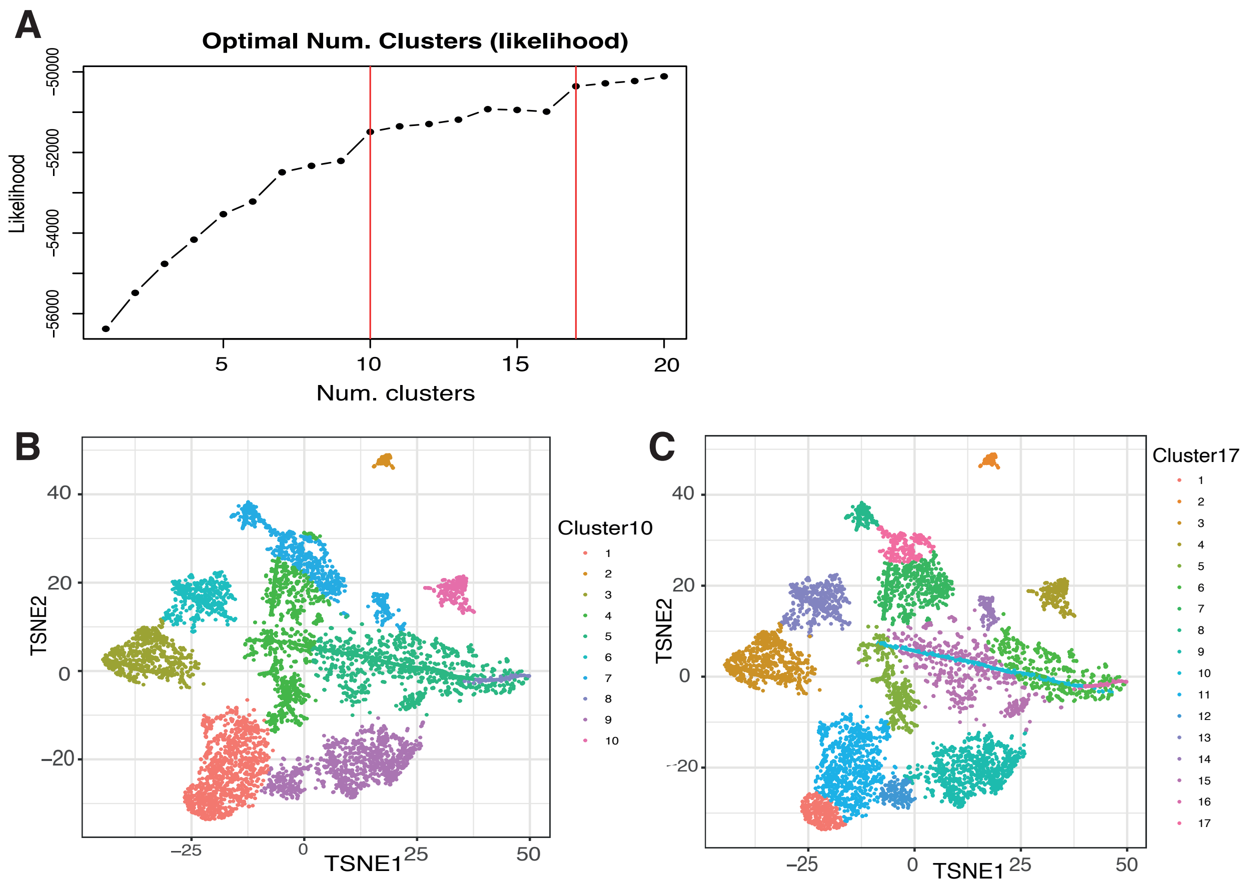


**Supplementary Figure 29. Expectation-Maximization clustering of 6,410 mutational signature profiles. (A)** The decrease in absolute likelihood associated with the determined clusters highlighted 10 and 17 clusters as the optimum number for the cohort. **(B)** tSNE dimensionality reduction of mutational signature profiles across the cohort, separated into 10 clusters. **(C)** tSNE dimensionality reduction of mutational signature profiles across the cohort, separated into 17 clusters.

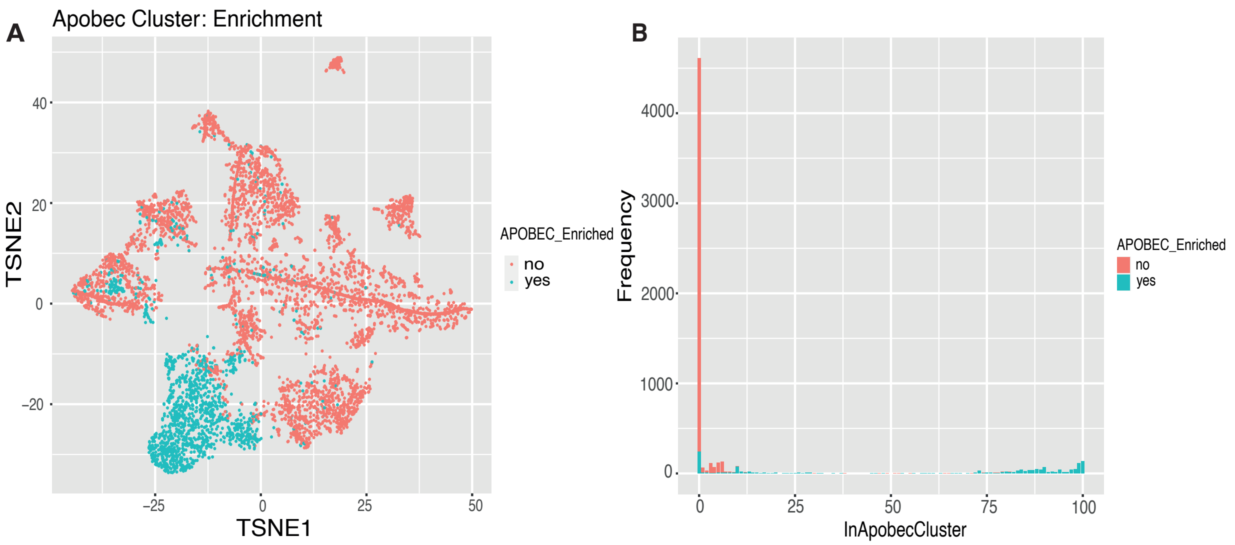


**Supplementary Figure 30. APOBEC signature cluster assignment matches strongly with APOBEC enrichment scores. (A)** A tSNE dimensionality reduction plot based on the mutational signature profiles of 6,410 TCGA primary tumour samples, coloured by APOBEC mutagenesis enrichment as determined by Roberts et al (Nat Genetics, 2013). **(B)** Barplot displaying, for each sample, how often it is assigned to the APOBEC enrichment cluster across 100 iterations of tSNE and expectation-maximization clustering. Samples assigned to this cluster on more than 50/100 occasions were labelled as 'APOBEC Enriched'.

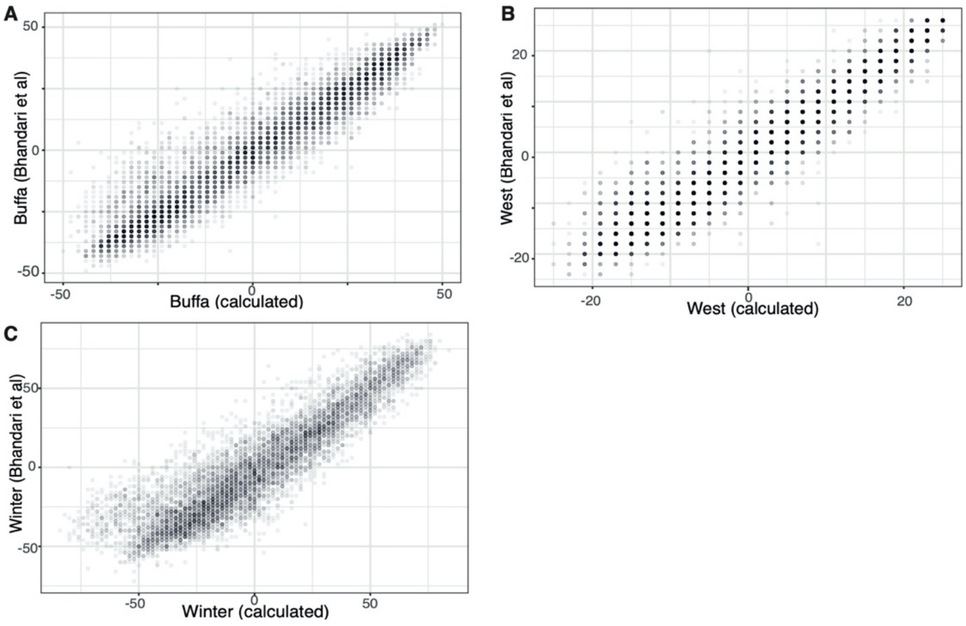


**Supplementary Figure 31. Validation of hypoxia scores.** Comparison of hypoxia signature scores with those calculated by Bhandari et al (Nat Genetics, 2019) in 7,271 overlapping samples. Hypoxia score calculation procedure is verified using the **(A)** Buffa, **(B)** West and **(C)** Winter signatures independently. Differences are attributable to the removal of genes from the signatures if they were also included within the TMD and exhaustion programme signatures, and to the fact that the scores were calculated using different cohorts.

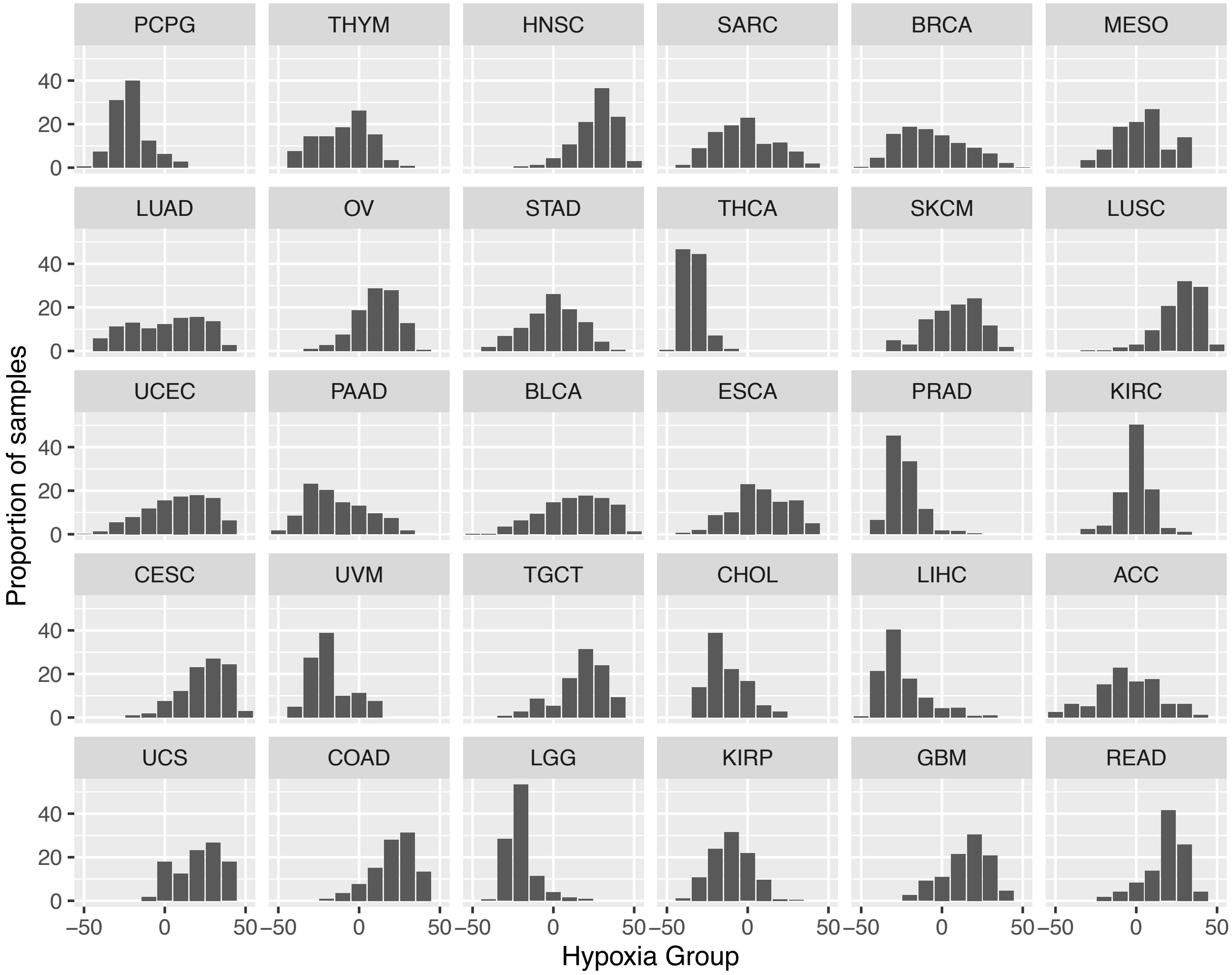


**Supplementary Figure 32. Distribution of hypoxia scores by cancer type.** Hypoxia scores were binned by rounding them to the nearest 10 and are plotted as a proportion of samples of each specific cancer type. Cancer types are ordered by descending frequency of TMD samples, so as to match Figure 5C. Consequently, KICH was not plotted because it lacked TMD samples.

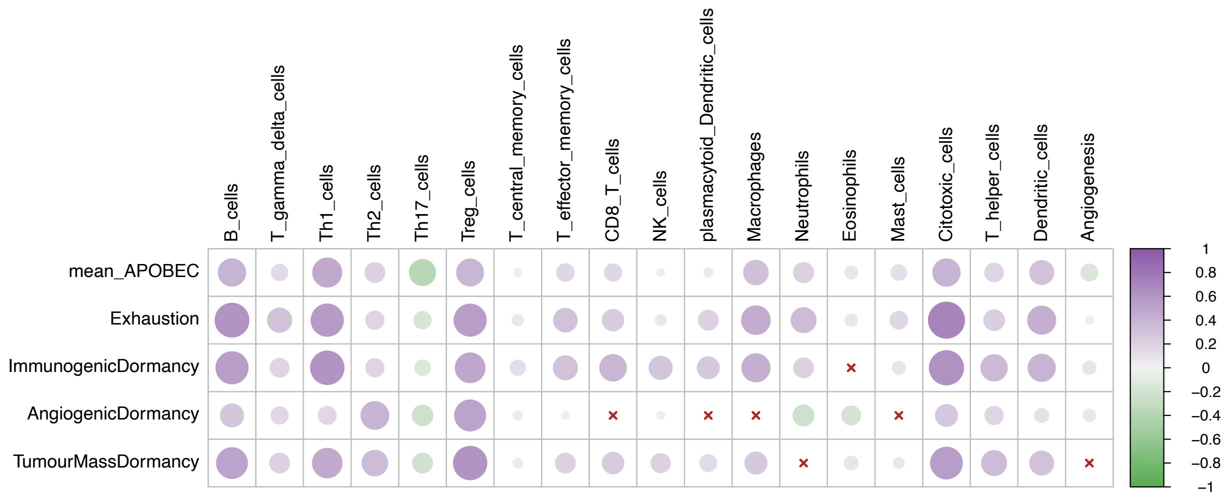


**Supplementary Figure 33. Tumour microenvironment activity correlates with the TMD programme in early-stage cancers only.** Correlation between cell infiltration estimates and the APOBEC, exhaustion and dormancy programme scores across the TCGA primary tumor samples. The colour gradient reflects the Pearson correlation coefficient.

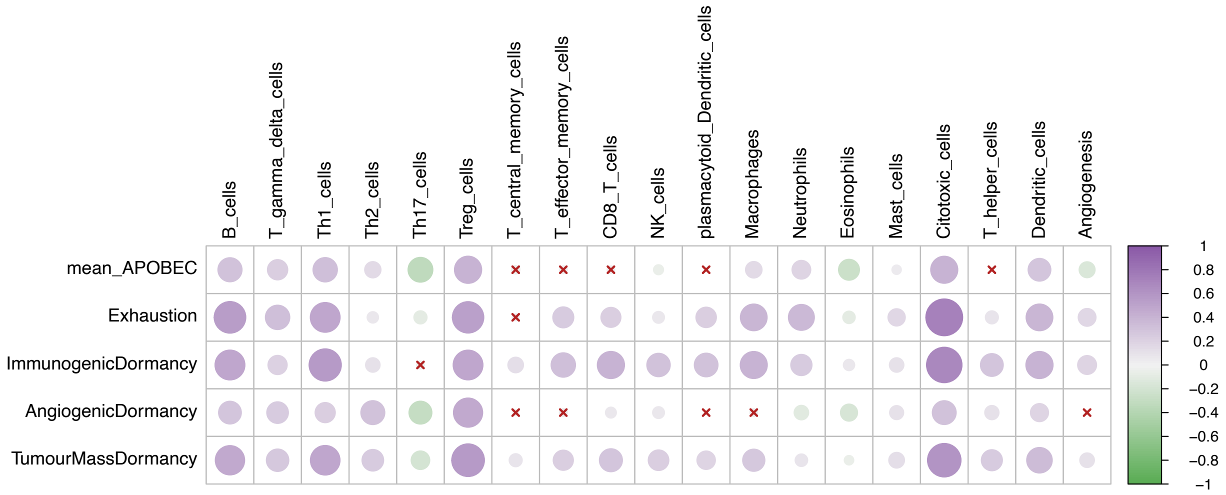


**Supplementary Figure 34. Tumour microenvironment activity correlates with the TMD programme in late-stage cancers only.** Correlation between cell infiltration estimates and the APOBEC, exhaustion and dormancy programme scores across the TCGA primary tumor samples. The colour gradient reflects the Pearson correlation coefficient.

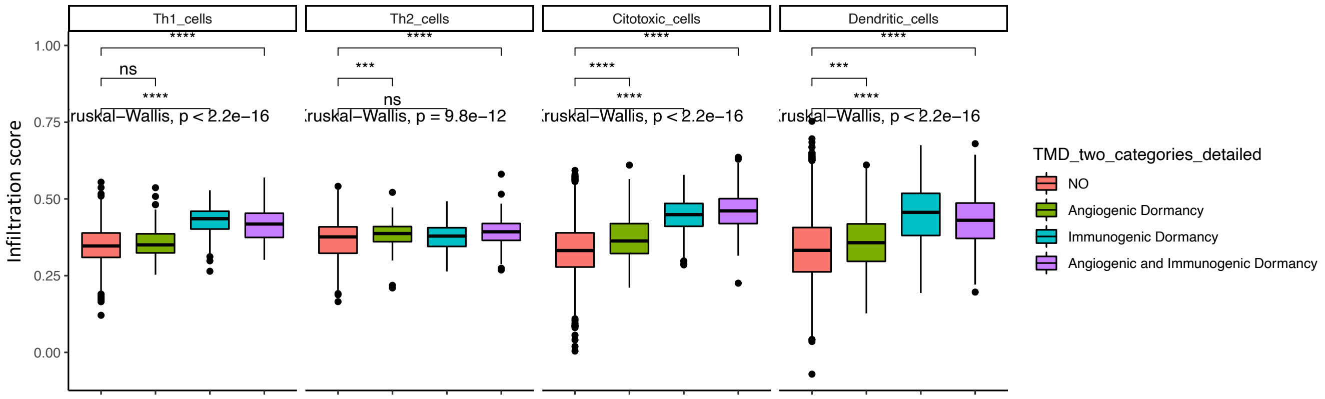


**Supplementary Figure 35. TMD categories differences in immune cell composition in early-stage cancers.** Cell infiltration scores are compared between samples with angiogenic dormancy, immunological dormancy, both angiogenic and immunological dormancy and expanding tumours without evidence of TMD (NO); **** p<0.0001.

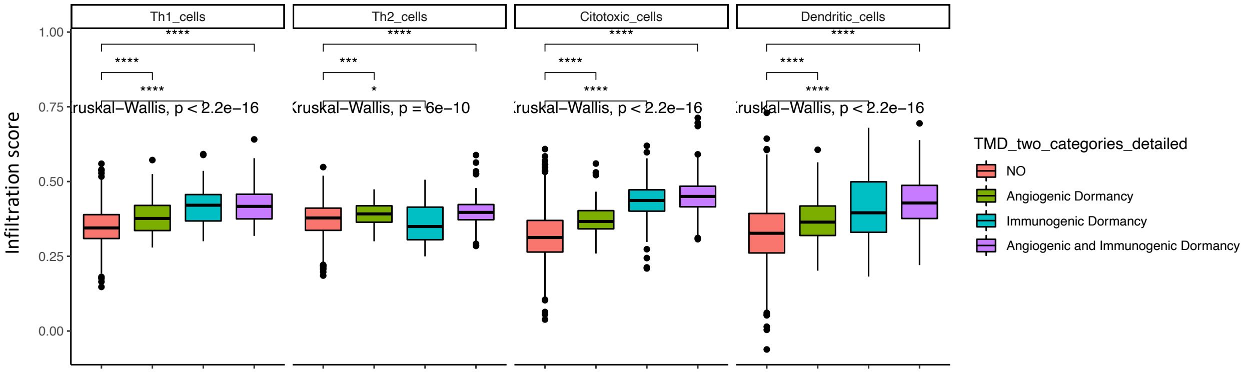


**Supplementary Figure 36. TMD categories differences in immune cell composition in late-stage cancers.** Cell infiltration scores are compared between samples with angiogenic dormancy, immunological dormancy, both angiogenic and immunological dormancy and expanding tumours without evidence of TMD (NO); **** p<0.0001.

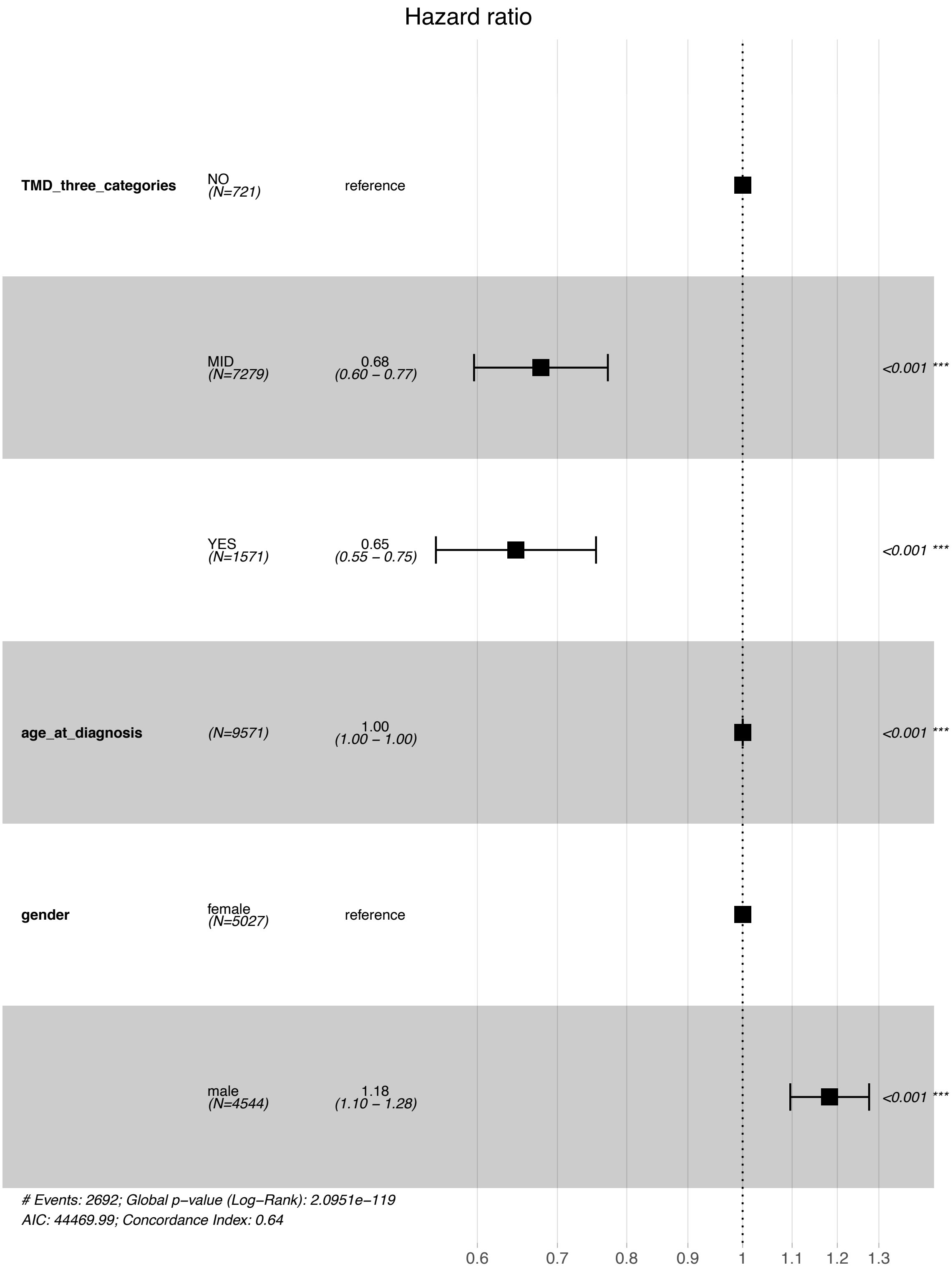


**Supplementary Figure 37. Cox proportional hazards model of overall survival modelled on TMD category while accounting for patient age and gender.** Hazard ratios are displayed.

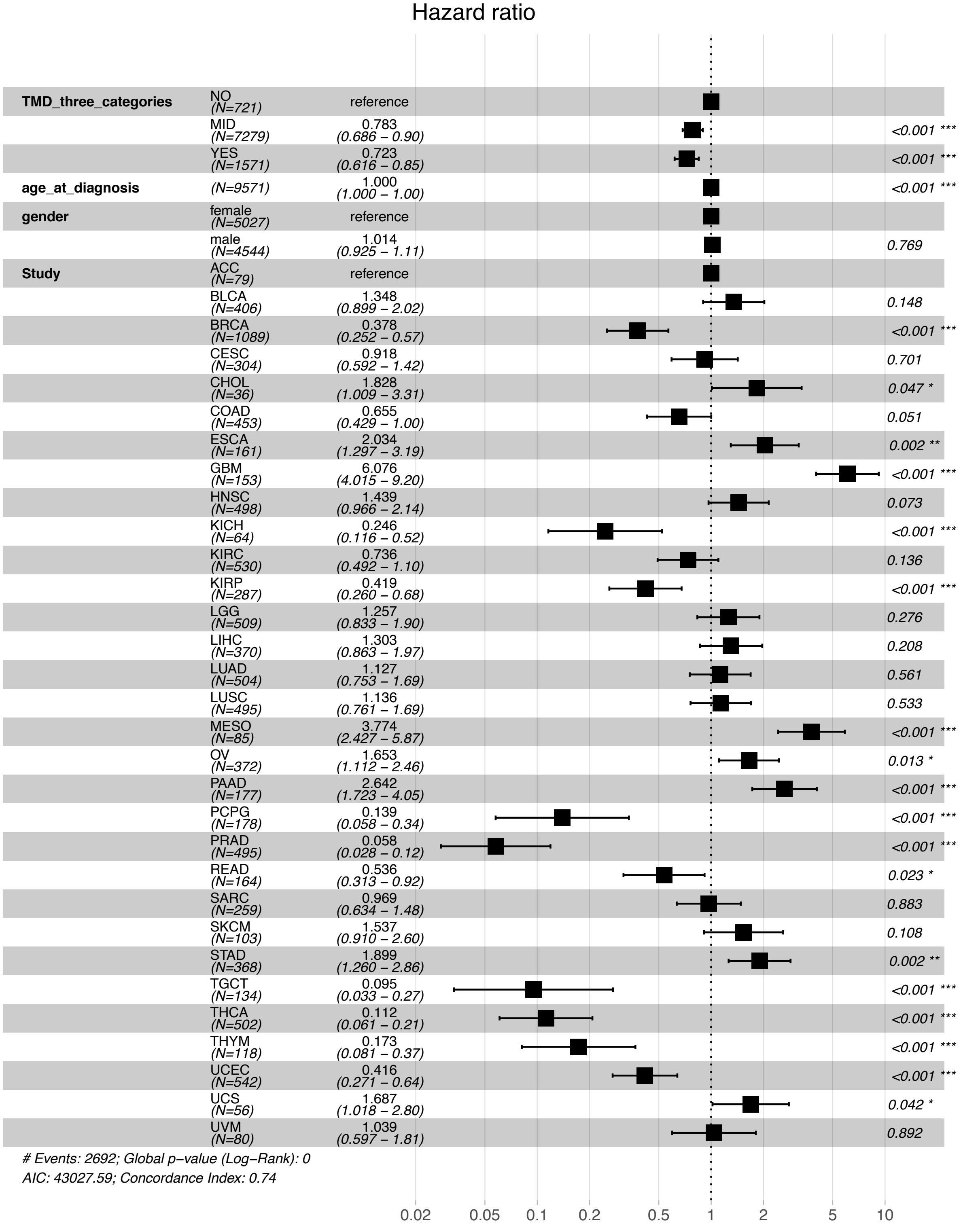


**Supplementary Figure 38. Cox proportional hazards model of overall survival modelled on TMD category while accounting for patient age, gender and cancer study.** Hazard ratios are displayed.

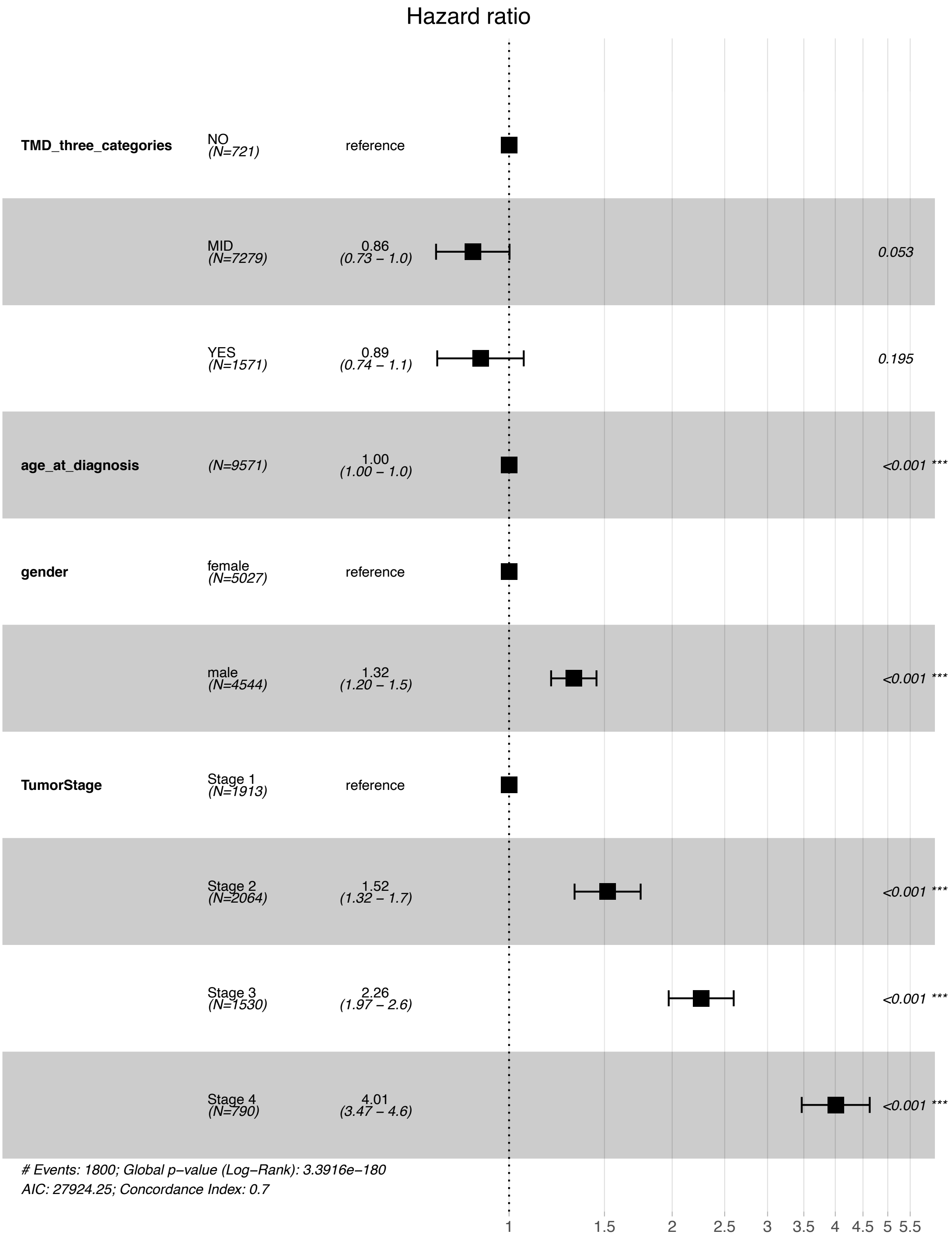


**Supplementary Figure 39. Cox proportional hazards model of overall survival modelled on TMD category while accounting for patient age, gender and cancer stage.** Hazard ratios are displayed.

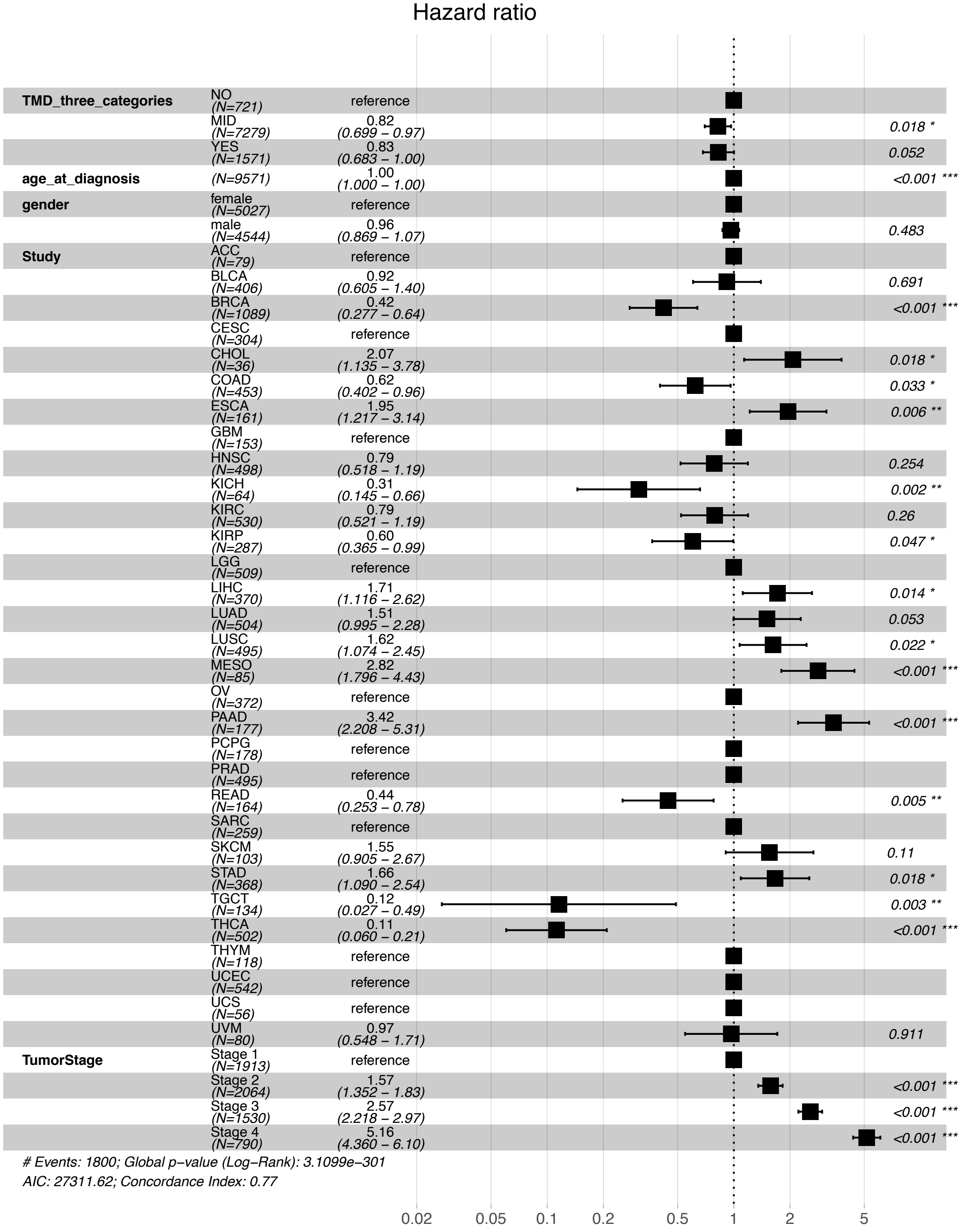


**Supplementary Figure 40. Supplementary Figure 15. Cox proportional hazards model of overall survival modelled on TMD category while accounting for patient age, gender, cancer study and stage.** Hazard ratios are displayed. Note that this model has the highest concordance index out of all the Cox models, at 0.77.

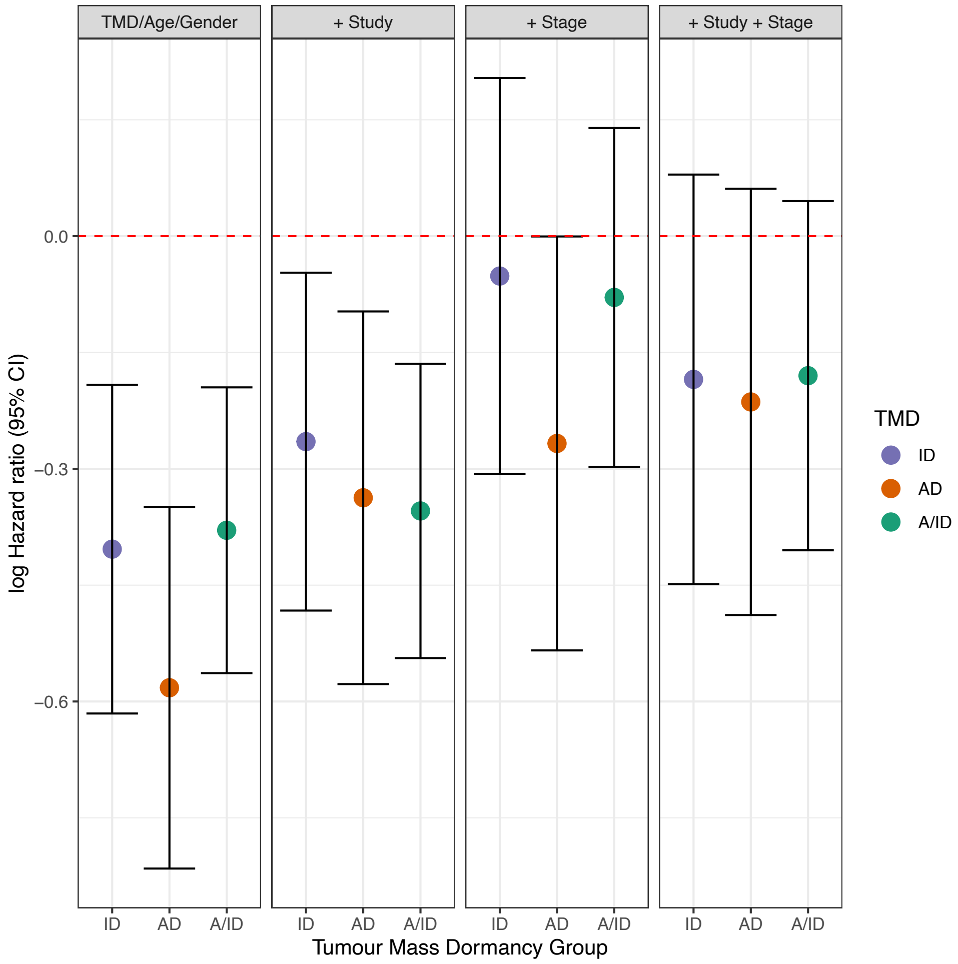


**Supplementary Figure 41. Log hazard ratios of survival for detailed dormancy categories.** Overall survival is compared to the 'NO' dormancy category, with tumour stage labelled between Stage I and Stage IV, as opposed to a binary early/late classification. AD – tumours with angiogenic dormancy; ID – tumours with immunological dormancy; A/ID – tumours with both angiogenic and immunological dormancy.

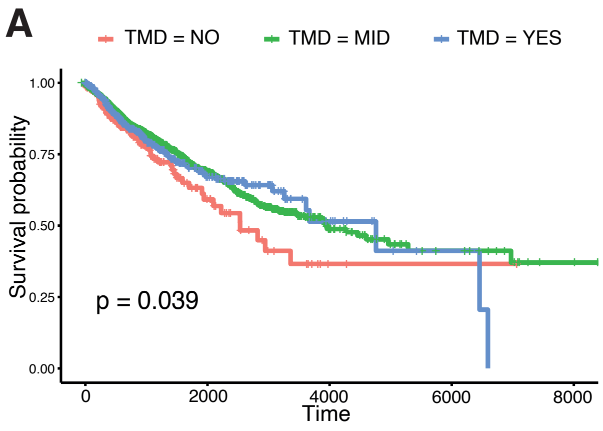


**Supplementary Figure 42. Overall survival differences by TMD category in early- and late-stage cancers.** Kaplan-Meier plots demonstrate consistently worse prognosis for tumours with no evidence of TMD (‘NO’) individually for **(A)** early-stage cancers (Stages I and II) and **(B)** late-stage cancers (Stages III and IV). ‘YES’ denotes high TMD; ‘MID’ denotes average proliferating tumours.

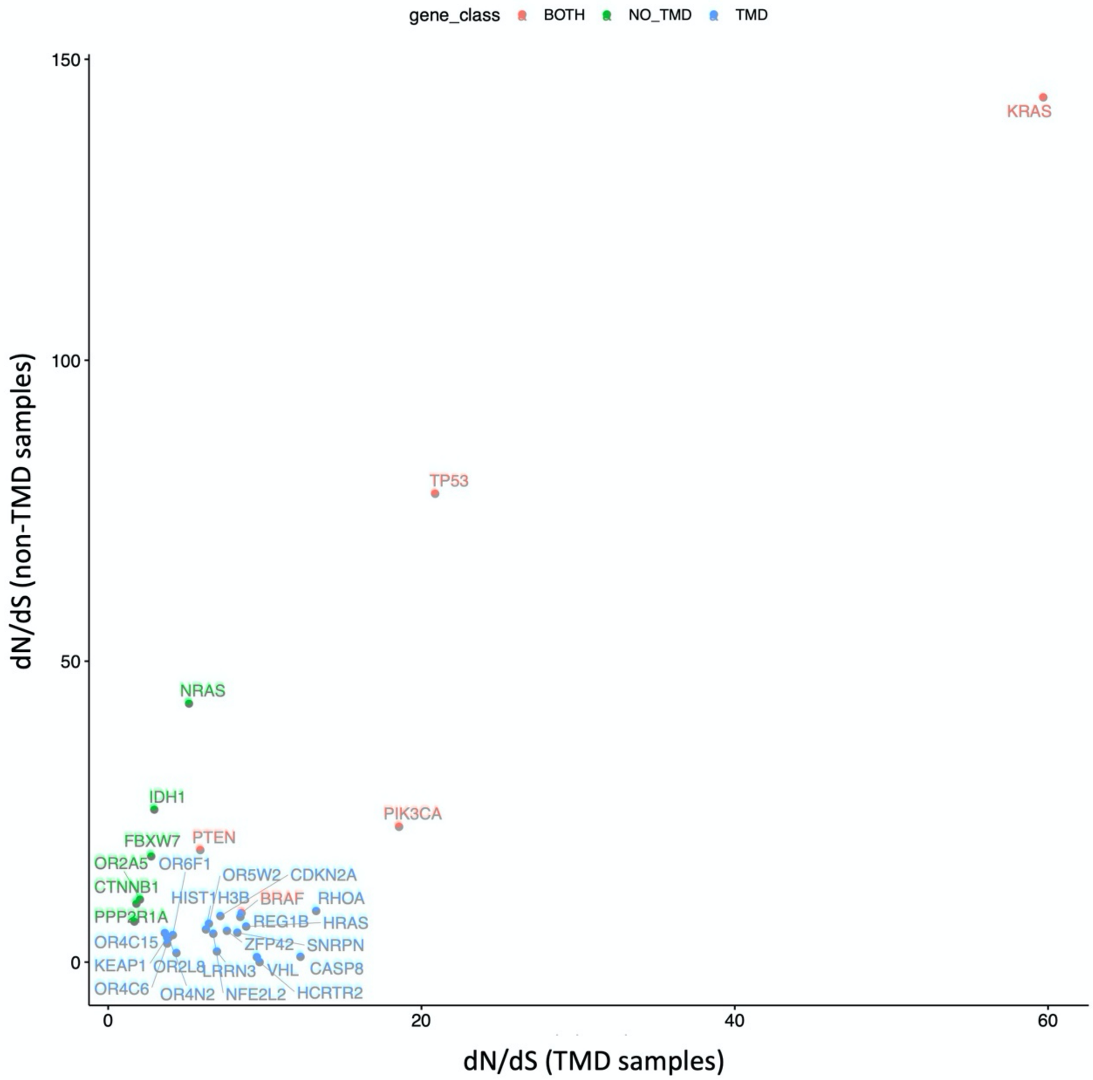


**Supplementary Figure 43. Genes under positive selection across TCGA in the context of TMD, as derived from PCA-based methodology.** Genes showing signals of positive selection in samples with high TMD are highlighted in blue, in samples with low TMD in green and across both groups (high/low TMD) in red.

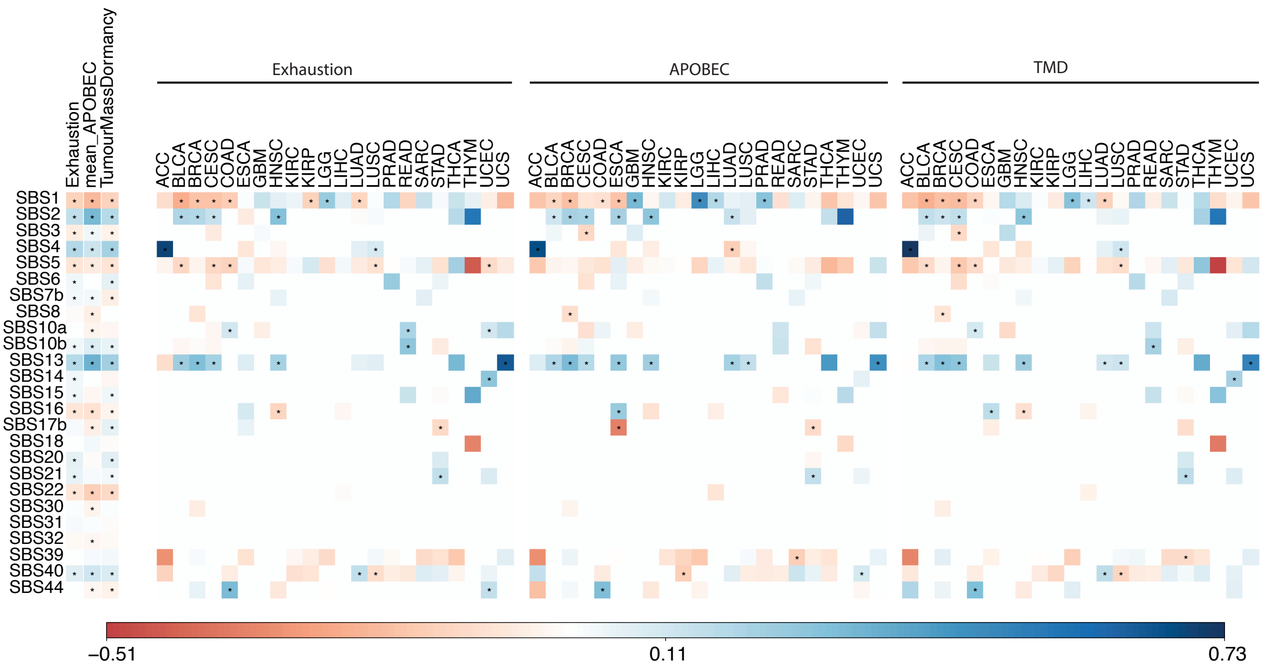


**Supplementary Figure 44. Mutational processes linked with PCA-derived TMD levels.** Matrices depicting the Person correlation between mutational signatures and the TMD, exhaustion and APOBEC programmes, both pan-cancer (first 3 columns) and within individual cancer tissues are shown for the case where TMD is calculated using the PCA-based method. Statistically significant correlations (p<0.05) are highlighted with an asterisk. Only signatures with at least one significant correlation are shown.

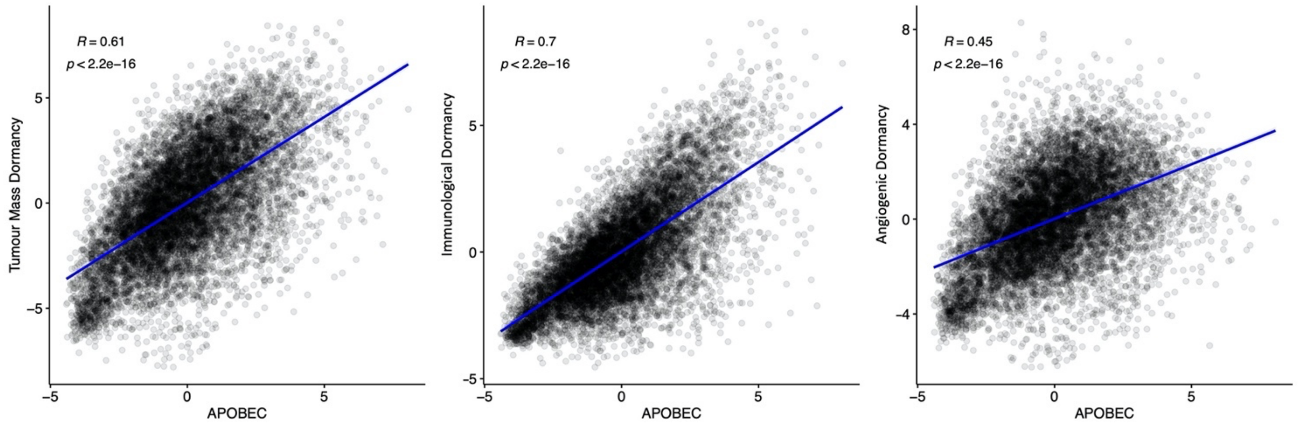


**Supplementary Figure 45. Correlation between PCA-derived dormancy programmes and APOBEC activity.** The x and y axes depict the per-sample expression score for the respective programmes.

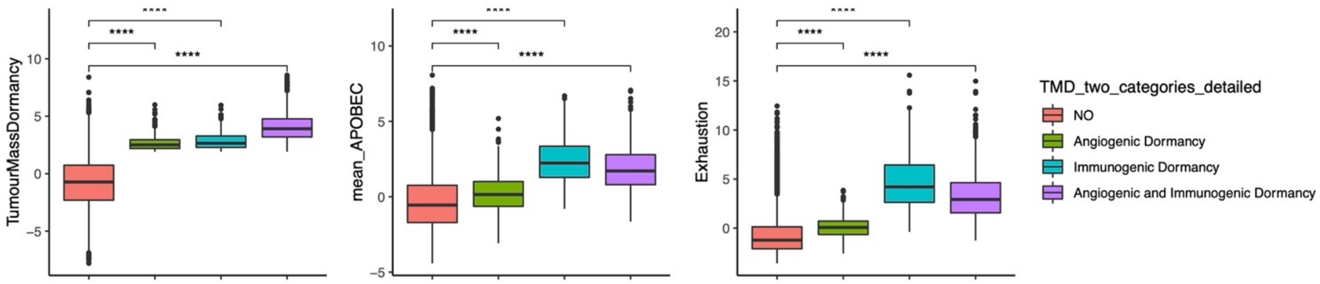


**Supplementary Figure 46. PCA-derived dormancy categories and linked progammes.** TMD, APOBEC and exhaustion programme scores compared between samples with angiogenic dormancy, immunological dormancy, both angiogenic and immunological dormancy and expanding tumours without evidence of TMD (NO); **** p<0.0001.

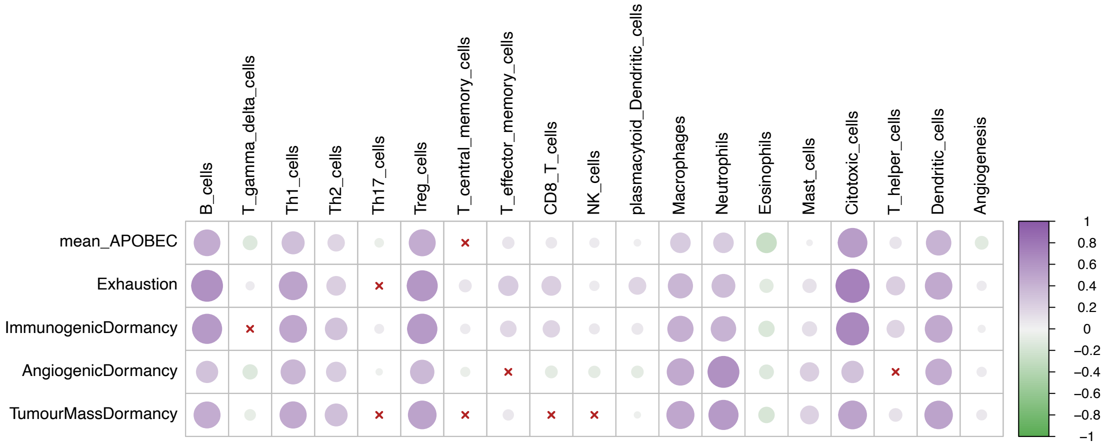


**Supplementary Figure 47. Tumour microenvironment activity correlates with PCA-derived TMD levels.** Correlation between cell infiltration estimates and the APOBEC, exhaustion and PCA-derived dormancy programme scores across the TCGA primary tumor samples. The colour gradient reflects the Pearson correlation coefficient.

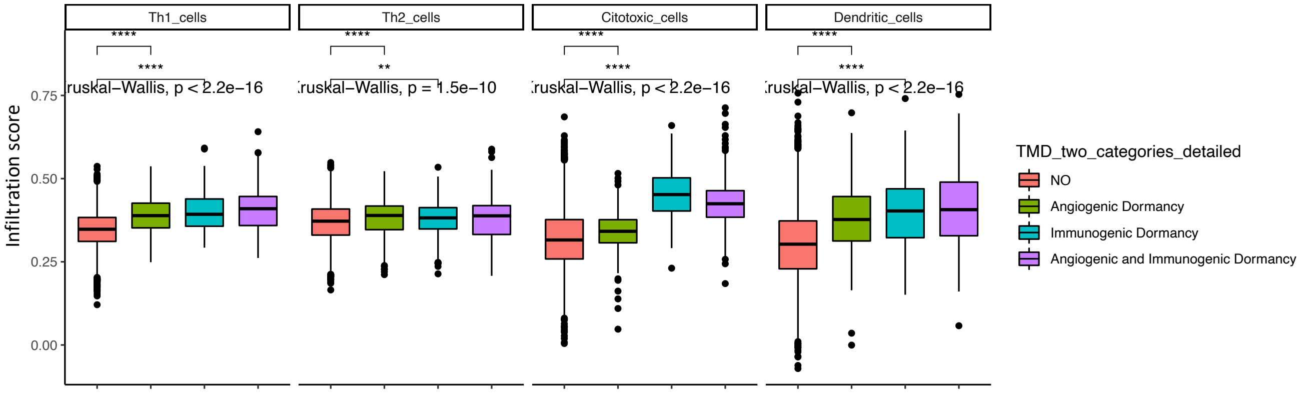


**Supplementary Figure 48. PCA-derived TMD categories differ in immune cell composition.** Cell infiltration scores are compared between samples with angiogenic dormancy, immunological dormancy, both angiogenic and immunological dormancy and expanding tumours without evidence of TMD (NO); **** p<0.0001.

**Supplementary Table 1. Markers of immunologic/angiogenic dormancy and exhaustion.**

| Gene | Direction | Type | Annotation |
|---|---|---|---|
| CD8A | up | immunologic | Classical markers of CD8+ T cells. CD8+ T cells are key in the immunological dormancy process through the elimination of cancer cells that limits tumour growth [1-3]. |
| CD8B | up | immunologic | |
| CD4 | up | immunologic | Classical marker of CD4+ T cells. These cells would act in conjunction with the CD8+ T cell to generate antitumour responses specific of the TMD stage of cancer development [2]. An increased ratio of CD8/CD4 T cells is indicative of immunological dormancy [3]. |
| IFNG | up | immunologic | Interferons are important activators of the anti-tumour response. However, accumulation of interferon gamma in the tumour microenvironment leads to immunosuppression and upregulation of PD1 ligand on CD8+ T cells [4]. Moreover, it was shown that longstanding interferon gamma signalling in tumour cells leads to acquisition of epigenetic modifications leading to the expression of more interferon-stimulated genes as well as the expression ligands for multiple T cell inhibitory receptors [5]. |
| STAT1 | up | immunologic | STAT1 mediates the activation of cell cycle inhibitors, p21 WAF1/CIP1 and p27 KIP1, which results in a reduction of cell proliferation [6]. |
| IL12A | up | immunologic | IL-12 is critical for IFN-gamma production and its inhibition was shown to induce tumour growth in sarcoma [7]. |
| ITGB1 | down | immunologic | Integrins drive a cascade of Ras/ERK activation which sustains cell proliferation. Downregulation of integrins has been shown to induce dormancy [8, 9]. |
| ITGA5 | down | immunologic | |
| EGFR | down | immunologic | EGFR is upregulated as a result of uPAR activation, which regulates tumour growth. Blocking this pathway induces growth arrest and dormancy [10]. |
| PTK2 | down | immunologic | FAK (PTK2) acts downstream of uPAR to activate mitogen signalling and cell proliferation. Blocking it has been shown to induce dormancy [11]. |
| ERBB2 | down | immunologic | uPAR also triggers mitogenic signalling of ERBB2, inducing tumour growth. Blocking ERBB2 has been stipulated to be a likely target switch for manipulating tumour dormancy [2]. |
| GAS6 | up | immunologic | GAS6 and BMP7 have been shown to induce dormancy in bone cancers by activating TAM kinase signalling and p38, respectively [12]. |
| BMP7 | up | immunologic | |
| DKK1 | up | immunologic | Inhibition of Wnt signalling by DKK1 has been shown to induce quiescence [13]. |
| IFNB1 | up | immunologic | Type I interferons, such as interferon beta, are known regulators of cancer immunity, and immune response in general. They bind to interferon receptor (IFNAR) and activate the transcription of many genes through the activation of intracellular transcription factors, such as IRF7. It was shown that chemotherapy induces a type I IFN response in tumour cells as a result of MDSC signalling. This results in a self-sustained overexpression of IRF7 and consequently, IFNβ, which triggers the dormancy programme [14]. |
| IDO1 | up | immunologic | Interferon-gamma mediates dormancy in tumour-repopulating cells via the IDO1/p27 axis [3]. |
| TGFB2 | up | immunologic | TGF-beta 2 was found to decrease the ERK/p38 signalling ratio and induce cells into dormancy through DEC2 expression [3]. |
| DIRAS3 | up | immunologic | ARHI (DIRAS3) expression leads to cell death and dormancy through inhibition of PI3K signalling [15]. |

| | | | |
|---|---|---|---|
| VEGFA | down | angiogenic | VEGFA, VEGFB and KDR are major regulators of angiogenesis [16]. |
| VEGFB | down | angiogenic | |
| KDR | down | angiogenic | |
| CXCL8 | down | angiogenic | CXCL8 stimulates VEGF expression and vascularization [17]. |
| CXCR1 | down | angiogenic | Molecules expressed on endothelial cells. Their knockdown was shown to inhibit endothelial cell proliferation [18]. |
| CXCR2 | down | angiogenic | |
| THBS1 | up | angiogenic | Thrombospondin 1 is a potent angiogenesis inhibitor [19]. |
| SPP1 | down | angiogenic | Osteopontin has been shown to induce angiogenesis through PI3K/AKT and ERK activation [20]. |
| CXCL9 | up | angiogenic | Antiangiogenic chemokines released by immune cells have been reported to act as a bridge between immune dormancy and angiogenic dormancy [107]. CD4+ T cells were shown to release CXCL9 and CXCL10, which inhibit vascularization processes in the growing tumour, indirectly contributing to the induction of cancer dormancy [5]. |
| CXCL10 | up | angiogenic | |
| ITGAV | down | angiogenic | ITGAV has been shown to promote angiogenesis and cancer progression [21]. |
| COL18A1 | up | angiogenic | Endostatin is a potent inhibitor of angiogenesis [22]. |
| ADGRB1 | up | angiogenic | Angiogenesis inhibitor whose expression inversely correlates with metastatic spread [23]. |
| PLG | down | angiogenic | Plasminogen degrades extracellular matrix proteins thereby facilitating cell migration, and has been shown to play a role in angiogenesis [24]. |
| PLAUR | down | angiogenic | The urokinase plasminogen activator surface receptor (uPAR) modulates VEGF-induced angiogenesis [25]. |
| SERPINE1 | up | angiogenic | SERPINE1 inhibits uPA, thereby inhibiting angiogenesis [26]. |
| HIF1A | up | angiogenic | HIF1A is a master transcriptional regulator of hypoxia [27]. |
| CTLA4 | up | exhaustion | CTLA4 and PD-1 (PDCD1) are well characterised inhibitory receptors expressed on the surface of the immune cells. Antibodies directed against those ligands are currently used in the clinic [28]. |
| PDCD1 | up | exhaustion | |
| TIGIT | up | exhaustion | TIGIT is an immunoglobulin superfamily member shown to be expressed on CD8+ tumour infiltrating T cells and natural killer cells [29]. In human cancers, TIGIT acts in conjunction with PD-1 to inhibit effector T cells, hence enabling progression of cancer . Tigit–/– mice do not exhibit NK cell exhaustion and show fewer metastases and improved survival, thereby further highlighting the key role of TIGIT in regulating cancer immunosurveillance [30]. |
| LAG3 | up | exhaustion | LAG3 is one of the known inhibitory receptors (IRs) expressed on the surface of the T cells. The exact signalling mechanisms downstream of LAG3 and interplay with other IRs remains unknown due to LAG3's structure which is unique and distinct from other IRs [31]. However, in vivo blockade of the PD1 and LAG3 IRs together led to a greater reversal of T cell exhaustion and viral control compared to blockade of either one of those pathways alone [31]. |
| HAVCR2 | up | exhaustion | TIM3 (HAVCR2) is expressed on CD8+ T cells in the tumour microenvironment in mouse models of solid tumours. They co-express PD1 and exhibit a severe exhausted phenotype [32]. Very recently, TIM3 was linked to myeloid derived suppressor cells, which trigger the TIM3+ CD8+ T cells through a Gal9 receptor in human studies, causing T cell exhaustion [33, 34]. |

| | | | |
|---|---|---|---|
| EOMES | up | exhaustion | EOMES and TBET are paralogous transcription factors and master regulators of cytotoxic T cell lineage commitment and exhaustion programme activation [35, 36]. EOMES is required for induction of effector CD8+ T cells by IFNγ induction in CD8+ T cells [35] and is upregulated, along with PD1, in chronic viral infections in vivo [37]. This points to its role in T cell exhaustion. TBET, similarly to BLIMP1, promotes differentiation of cytotoxic T cells at the early stages of infection [38]. Persistent antigenic stimulation causes downregulation of TBET, which competes for genomic binding sites with EOMES and causes an exhaustion phenotype [35, 38]. |
| TBX21 | up | exhaustion | |
| BTLA | up | exhaustion | BTLA is a well-known exhaustion marker in cancer [39]. |
| CD274 | up | exhaustion | PD-L1 (CD274) is one of the central regulators of T cell exhaustion [40]. |
| PTGER4 | up | exhaustion | Implicated in T cell exhaustion in metastases of melanoma patients [41]. |
| CD244 | up | exhaustion | Demonstrated to have an inhibitory role leading to exhaustion of CD8+ T cells and NK cells [42]. |
| CD160 | up | exhaustion | CD160 binds to MHC class I molecules and delivers a co-stimulatory signal necessary for CD8+ T cell activation [43]. However, prolonged CD160 signalling seems to lead to inhibition of TCR signalling and, consequently, T cell exhaustion. CD8+ T-cell populations expressing CD160 have reduced proliferation capacity and perforin expression in vitro. Conversely, the blockade of CD160 interaction with its ligand restores the proliferation of CD8+ T cells [44]. |

### Supplementary Table 1 references:

**Supplementary Table 2. Prevalence of TMD, angiogenic and immunological dormancy in different tissues and pan-cancer.**

| Cancer type | TMD frequency | AD frequency | ID frequency | AD+ID frequency |
|---|---|---|---|---|
| Pan-cancer | 16.5% | 4.4% | 5% | 7% |
| KICH | 0 | 0 | 0 | 0 |
| READ | 0.006 | 0 | 0.006 | 0 |
| GBM | 0.019 | 0 | 0.013 | 0.006 |
| KIRP | 0.024 | 0.003 | 0.010 | 0.010 |
| LGG | 0.027 | 0.002 | 0.023 | 0.002 |
| COAD | 0.035 | 0.009 | 0.020 | 0.007 |
| UCS | 0.036 | 0 | 0 | 0.036 |
| ACC | 0.063 | 0.025 | 0.013 | 0.025 |
| LIHC | 0.067 | 0.032 | 0.013 | 0.022 |
| CHOL | 0.083 | 0 | 0.056 | 0.028 |
| TGCT | 0.087 | 0.020 | 0.013 | 0.053 |
| UVM | 0.088 | 0.013 | 0.013 | 0.063 |
| CESC | 0.089 | 0.010 | 0.036 | 0.043 |
| KIRC | 0.100 | 0.008 | 0.066 | 0.026 |
| PRAD | 0.137 | 0.107 | 0.002 | 0.028 |
| ESCA | 0.143 | 0.075 | 0.012 | 0.056 |
| BLCA | 0.159 | 0.047 | 0.015 | 0.098 |
| PAAD | 0.169 | 0.011 | 0.107 | 0.051 |
| UCEC | 0.173 | 0.009 | 0.099 | 0.064 |
| LUSC | 0.178 | 0.014 | 0.088 | 0.076 |
| SKCM | 0.194 | 0.029 | 0.049 | 0.117 |
| THCA | 0.209 | 0.006 | 0.151 | 0.052 |
| STAD | 0.216 | 0.064 | 0.027 | 0.125 |
| OV | 0.217 | 0.024 | 0.083 | 0.110 |
| LUAD | 0.220 | 0.025 | 0.113 | 0.082 |
| MESO | 0.233 | 0.035 | 0.070 | 0.128 |
| BRCA | 0.257 | 0.138 | 0.009 | 0.109 |
| SARC | 0.316 | 0.073 | 0.077 | 0.166 |
| HNSC | 0.322 | 0.134 | 0.034 | 0.154 |
| THYM | 0.327 | 0 | 0.042 | 0.29 |
| PCPG | 0.331 | 0.039 | 0.213 | 0.079 |

**Supplementary Table 3. Tumour drivers with enrichment or depletion of SNVs within primary tumour samples exhibiting tumour mass dormancy.**

| Gene | Odds ratio | Adjusted p-value | Annotation |
|---|---|---|---|
| **APC** | 0.24 | 2.56e-12 | -Suppresses tumour growth though repression of the Wnt signalling pathway [1]<br>-Promotes apoptosis by downregulating the expression of survivin [2] |
| **TP53** | 0.52 | 1.09e-08 | -Transcription factor which coordinates signals of stresses caused by a variety of stresses including DNA damage and aberrant growth signalling and induces cell cycle arrest, apoptosis, senescence, DNA repair and/or changes in metabolism [3][4]<br>-Inhibits angiogenesis by upregulating maspin expression [5]<br>-Stimulates the innate immune system to suppress tumorigenesis by inducing senescence and SASP [6] |
| **KRAS** | 0.32 | 1.48e-08 | -Activating mutations of the small GTPase cause hypersensitivity to external growth-simulating factors resulting in increased signalling though PI3K, MEK/ERK and other pathways regulating proliferation [7] |
| **FAT4** | 0.52 | 4.43e-03 | -Cadherin protein which inhibits cell proliferation by suppressing the phosphorylation and nuclear accumulation of Yap member of the Hippo signalling pathway [8] |
| **CASP8** | 4.93 | 4.60e-03 | -A cysteine-aspartic acid protease which plays a central role in the execution of apoptosis by cleaving and thereby activating caspase 3 and caspase 7 [9]<br>-Following NF-kB activation it promotes angiogenesis by enhancing VEGF expression<br>-CASP8 inactivation allows cancer cells to evade apoptosis initiated by T and NK immune cells [10] |
| **KMT2D** | 0.50 | 6.58e-03 | -A histone-lysine N-methyltransferase whose KO results in cell cycle arrest in oesphageal squamous cell carcinoma cell lines [11] |
| **DCC** | 0.44 | 6.94e-03 | -Encodes the Nectin-1 receptor, which when bound by nectin-1 triggers caspase-9 dependent apoptosis. Production of nectin-1 at the base of villi within the gastrointestinal tract inhibits apoptosis of the epithelial cells until they reach the tip of the villus [12] |
| **HRAS** | 5.47 | 6.94e-03 | -Activating mutations of the small GTPase cause hypersensitivity to external growth-simulating factors resulting in increased signalling though PI3K, MEK/ERK and other pathways regulating proliferation [7] |
| **KDM6A** | 0.35 | 1.92e-02 | --An oxygen-sensitive histone methylase which functions to demethylase H3K27 residues, under normoxia conditions [13]<br>-Slow-cycling glioma cells are dependent on KDM6A epigenetic regulation [14]<br>-KDM6A activates Th1-type chemokines needed for T cell migration in mouse and human medulloblastoma models [13] |
| **IDH1** | 0.32 | 2.20e-02 | -Encodes isocitrate dehydrogenase 1, an enzyme which catalyses the oxidative decarboxylation of isocitrate to α-ketoglutarate within the Krebs cycle. |

| | | | |
|---|---|---|---|
| | | | -Mutations in codon 132 result production of 2-hydroxyglutarate which results in changes of specific histone marks and extensive DNA hypermethylation, which result in the downregulation of leukocyte chemotaxis factors [15] [16]<br>-Angiogenesis is inhibited by 2-hydroxyglutarate, which by regulating the activity of α-ketoglutarate dependent dioxygenases causes the ubiquitination and proteasomal degradation of HIF1A [17] |
| **PRDM16** | 0.30 | 2.20e-02 | -A transcriptional coactivator and corepressor which is associated with evasion of apoptosis in prostate cancer [18], [19]<br>-In renal cancer it suppresses the expression of a HIF target semaphorin 5B, which normally promotes tumour growth [20] |
| **FGFR3** | 0.29 | 2.45e-02 | -A tyrosine-protein kinase receptor for fibroblast growth factors which plays an essential role in regulating cell proliferation and apoptosis.<br>-Ectopic expression of FGFR3 results in increased cell proliferation and lower levels of apoptosis in myeloma [21] |
| **NCOA1** | 0.27 | 3.00e-02 | -A transcriptional coactivator for steroid and nuclear hormone receptors, which upregulates VEGF α in breast cancer tumours [22] |
| **FBXW7** | 0.50 | 3.05e-02 | -A ubiquitin ligase which regulates quiescence by mediating ubiquitin-dependent proteolysis of key cell cycle proteins including cyclin E1 and c-Myc [23] |
| **NRAS** | 0.28 | 3.40e-02 | -Activating mutations of the small GTPase result in hypersensitivity to external growth-simulating factors and increased signalling though PI3K, MEK/ERK and other pathways regulating proliferation [7] |

**Supplementary Table 3 references:**


[1] B. M. Boman and J. Z. Fields, "An APC: WNT counter-current-like mechanism regulates cell division along the human colonic crypt axis: A mechanism that explains how APC mutations induce proliferative abnormalities that drive colon cancer development," *Front. Oncol.*, 2013.

[2] T. Zhang *et al.*, "Evidence that APC regulates survivin expression: A possible mechanism contributing to the stem cell origin of colon cancer," *Cancer Res.*, 2001.

[3] B. Vogelstein, D. Lane, and A. J. Levine, "Surfing the p53 network," *Nature*, 2000.

[4] K. Polyak, Y. Xia, J. L. Zweier, K. W. Kinzler, and B. Vogelstein, "A model for p53-induced apoptosis," *Nature*, 1997.

[5] M. J. C. Hendrix, "De-mystifying the mechanism(s) of maspin," *Nature Medicine*. 2000.

[6] N. Raj and L. D. Attardi, "Tumor suppression: P53 alters immune surveillance to restrain liver cancer," *Current Biology*. 2013.

[7] S. Schubbert, K. Shannon, and G. Bollag, "Hyperactive Ras in developmental disorders and cancer," *Nature Reviews Cancer*. 2007.

[8] L. Ma, J. Cui, H. Xi, S. Bian, B. Wei, and L. Chen, "Fat4 suppression induces Yap translocation accounting for the promoted proliferation and migration of gastric cancer cells," *Cancer Biol. Ther.*, 2016.

[9] B. Tummers and D. R. Green, "Caspase-8: regulating life and death," *Immunological Reviews*.

2017.

[10] A. Sistigu, M. Musella, C. Galassi, I. Vitale, and R. De Maria, "Tuning Cancer Fate: Tumor Microenvironment's Role in Cancer Stem Cell Quiescence and Reawakening," *Frontiers in Immunology*. 2020.

[11] A. Abudureheman *et al.*, "High MLL2 expression predicts poor prognosis and promotes tumor progression by inducing EMT in esophageal squamous cell carcinoma," *J. Cancer Res. Clin. Oncol.*, 2018.

[12] C. Forcet *et al.*, "The dependence receptor DCC (deleted in colorectal cancer) defines an alternative mechanism for caspase activation," *Proc. Natl. Acad. Sci. U. S. A.*, 2001.

[13] J. Yi, X. Shi, Z. Xuan, and J. Wu, "Histone demethylase UTX/KDM6A enhances tumor immune cell recruitment, promotes differentiation and suppresses medulloblastoma," *Cancer Lett.*, 2021.

[14] B. B. Liau *et al.*, "Adaptive Chromatin Remodeling Drives Glioblastoma Stem Cell Plasticity and Drug Tolerance," *Cell Stem Cell*, 2017.

[15] S. Turcan *et al.*, "IDH1 mutation is sufficient to establish the glioma hypermethylator phenotype," *Nature*, 2012.

[16] S. Tommasini-Ghelfi, K. Murnan, F. M. Kouri, A. S. Mahajan, J. L. May, and A. H. Stegh, "Cancer-associated mutation and beyond: The emerging biology of isocitrate dehydrogenases in human disease," *Sci. Adv.*, 2019.

[17] D. Ye, S. Ma, Y. Xiong, and K. L. Guan, "R-2-hydroxyglutarate as the key effector of IDH mutations promoting oncogenesis," *Cancer Cell*. 2013.

[18] S. Zhu *et al.*, "PRDM16 is associated with evasion of apoptosis by prostatic cancer cells according to RNA interference screening," *Mol. Med. Rep.*, 2016.

[19] J. Chi and P. Cohen, "The multifaceted roles of PRDM16: Adipose biology and beyond," *Trends in Endocrinology and Metabolism*. 2016.

[20] A. Kundu *et al.*, "PRDM16 suppresses HIF-targeted gene expression in kidney cancer," *J. Exp. Med.*, 2020.

[21] E. E. Plowright *et al.*, "Ectopic expression of fibroblast growth factor receptor 3 promotes myeloma cell proliferation and prevents apoptosis," *Blood*, 2000.

[22] L. Qin *et al.*, "NCOA1 promotes angiogenesis in breast tumors by simultaneously enhancing both HIF1a- and AP-1-mediated VEGFa transcription," *Oncotarget*, 2015.

[23] A. C. Minella and B. E. Clurman, "Mechanisms of tumor suppression by the SCFFbw7," *Cell Cycle*. 2005.
